# Spatial and single-cell transcriptomics illuminate bat immunity and barrier tissue evolution

**DOI:** 10.1101/2023.10.30.564705

**Authors:** Roy Levinger, Dafna Tussia-Cohen, Sivan Friedman, Yan Lender, Yomiran Nissan, Evgeny Fraimovitch, Yuval Gavriel, Jacqueline Tearle, Aleksandra A. Kolodziejczyk, Tomás Gomes, Natalia Kunowska, Maya Weinberg, Giacomo Donati, Kylie R James, Yossi Yovel, Tzachi Hagai

**Author notes:** These authors contributed equally (Roy Levinger, Dafna Tussia-Cohen).

## Abstract

The Egyptian fruit bat displays tolerance to lethal viruses and unique dietary adaptations, but the molecular basis for this is poorly understood. To this end, we generated detailed maps of bat gut, lung and blood cells using spatial and single-cell transcriptomics. We compared bat with mouse and human cells to reveal divergence in genetic programs associated with environmental interactions and immune responses. Complement system genes are transcriptionally divergent, uniquely expressed in bat lung and gut epithelium, and undergo rapid coding-sequence evolution. Specifically in the tip of the gut villus, bat enterocytes express evolutionarily young genes while lacking expression of genes related to specific nutrient absorption. Profiling immune stimulation of PBMCs revealed a monocyte subset with conserved cross-species interferon expression, suggesting strong constraints to avoid an excessive immune response. Our study thus uncovers conserved and divergent immune pathways in bat tissues, providing a unique resource to study bat immunity and evolution.

## Introduction

Accounting for ∼20% of all mammals and distributed across the globe, bats constitute a large and ecologically diverse clade that evolved several unique adaptations, including powered flight, echolocation and extreme longevity. Several bat species display mild or no clinical symptoms when infected by some of the viruses deadliest to humans, including Henipaviruses^1^, Filoviruses^2^ and Coronaviruses^3,4^. The majority of these cases involve members from two bat families – Old World fruit bats (Megabats, *Pteropodidae*) and horseshoe bats (*Rhinolophidae*). Bats are suspected to be the natural reservoir for several of these viruses, and the potential source of recent zoonotic events, including the zoonotic transfers of Betacoronaviruses, SARS-CoV-1, SARS-CoV-2 and MERS, to humans^3,5–7^. The mechanisms of bat adaptation to these viruses have been a continuous source of interest, leading to numerous studies that focused on the immune systems of different bat species^8^. These studies found evolutionary changes in specific genes (such as in recent duplication and rapid coding sequence evolution of antiviral restriction factors^9,10^) and in entire pathways (such as in constitutive expression of type-I interferons^11^ (**IFNs**), and in dampening the inflammasome-mediated response through diverse mechanisms^8,12^).

The Egyptian fruit bat (*Rousettus aegyptiacus*) is a megabat that is thought to serve as the natural reservoir of the highly pathogenic Marburgvirus^2^. In addition, *R. aegyptiacus* infection with Ebola virus is asymptomatic, despite successful viral replication in several bat tissues^13–15^. As such, *R. aegyptiacus* is considered an important model for studying the mechanisms of immune tolerance to viral infection, and was proposed as one of the key bat species for comprehensive immune characterization^16^. Consequently, *R. aegyptiacus* was the focus of several works studying different aspects of its immune system and viral infection^13,15,17–23^.

*R. aegyptiacus*, like other Old World fruit bats, possess unique adaptations to flight and to a high-sugar and low-fat fruit-based diet. Flight adaptation in these bats is associated with relatively large lungs, major structural changes in the intestine to reduce energetic costs associated with carrying food during flight^24^, and a fast metabolism to meet high caloric demands^25^. Metabolic adaptation of *R. aegyptiacus* to its diet is poorly understood at the molecular and cellular levels. Evolutionary analyses of the two bat lineages that have independently evolved obligate frugivory found evidence for adaptation to dietary changes through expansion and contraction of specific gene families and in amino acid substitutions in metabolic genes^26–28^. More recently, a comparative analysis of insectivorous and frugivorous microbats, found divergence in expression of genes related to fluid and electrolyte balance in the kidney and in insulin secretion in the pancreas^29^. These studies suggest that transcriptomics analysis of *R. aegyptiacus* cells from relevant tissues can deepen our understanding of the molecular mechanisms that underly bat unique metabolism and diet adaptation.

Here, we present a spatially resolved single-cell transcriptomics map of *R.aegyptiacus* cells from tissues relevant for immune and metabolic adaptations. We created a comprehensive atlas of *R. aegyptiacus* single-cell transcriptomes from lung and gut tissues, and of peripheral blood mononuclear cells (**PBMCs**) (**Fig 1a**). We also profiled an equivalent set of mouse cells in a comparative manner and used publicly available single-cell data from human tissues to find conserved and divergent pathways in *R. aegyptiacus* tissues. In the gut, we combined spatial with single-cell transcriptomics, allowing us to study the links between genetic programs and their expression in specific gut regions. This revealed the unique expression of complement system genes in the bat intestinal crypt and enabled profiling of enterocyte differentiation during their migration along the villus. Furthermore, we compared mouse and bat PBMCs after triggering antiviral and antibacterial responses. This allowed us to characterize the intrinsic innate immune response across cell types and species, finding conserved and divergent pathways. Overall, this comprehensive set of nearly a quarter of a million cells across organs provided the basis for comparing genetic programs between mouse, human and *R. aegyptiacus*, and for analyzing important environmental and pathogen interaction pathways in this unique species.

**Figure 1:**
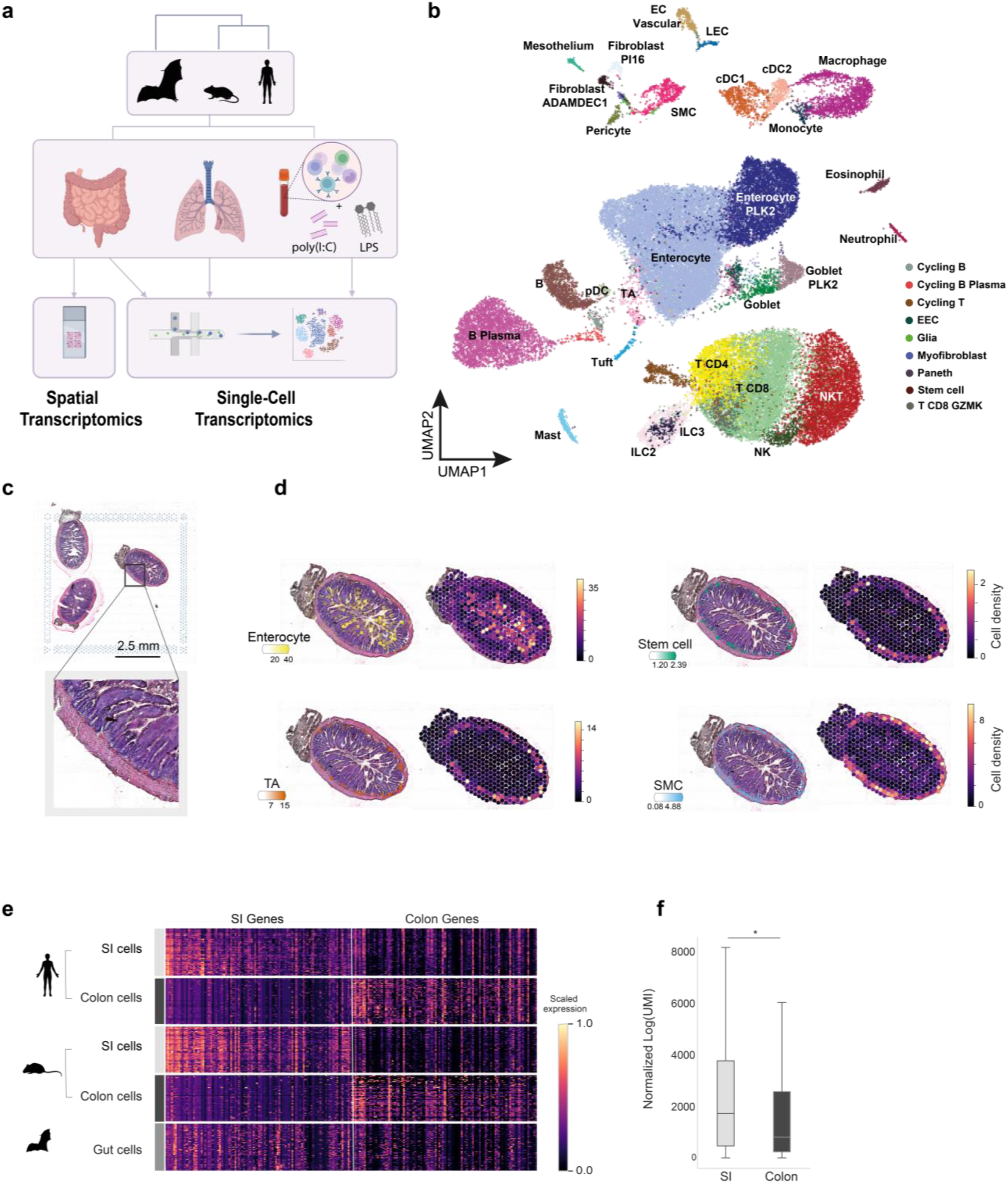
Spatial and transcriptional characterization of *R. aegyptiacus* gut cells. (**a**) System overview: Gut– and lung-resident cells from *R. aegyptiacus* and mouse were profiled using single-cell transcriptomics. PBMCs were profiled in steady-state and following the triggering of the antiviral and antibacterial innate immune responses using dsRNA (poly I:C) and lipopolysaccharide (LPS), respectively. *R. aegyptiacus* gut samples were also characterized using spatial transcriptomics analysis. *R. aegyptiacus* and mouse cells along with publicly available human cells in each of the respective tissues, were compared using one-to-one orthologous genes and following gene orthology assignments using EggNOG^31^. **(b)** Uniform manifold approximation and projection (UMAP) of *R. aegyptiacus* intestinal cell clusters (8 samples, n= 46,316 cells). **(c)** Hematoxylin and eosin (H&E) staining of transverse cross-section of *R. aegyptiacus* gut (n=3 samples). **(d)** Spatial mapping of scRNA-seq data to 10x Genomics Visium data showing estimated abundance (colour intensity) of cell subsets (colour) of a cross-section. **(e)** Heatmaps showing gene expression of SI– and colon-specific genes in human and mouse SI– and colon-resident enterocytes and in *R. aegyptiacus* enterocytes. **(f)** Distribution of SI– and colon-specific gene expression in *R. aegyptiacus* enterocytes compared using a one-sided Mann-Whitney test (*=P<0.05).

## Results

### An integrated spatial and transcriptional map of *R. aegyptiacus* intestinal cells

To investigate cell composition and gene expression in the *R. aegyptiacus* intestinal tract and how these may be related to structural and functional changes associated with flight, innate immunity and fast sugar metabolism adaptations, we performed single-cell RNA sequencing (**scRNA-seq**) on samples from the upper gut section of 8 adult males (roughly equivalent to the jejuno-ileal region, see **Methods** for details). For comparison, we also performed an analogous procedure on small intestine (**SI**) and colon samples from mice. With both *R. aegyptiacus* and mouse tissues, we employed tissue dissociation procedures similar to a previously published protocol used with human samples^30^, to minimize technical differences between samples from the three species. In addition, we analyzed cell spatial localization in *R. aegyptiacus* gut by performing spatial transcriptomics of 11 gut sections from two *R. aegyptiacus* individuals. To annotate *R. aegyptiacus* cell clusters and compare them to analogous cell populations in human and mouse, we first employed EggNOG^31^ to identify orthology relationship in a genome-wide manner between *R. aegyptiacus,* mouse and human coding genes, followed by manual inspection and further annotations (see **Methods** and **Table S1**). We then used computationally derived and experimentally established markers of various cell types expected to be found in our tissues, to associate *R. aegyptiacus* cell clusters with their likely cell lineages.

Following QC, the *R. aegyptiacus* single-cell data included more than 46,000 high-quality gut cells (**Fig 1b**). Using clustering, orthologous marker-gene analysis and transcriptome correlation with human and mouse data, we identified 38 cell subsets from the immune, epithelial, mesenchymal and endothelial lineages (**Fig 1b** and **Fig S1**, and see corresponding analysis on mouse, where we obtained over 30,000 high-quality cells, in **Fig S2-3)**. To further validate our annotations, we compared gene expression of different cell clusters between species (**Fig S4-5**). In the epithelial compartment, which will be the focus of most of our subsequent analysis, we identified absorptive cells, enterocytes and precursors (stem cells and transit amplifying (**TA**) cells). Secretory cells included goblet, tuft and Paneth cells as well as enteroendocrine cells (**EECs**). Finally, spatial transcriptomics analysis using cell2location^32^ identified stem and TA cells at the crypt and enterocytes across the villus in the mucosa. In addition, smooth muscle cells (**SMCs**) were found in the most peripheral layer of the section, enabling distinctions between the different layers (**Fig 1c-d**). Thus, this cell localization analysis of bat gut sections identified the spatial location of diverse cell populations and showed conserved overall architecture in the transverse cross-section, similar to human and mouse gut.

### Transcriptional comparison of bat enterocytes with small intestine enterocytes and colonocytes from mouse and human

*R. aegyptiacus* gut morphology has not been comprehensively characterized at the molecular and cellular levels. Similarly to other megabats^25^, there are no clear macro-morphological differences between various gut regions^24^. Our histological sections showed structural similarity to small intestine (**SI**) architecture, with long villi (longer than in rodents) and crypts (see **Fig 1c**). To better understand to which of the two main gut regions, SI and colon, bat cells are more transcriptionally similar to, we compared the *R. aegyptiacus* scRNA-seq data to our SI and colon mouse data and to previously published SI and colon human data^30^. Enterocytes residing in the SI and the colon (i.e., colonocytes) are thought to display the greatest transcriptional differences between the two gut regions, in comparison with inter-region differences displayed by other cell types^30,33^. We thus focused on enterocytes, and performed a differential expression (**DE**) analysis between mouse SI enterocytes and colonocytes, and, separately, between these two cell groups in human. Using SI versus colon DE analyses from both mouse and human, we found the top 100 SI– and colon-specific orthologous genes, and compared their relative expression levels in human and mouse SI and colon as well as in *R. aegyptiacus* gut enterocytes (**Fig 1e**). We observed that *R. aegyptiacus* enterocytes express SI-specific genes in higher levels than colon-specific genes (P-value = 0.03, Mann-Whitney test, **Fig 1f**). We note however, that some colon-specific genes are still expressed in bat enterocytes, although significantly lower than SI-specific genes. This analysis suggests that *R. aegyptiacus* gut cells, at least from the region we sampled, are more transcriptionally similar to SI cells. Therefore, in the following analyses we compare them to human and mouse SI enterocytes.

### Enterocyte expression dynamics along the villus is largely conserved

The intestinal epithelium is a well-structured tissue, responsible for nutrient absorption, hormone and mucus secretion, interactions with commensal microbiota and defense against pathogens^34,35^. Enterocytes, the most abundant cells in the epithelial layer, emerge in the crypt and migrate along the villus, until being shed off from the top of the villus a few days later^36^. Enterocytes were shown to traverse a series of cell states, identified by distinct gene expression and related to different functions, during their migration from the crypt along the villus axis^37^ (**Fig 2a**). As these functions are related to innate immunity and nutrient absorption, we asked whether these region-dependent transcriptional programs originally found in mouse SI, are conserved in the bat villus. For this, we used genes previously found to be highly and specifically expressed in each of the 5 consecutive regions located from the bottom to the top of the villus in the mouse (denoted as regions 1 to 5, **Fig 2a**). Using the *R. aegyptiacus* spatial transcriptomics data, we observed a clear trend where the sets of the bat orthologs of the zonation marker genes are expressed from proximal to distal regions of the section (**Fig 2b**, see **Fig S6-7a-b**). This observation suggests a largely conserved gene expression dynamics along the villus in mouse and bat.

**Figure 2:**
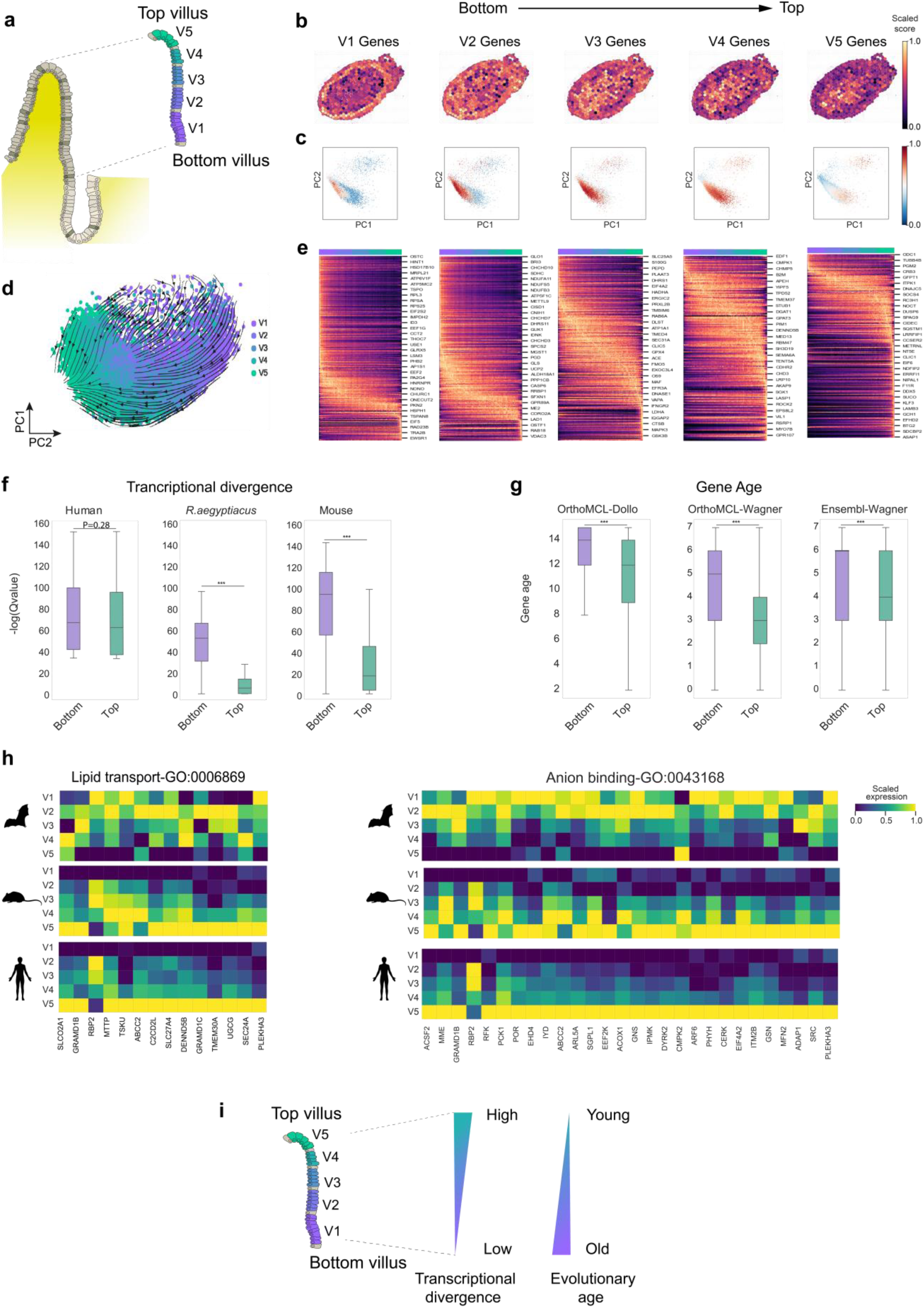
Evolutionary analysis of enterocyte differentiation along the intestinal villus. (**a**) Schematic of a transverse cross-section of the intestinal crypt and villus. The villus is partitioned into 5 consecutive bottom-to-top regions (denotes as v1-v5). **(b)** Scores of landmark gene expression (using mouse-to-*R. aegyptiacus* orthologs) from each of the bottom-top regions (v1-v5) overlayed on a 10x Genomics Visium data of a *R. aegyptiacus* gut cross-section (additional samples are found in Fig S6). **(c)** Principal component analysis (PCAs) of *R. aegyptiacus* enterocytes (based on single-cell data) overlayed with scores of bottom-to-top regions (v1-v5) (red and blue mark high and low expression, respectively). **(d)** UMAP of *R. aegyptiacus* enterocytes colored by their inferred spatial localization along the tissues (v1-v5) with dashed lines and arrows depicting inferred transcriptional trajectories as determined by scVelo. **(e)** Heatmaps showing landmark gene expression in *R. aegyptiacus* enterocytes, each panel including a different set of regional genes (v1-v5 regions, genes are based on mouse data^37^). Cells are ordered based on PC1 axis from 2c that denotes bottom-to-top villus localization. **(f)** Transcriptional conservation of top and bottom villus genes: using a matched (by Q-value) set of top and bottom human genes (left), we look at the DE values (in Q-values) of these sets in mouse and *R. aegyptiacus* enterocytes, using one-to-one orthologs across the three species (high values denote high transcriptional conservation). Top versus bottom DE value (Q-value) distributions of bottom and top gene sets are compared using a Mann-Whitney test (one-sided in human and one-sided in the mouse and *R. aegyptiacus* DE analysis). See full volcano plots, including both Q-values and fold change, in Fig S7g. **(g)** Gene evolutionary age analysis of top and bottom genes (based on human genes and their age estimation using three different approaches, and taken from proteinHistorian^39^). Age estimation distributions of bottom and top gene sets are compared using a one-sided Mann-Whitney test. **(h)** Heatmaps of gene expression across the 5 bottom-to-top villus regions (v1-v5), of genes involved in lipid transport (GO:0006869) and in anion binding (GO:0043168) in *R. aegyptiacus*, mouse and human enterocytes. **(i)** Schematic summary of evolutionary analysis of enterocyte differentiation along the villus: top landmark genes tend to be evolutionarily younger, transcriptionally divergent across species. Among the divergent genes, top landmark genes in human and mouse, associated with absorption of specific nutrients, transcriptionally diverge in *R. aegyptiacus* villus. (***=P<0.001)

We then employed principal component analysis (**PCA**) of single-cell data from human, mouse and *R. aegyptiacus* enterocytes. Expression of genes belonging to each of the five regions along the villus showed that PC1 captures the spatial bottom-to-top differentiation dynamics in each of the three species (**Fig 2c** and **Fig S7c&f**). This allowed us to use this axis to define enterocytes belonging to the bottom, intermediate and top regions of the villus in each of the species. We next employed scVelo (version 0.2.4)^38^ to infer the velocities of enterocyte transcriptional states in the single-cell data. We again observed that the inferred velocities largely recapitulate the transcriptional changes from bottom-to-top villus (**Fig 2d** and **Fig S7d**, for bat and mouse analysis, respectively).

We next focused on gene expression along the above-mentioned PC1 with gene markers from each of the five regions in the *R. aegyptiacus* and the mouse single-cell data (**Fig 2e** shows bat analysis, and mouse analysis is in **Fig S7e**). In agreement with the above analysis, we observed that the sets of zonation marker genes are most highly expressed in the expected region along PC1 axis. For example, when looking at the first set of genes (left-most panel in **Fig 2e**), we observed that their *R. aegyptiacus* orthologs tend to be highly expressed in enterocytes inferred to be located in the bottom villus region. This is also true in mouse genes (**Fig S7e**). However, when looking at individual genes, we also observed genes that deviate from expected transcriptional patterns between species, e.g., genes expected to be expressed mostly in the top of the villus, are expressed in other regions of the bat villus (right-most panel in **Fig 2e**). In summary, the significant correspondence of landmark gene expression along the villus of the three studied mammals suggests an overall conservation of the bottom-to-top enterocyte expression program. However, certain genes diverge in expression patterns between species and their characterization is the focus of the following section.

### Top villus genes are transcriptionally divergent and evolutionarily younger than bottom villus genes

Following this, we quantified the transcriptional conservation of top versus bottom genes across species. For this, we performed a DE analysis between top and bottom enterocytes in each of the three species. We then compared the transcriptional divergence in top and bottom gene classes (after controlling for biases in DE values between the two sets of genes). We observed that bottom genes are more transcriptionally conserved across species whereas top genes tend to be divergent (**Fig 2f and Fig S7g**). Thus, a gene that is highly expressed in the bottom villus region in one species is likely to be expressed in a similar manner in other species. In contrast, genes expressed in top regions of the villus in one species tend to diverge in their transcriptional patterns across species.

Next, we compared the evolutionary age of the top and bottom villus genes, using several gene age estimations from ProteinHistorian^39^. We observed that top villus genes are evolutionarily younger than bottom villus genes (**Fig 2g**). This trend was also observed when looking at gene age distribution across all 5 consecutive villus regions (**Fig S7h)**. Thus, the set of genes expressed in the top villus region tends to be rewired in the course of evolution, with higher transcriptional divergence between species, and includes evolutionarily younger genes. In contrast, genes expressed at the bottom of the villus are transcriptionally conserved and are evolutionarily more ancient.

Finally, we asked which genes highly expressed in the top villus significantly diverge between *R. aegyptiacus* and the other two species. Interestingly, the two most striking sets of genes observed are those involved in lipid transport and in anion binding. Most genes from these sets are expressed in the top of the villus in both human and mouse. In *R. aegyptiacus*, however, they are mostly expressed in lower regions or the bottom of the villus (**Fig 2h**). We note that we did not observe such consistent patterns with sets of other genes involved in nutrient absorption and transport (see **Fig S8**).

In summary, the dynamics of gene expression in enterocytes as they migrate along the villus is largely conserved across the studied mammals. However, the expression program in the top villus is more divergent, enriched with evolutionarily young genes, and involves specific transcriptional changes in genes associated with nutrient absorption in *R. aegyptiacus* (**Fig 2i**).

### Complement system genes are expressed in *R. aegyptiacus* intestinal epithelial cells

In the above analysis, we observed that specific classes of nutrient absorption genes diverge in their transcriptional patterns along the villus between species. In contrast to our initial expectations, we did not observe such evolutionary changes in innate immune pathways. This still leaves the possibility that steady-state expression levels of innate immune genes differ in bat cells with respect to mouse and human. We thus performed DE analysis between homologous cell clusters in the gut between species, comparing orthologous gene expression. We searched for genes more highly expressed in *R. aegyptiacus* in comparison with both human and mouse cells. When looking for pathways enriched in such DE genes, we found many terms related to the complement system, including early and late stages of both the classical and the alternative pathways (see, for example, **Table S2** for enriched terms in *R. aegyptiacus* stem cells). Specifically, we observed that many of the complement components (C3,4,5,6,7,9) and complement-associated proteases (C2,CFB,CFD,CFI) were significantly more highly expressed in several types of *R. aegyptiacus* epithelial cells, mostly stem cells, TA and enterocytes, in comparison to human and mouse cells (**Fig 3a-c, Fig S9** and **Table S3&4**). Importantly, many genes of the membrane attack complex, a central immune effector and the final product of the complement cascade^40^, including C5,C6,C7 and C9, are more highly expressed in *R. aegyptiacus* gut epithelia. CFI, an important protease that acts as a regulator of complement activation, is also more highly expressed in *R. aegyptiacus* cells than in both human and mouse. This pattern of higher expression of complement system genes in *R. aegyptiacus* epithelial cells is not observed in most non-epithelial cell types (**Fig S8**). We then asked whether such transcriptional signatures of higher complement gene expression in bat cells also occur in other innate immune genes. However, comparison of the expression of a large set of genes from diverse innate immune pathways (taken from innateDB^41^, see **Table S5**) between *R. aegyptiacus,* mouse and human showed no such pattern in other pathways apart from the complement system (**Fig 3b**).

**Figure 3:**
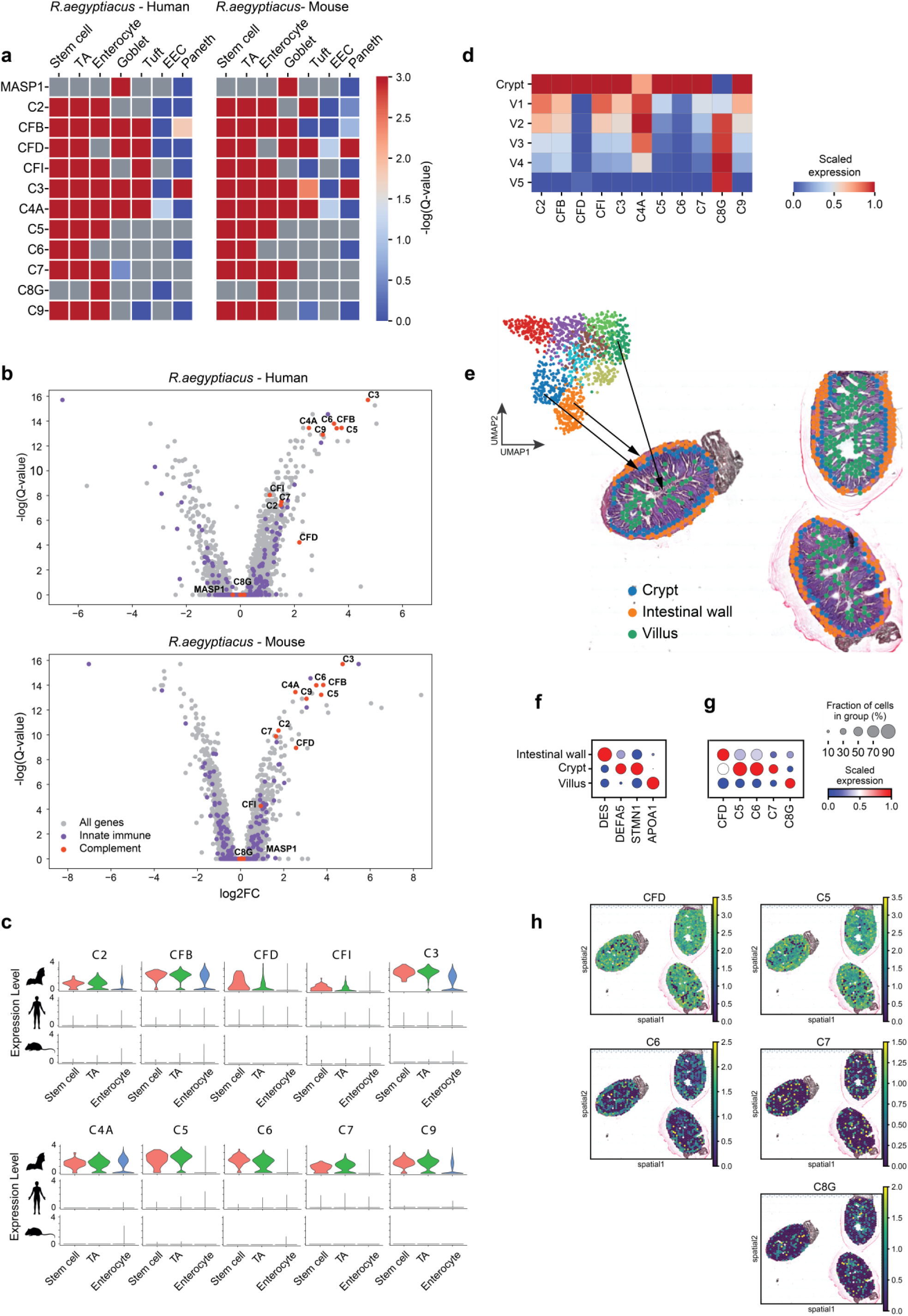
Comparative analysis of complement system gene expression in epithelial cells in the intestine. (**a**) Matrix plots showing DE results (-log[Q-value] between two species) of complement genes, between *R. aegyptiacus* and human (left) and *R. aegyptiacus* and mouse (right) homologous epithelial cells. Maximum values are trimmed at 3 (Q-value<10^-3^). In all cases of significant differences between species, the expression is higher in *R. aegyptiacus*, except for C2 in mouse Tuft cells, and for C8G in human enterocytes. **(b)** Volcano plots showing DE analysis between *R. aegyptiacus* and human (top) or mouse (bottom) epithelial stem cells. Complement system genes appear in red, innate immune genes (from innateDB^41^) in purple, all other genes in grey. **(c)** Violin plots of complement component and protease genes across different epithelial intestinal cell types in *R. aegyptiacus*, human and mouse. **(d)** Scaled expression levels of complement genes at the crypt (SC, TA and Paneth cells) and along the five regions of the villus (enterocytes) in *R. aegyptiacus* gut, using scRNA-seq and localization based on Fig 2c. **(e-h)** Analysis of complement gene expression using 10x Visium data: (**e**) Top: UMAP of grid spots from the *R. aegyptiacus* spatial transcriptomics data. Bottom: Localization of spots, colored by clusters from upper UMAP, in cross-sections of *R. aegyptiacus* gut, showing that the orange, blue and green clusters refer to the intestinal wall, crypt and villus regions. (**f-g**) Dot plots showing relative expression in the three clusters that refer to different gut regions (as shown in 4e). (**f**) known marker genes, and (**g)** selected complement genes. (**h**) Expression levels of selected complement genes across gut sections (see all complement genes in Fig S10).

We next analyzed the spatial expression patterns of complement system genes across *R. aegyptiacus* gut. Using our single-cell data, we observed that many of these genes tend to be expressed most highly in the crypt and then gradually decrease in expression along the villus (**Fig 3d**). These observations were further strengthened using the spatial transcriptomics data, where we clustered the spots into spatial regions (crypt, villus, and intestinal wall (which refers to outer layers, including submucosa, muscularis layers and serosa)) (**Fig 3e**), and estimated the expression of complement system genes in these regions (**Fig 3f-h** and **Fig S10a**, and **Fig S10b** for comparison in mouse). In summary, complement component and protease genes are uniquely expressed in *R. aegyptiacus* epithelial cells in comparison with human and mouse cells, and this expression is strongest in the crypt for the majority of these genes.

### Complement system genes are highly expressed in *R. aegyptiacus* epithelial lung cells

Following our gut analysis, we asked whether similar signatures of complement system gene expression can be observed in epithelial cells in other barrier tissues in *R. aegyptiacus*. To investigate this, we profiled 60,000 high-quality lung cells from 8 individual bats, and identified 45 cell clusters (**Fig 4a and Fig S11**). In the lung epithelium, we identified Alveolar type 1 and 2 (AT1 and AT2, respectively), club, goblet, brush (tuft) and basal cells. To compare these cells with other mammals, we profiled over 21,000 cells from mouse lungs (see **Fig S12**), using the same protocol as the one we used with bat lungs, and also utilized publicly available lung data from the human cell atlas^42^. Using correlations between cells across species and cross-species dataset integration, we observed significant transcriptional similarity between homologous cell populations across species (**Fig S13** and **Fig S14**).

**Figure 4:**
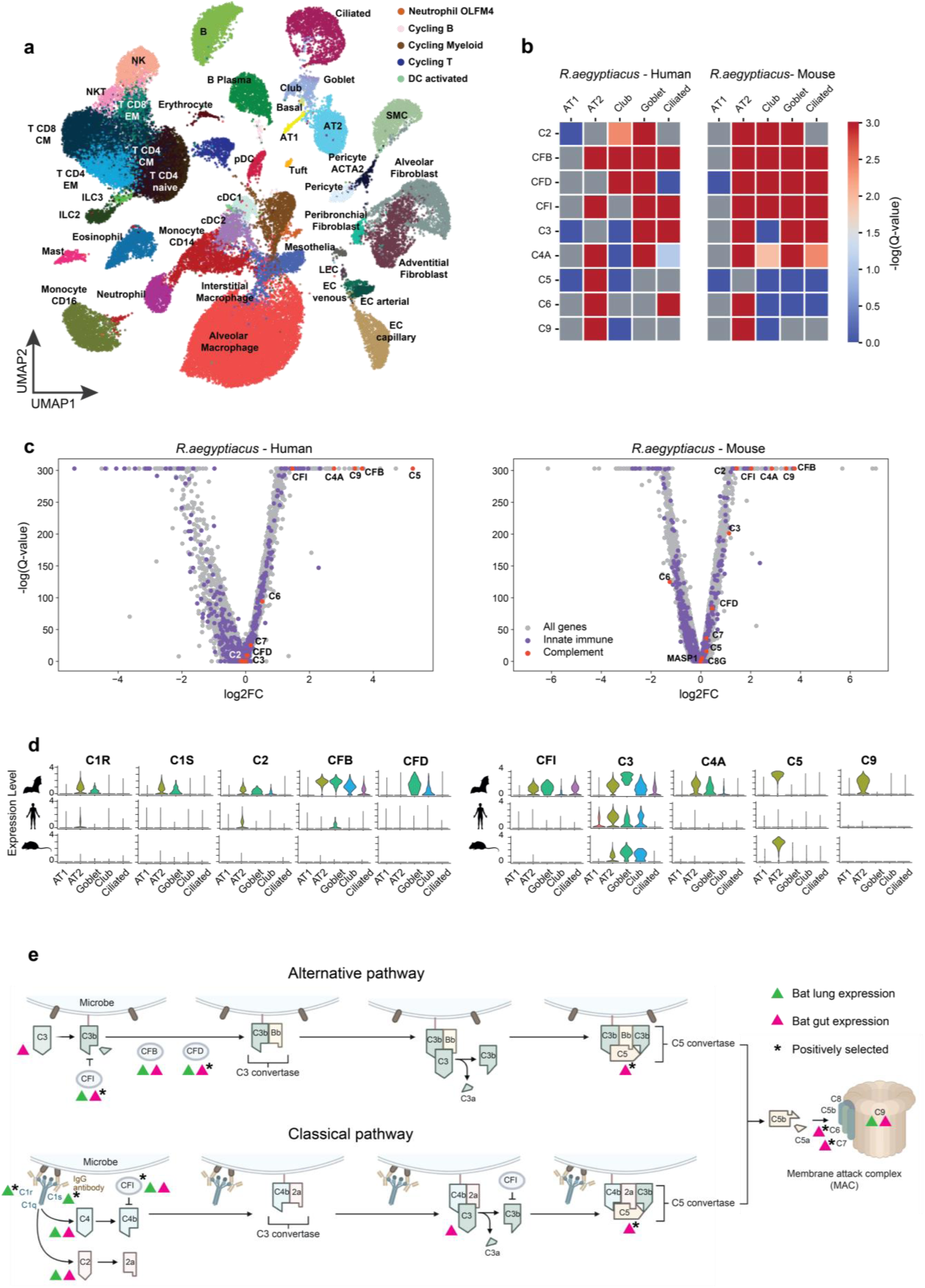
Transcriptional comparison of lung-resident epithelial cells from *R. aegyptiacus*, mouse and human. (**a**) UMAP of *R. aegyptiacus* lung cell clusters (10 samples, n=60,000 cells). **(b)** Matrix plots showing DE results (-log[Q-value] between two species) of complement genes, between *R. aegyptiacus* and human (left) and *R. aegyptiacus* and mouse (right) homologous epithelial cells. Maximum values are trimmed at 3 (Q-value<10^-3^). In all cases of significant differences between species, the expression is higher in *R. aegyptiacus*, except for C6 expression that is higher in mouse AT2 and in human ciliated cells. **(c)** Volcano plots showing DE analysis between *R. aegyptiacus* and human (left) or mouse (right) AT2 cells. Complement system genes appear in red, innate immune genes (from innateDB^41^) in purple, all other genes in grey. **(d)** Violin plots of complement component and protease genes across different epithelial lung cell types in *R. aegyptiacus*, human and mouse. **(e)** Schematic of the complement pathway overlayed with several evolutionary analyses – genes expressed uniquely in *R. aegyptiacus* gut and lung epithelial cells are marked in pink and green, respectively. Genes with signatures of positive selection across bat species are marked with *. C8A that is part of the C8 complex together with C8B and C8G, also displays signatures of positive selection.

As in the gut, we performed DE analysis between homologous cell clusters between species to find transcriptionally divergent genes. Interestingly, we again observed that a set of complement system genes is more highly expressed in various *R. aegyptiacus* epithelial cells, including in ciliated, club, goblet and, most prominently, in AT2 cells (**Fig 4b-c** and **Table S6-9**^43^). The expression of these complement genes was unique to *R. aegyptiacus* in most of these epithelial cell types in comparison with mouse and human, and included genes expressed in early stages of the classical pathway (C1S, C1R,C2 and C4A) and in the alternative pathway (CFB and CFD) (**Fig 4d and Fig S15**). Most late-stage genes of these pathways do not differ between species, with the exception of C9 that is higher in bat AT2 cells. The results regarding complement expression in human and mouse cells were corroborated by previous analyses of unrelated single-cell lung data^44^. The set of complement system genes more highly expressed in *R. aegyptiacus* lung epithelium overlaps with genes expressed more highly in *R. aegyptiacus* gut epithelium, but is not identical. As in the gut epithelium, innate immune genes in general are not more highly expressed in bat lung cells in comparison with human or mouse cells (**Fig 4c**). In addition, these expression patterns of complement system genes were not observed in most non-epithelial cell types in the lung. Thus, *R. aegyptiacus* epithelial cells from both the lung and the gut uniquely express an array of complement system genes.

### Complement system genes display signatures of positive selection across bat species

Following our observations of transcriptional differences between *R. aegyptiacus* and human and mouse cells in basal expression of numerous components of the complement pathway, we next asked whether these genes also evolve rapidly in their coding sequences. We employed a site model implemented in PAML^45^ to test for positive selection in coding sequences of 14 complement system genes using orthologous genes from different bat species representative of the Chiroptera order. We observed that 8 of the 14 genes show strong statistical evidence of positive selection (see **Table S10**).

In summary, we observed significant evolutionary changes in both basal expression in lung and gut epithelia as well as in coding sequences in bat complement system genes. **Figure 4e** summarizes these changes along the classical and alternative complement pathways. These observations are particularly interesting when combined with recent findings regarding the evolution of other innate immune genes encoding for secreted proteins, such as defensins that have been lost in bat genomes through pseudogenization^27^. Unconstrained by intra-cellular interactions and complex protein networks, secreted proteins can have a greater capacity to evolve. Secreted immune proteins, such as complement proteins and defensins, can thus serve as ‘hotspots’ for evolutionary changes that can shape and fine-tune the innate immune response. We elaborate on this and on the wider context of our results regarding complement system evolution in Discussion.

### Systematic comparison of antibacterial and antiviral innate immune responses in PBMCs across cells and species

Thus far we analyzed innate immune gene expression in bat tissues in steady-state conditions. We next focused on the innate immune response, the expression program triggered following pathogen infection that involves the upregulation of numerous innate immune genes, and asked how this transcriptional response has evolved in bats. For this, we profiled nearly 85,000 bat and mouse PBMCs in steady-state conditions as well as following stimulation with two pathogen-associated molecular patterns (**PAMPs**): dsRNA (poly I:C) and lipopolysaccharide (LPS), which respectively trigger the antiviral and the antibacterial responses. These are conserved and well-characterized PAMPs that enable studying the unmodulated host response to pathogens in a comparative manner across cells and species while avoiding host-pathogen-specific biases^46^. Among the profiled PBMCs, we identified 20 major cell populations (**Fig 5a-b** and **Fig S16-17**). These included monocytes, dendritic, T, NK, and B cells as well as neutrophils. Marker gene-based correlations with annotated mouse and human PBMC data further confirmed our cell type annotations in *R. aegyptiacus* (**Fig S18-19**). Some of the clusters we observed are enriched with cells from a particular stimulation condition, such as a monocyte cluster enriched with dsRNA-stimulated cells (**Fig 5b**).

**Figure 5:**
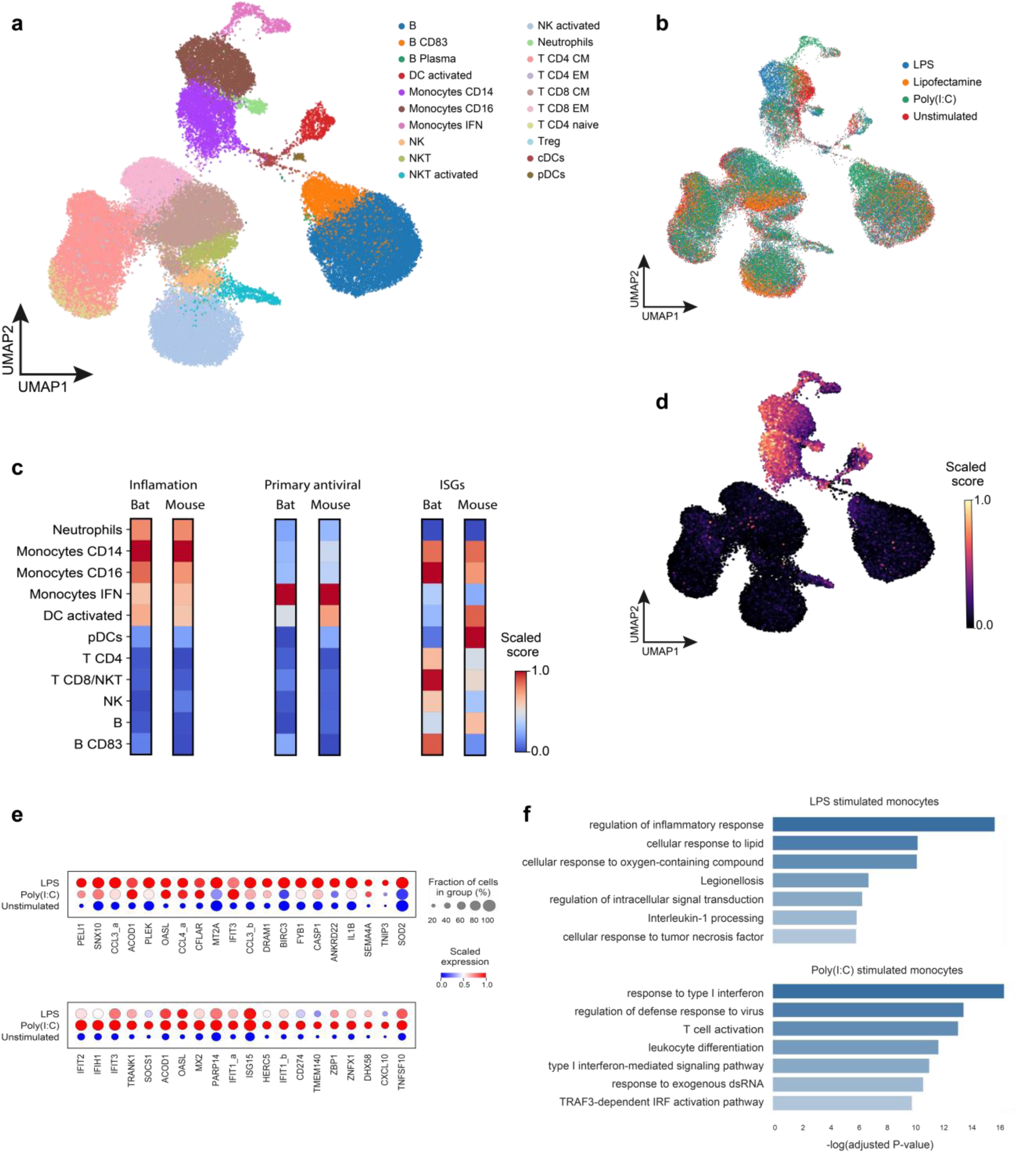
Comparative analysis of PBMC gene expression and transcriptional response to innate immune stimuli between *R. aegyptiacus* and mouse. (**a-b**) UMAP of *R. aegyptiacus* PBMC clusters (5 samples, n=59,443 cells). Cell distribution, colored by condition is shown in (b). **(c)** Normalized expression scores of selected sets of innate immune genes across different PBMCs in *R. aegyptiacus* and mouse: inflammatory score, primary antiviral score and ISG score (see main text for details). **(d)** UMAP of *R. aegyptiacus* PBMCs colored by inflammatory score, showing most inflammatory genes are upregulated in monocytes and DCs. **(e)** Dot plots showing expression levels and percentage of cells expressing the 20 most upregulated genes in *R. aegyptiacus* monocytes in response to LPS (top) and dsRNA (bottom) stimulation. Genes identified during orthology were named as follows (original identifier in brackets): CCL3_a (LOC107506475), CCL3_b (LOC107506481), CCL4_a (LOC107508347), IFIT1_a (LOC107501624), IFIT1_b (LOC118603801). **(f)** Enrichment analysis of the 100 most upregulated genes in response to LPS (top) and dsRNA (bottom), showing GO terms unique to each of the two gene sets (distributions of –log[Q-value] are shown as bars).

Next, we systematically compared the response to stimuli across cell clusters, by computing the numbers of DE genes between stimulated and control cells for each such cell cluster (using an equal number of cells per cluster). We observed that in both antiviral and antibacterial responses and in both mouse and *R. aegyptiacus*, monocytes have the highest numbers of DE genes across cells (**Table S11**). We then dissected the response in a comparative manner across different cell clusters by quantifying the expression of genes known to be associated with different stages and modules of the innate immune response (**Fig 5c-d**). We asked which stimulated cell types tend to upregulate sets of genes belonging to different stages and modules of the innate immune response, and how conserved this expression profile is between mouse and *R. aegyptiacus.* We used two sets of innate immune genes upregulated during early stages of the immune response, namely, primary antiviral response^46^ and inflammatory genes^47^, and a set interferon-stimulated genes (**ISGs**)^48^, that are upregulated in the second wave of response. We observed that the transcriptional patterns of the two primary responses across cell types are more conserved than the ISG response: Genes related to inflammation and to the primary antiviral stage are mostly expressed in monocytes in both mouse and *R. aegyptiacus* (with lower levels of upregulation of inflammatory genes in DCs and neutrophils). In contrast, ISGs, upregulated during the secondary wave of response, are highly expressed in a range of different cell types in both species, but their relative levels differ between the species (**Fig 5d**). Overall, this comparison between mouse and *R. aegyptiacus* suggests a conserved innate immune response across PBMC types in the initial stages of response to infection, followed by greater divergence in later stages.

### Similarity and divergence in antiviral and antibacterial gene upregulation in *R. aegyptiacus* and mouse cells

The transcriptional responses to viral and to bacterial PAMPs were previously compared in human populations^49^ and in mouse models^50^. We thus asked how similar the two responses are in *R. aegyptiacus* cells. We examined the sets of genes upregulated in the antiviral and the antibacterial responses in *R. aegyptiacus* monocytes, the cells that display the strongest reaction according to the above analyses. As expected, the two responses significantly overlapped in terms of shared upregulated genes. However, the relative levels of upregulation vary between cell types (**Fig 5e**). For example, in the LPS response, several cytokines and chemokines, such as CCL3, CCL4 and IL1B, are more strongly upregulated than in LPS response, while the levels of major dsRNA sensors, MDA5 and RIG-I, are higher in dsRNA response. By looking at GO terms enriched in the top 100 upregulated genes in each response (using g:Profiler^43^, see **Table S12**), and searching for terms specific for either the antiviral and the antibacterial responses, we observed subtle distinctions between them (**Fig 5f**). The antibacterial response had terms associated with inflammation as well as with response to lipid and bacterial infection, while the antiviral response was associated with upregulation of dsRNA-sensing genes and with type I IFN response. These distinctions, identified in our *R. aegyptiacus* expression data, are in agreement with the expected differences and previous analysis of the two responses^46^. This points to an overall conservation in the regulation of the immediate pathogen-specific innate immune response among the examined mammals.

We next searched for genes more strongly upregulated in bat cells in comparison with mouse in either the antiviral or the antibacterial responses. This was done by comparing the fold change in response to stimulus in homologous cells in bat and mouse in three major types of PBMCs, CD4 T cells, B cells and CD14 monocytes, where a sufficient number of cells enabled statistical testing (this comparison was done after downsampling to have an equal number of cells across conditions and species). We then filtered for genes that are more strongly upregulated in all three cell types in bats versus mouse, and vice versa (**Table S13**). We found that genes more strongly upregulated in bat in the antiviral response across cell types include dsRNA-binding proteins: the three RIG-I-like receptors, RIG-I, MDA5 and LGP2, that are central for viral RNA recognition and initiation of the innate immune response, and PKR that upon binding to viral RNAs inhibits translation as part of the cellular defense against viral replication. In contrast, genes more highly upregulated in mouse in both antibacterial and antiviral responses include genes in the STAT signaling pathway, including PARP9, PARP14 as well as STAT1 and STAT2 themselves. In summary, this comparison found specific modules, involved in viral recognition and downstream signaling, that are more strongly upregulated following infection in a species-specific manner.

### Conservation of cytokine and stress-related gene upregulation in monocytes

Next, we focused on the dynamics of the antiviral response in *R. aegyptiacus* monocytes and its conservation in mouse. Among bat monocytes, we observed a cell cluster that is almost entirely composed of cells stimulated with dsRNA. Cells belonging to this cluster have high inflammatory and antiviral scores (**Fig 5a,c-d**). In addition, IFNB1, a major antiviral cytokine, is almost exclusively upregulated in this cluster. This cluster (termed ‘IFN Monocytes’) is also observed in mouse dsRNA-stimulated monocytes (**Fig S17**). To further investigate the architecture of the monocyte antiviral response, we compared the three major states of *R. aegyptiacus* CD16 monocytes: the two groups of dsRNA-stimulated cells (IFN and non-IFN) and a third group with all remaining cells in unstimulated or mock-stimulated conditions (we excluded LPS-stimulated cells from this analysis to focus on the antiviral response). We found genes that are expressed significantly higher in each of these three groups in comparison with the other two groups (termed “state-DE genes”). Using hierarchical clustering of the three sets of state-DE genes, we observed that the majority of genes upregulated in IFN monocytes, 263 out of 424 genes, are uniquely upregulated in this state and are not significantly expressed in other monocytes (**Fig 6a**). In addition, we observed hundreds of genes that are downregulated in IFN monocytes in comparison with the other two states. Thus, this monocyte cell-state is distinguished in unique differential expression of a large number of genes in comparison with both unstimulated and other dsRNA-stimulated CD16 monocytes.

**Figure 6:**
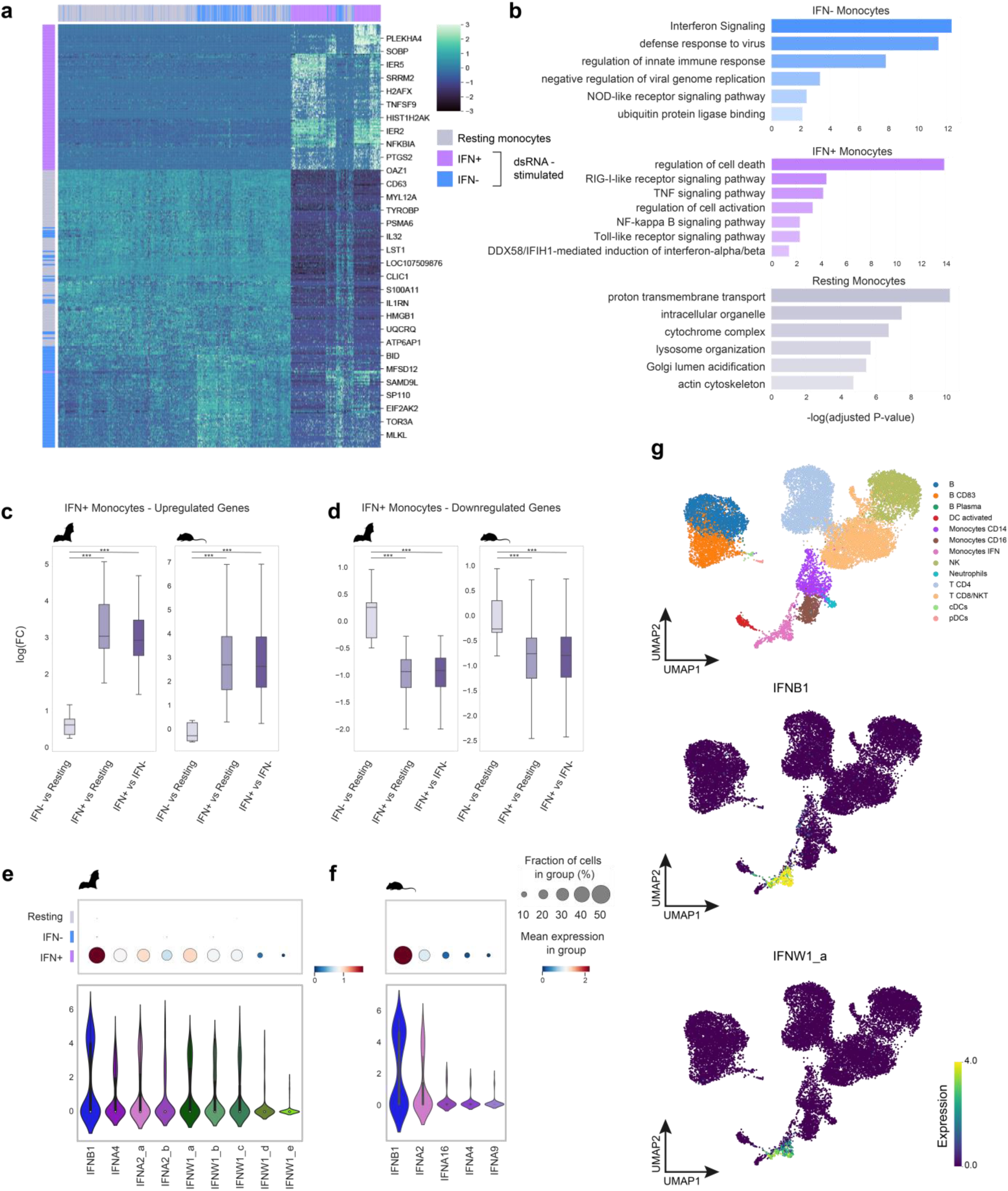
Comparison of the transcriptional antiviral response between *R. aegyptiacus* and mouse monocytes. (**a**) Hierarchical clustering heatmap (“ward” as clustering method) showing expression of the three sets of state-DE genes from unstimulated monocytes and the two dsRNA-stimulated monocytes clusters (with and without IFN upregulation) in *R. aegyptiacus*. **(b)** Enrichment analysis of state-DE genes, showing GO terms unique for each of the three sets of genes (as in 7a). **(c-d)** conservation of transcriptional response of genes in IFN monocytes between *R. aegyptiacus* and mouse: Fold changes (logFC) between each pair of the monocyte clusters are shown for up– and down-regulated genes (c and d, respectively). These up– and down-regulated genes are unique to the IFN monocyte cluster in *R. aegyptiacus*. In Each panel the logFC distribution is shown in *R. aegyptiacus* monocytes (left panel) and in mouse monocytes (right panel). Distribution of logFC between the sets are compared using a one-sided Mann-Whitney test ((***=P<0.001)). **(e)** Top: Dot plots showing various type-I IFNs (IFNAs, IFNB and IFNWs) expression level in *R. aegyptiacus*. Bottom: Violin plots showing the distribution of expression level of various type-I IFNs across *R. aegyptiacus* IFN monocytes. IFNA2_a (LOC107511568), IFNA2_b (LOC107508426), IFNW1 a-e (LOC107518592, LOC107518590, LOC107518583, LOC107518588, LOC107518582). **(f)** As in (e), in mouse monocytes. **(g)** UMAP of dsRNA-stimulated PBMCs, colored by cell type (top), and showing expression levels of IFNB and two of the IFNW duplicates in *R. aegyptiacus* (see the expression of additional type-I IFNs and similar analysis of mouse PBMCs in Fig S20).

To characterize the pathways enriched in genes expressed in each of these states, we used GO term analysis of the three sets of state-DE genes (using g:Profiler^43^). We observed that the group of unstimulated monocytes is enriched with processes related to resting monocyte functions, including mitochondrial and lysosomal processes, as also observed in a similar system in human^51^ (**Fig 6b and Table S14**). Genes upregulated in IFN monocytes are enriched with processes related to cell death as well as to TNF signaling and to cell activation, in agreement with the set of cytokines extensively expressed in this group of cells. Finally, genes strongly upregulated in non-IFN dsRNA-stimulated monocytes are associated with various functions related to the intracellular defense program, in line with their broad upregulation across stimulated monocytes. These upregulated genes include restriction factors against viral replication and numerous genes that regulate and limit the response to maintain healthy tissue following infection. In summary, the two dsRNA-stimulated cell states represent two monocyte sets, one that expresses a range of cytokines and gene related to cell death and inflammation and another set that upregulates a broad antiviral gene response.

We next asked whether the unique transcriptional patterns observed in *R. aegyptiacus* IFN monocytes are also observed in the corresponding mouse cells. We observed that the one-to-one orthologs of genes uniquely upregulated in *R. aegyptiacus* IFN monocytes are also upregulated in mouse IFN cells in comparison with the other two monocyte populations (**Fig 6c**). Furthermore, the orthologs of *R. aegyptiacus* genes that are downregulated in *R. aegyptiacus* IFN monocytes also show significant low expression levels in mouse IFN monocytes, suggesting a conserved architecture of transcriptional response to dsRNA in monocytes in the two species. These findings, of a small subset of monocytes that highly express antiviral cytokines and cell-death related genes in both mouse and bat, are in agreement with studies of human blood and bone-marrow derived monocytes, infected with bacteria and influenza^51,52^. Thus, the upregulation of IFN and other cytokines in a limited set of monocytes is conserved across species and infections.

### Comparison of transcriptional patterns of recent duplications of IFNs in *R. aegyptiacus* and in mouse

One of the major discoveries in the *R. aegyptiacus* genome was the finding of an expanded set of IFN Omega (IFNW) duplicates^19^. While a single IFNW gene exists in the human genome, nearly half of the *R. aegyptiacus* IFN genes belong to the IFN-ω family^19^. In contrast, *R. aegyptiacus* has fewer copies of IFNA – another type-I IFN that has numerous paralogs in both rodents and primates. Thus, type-I IFN gene families have undergone significant contraction and expansion during mammalian evolution. A previous analysis found differences in the set of ISGs upregulated upon binding of different types of *R. aegyptiacus* IFNW^18^. We thus asked whether these recent IFNW duplicates also show transcriptional divergence in response to dsRNA, and compared the expression of various type-I IFNs in mouse and *R. aegyptiacus* cells. We observed that among different types of PBMCs, IFNs are almost entirely expressed in monocytes (**Fig S20**). Additionally, the expression patterns of all *R. aegyptiacus* IFNW duplicates are similar to each other – they are all upregulated specifically in the IFN monocyte cell-state and their expression levels are significantly lower than that of IFNB1 (**Fig 6e-g**). The same transcriptional behavior is observed with the few existing copies of *R. aegyptiacus* IFNA. Interestingly, this mirrors the same pattern of type-I IFN expression in mouse cells: IFNB1 and all other type-I IFNs are almost entirely upregulated in IFN monocytes, and IFNB1 is significantly higher in its expression in comparison with mouse IFNAs (**Fig 6f and Fig S20**). Thus, in both mouse and *R. aegyptiacus*, recent duplicates of type-I IFNs (either IFNW or IFNA) are expressed in lower levels than the ancient and conserved single copy of IFNB1. These recent IFN duplicates share the same regulation with more ancient IFN genes, being exclusively expressed in a subset of cells that also upregulates additional cytokines, such as TNF, and genes related to apoptosis, such as DUSP1 and FOS. In summary, our findings and previous work show that IFNB1 is the major IFN upregulated in response to dsRNA in a range of mammals (primates, rodents, ungulates^46^ and also bats) and cells^20,46^. Other type-I IFNs, including recently duplicated copies, tend to be co-expressed in the same cells as IFNB1, but in lower levels, suggesting a strong regulatory conservation of upregulation of type-I IFN genes.

## Discussion

Previous studies using comparative single-cell analysis in various species gave insights into the origins and evolution of the immune system^46,53–55^, as well as to deeper understanding of cell type^56–60^ and tissue evolution^61^. In this work, we have profiled a comprehensive dataset of single-cells from *R. aegyptiacus* lung, gut and blood, in a comparative manner to cells from mouse tissues and to existing human single-cell data, to map transcriptional conservation and divergence in pathways associated with environmental and pathogen interactions. We have also augmented this single-cell data by performing innate immune stimulation and with spatial transcriptomics, which allowed us to study the conservation of dynamic gene expression programs and to analyze them both temporally and spatially.

Following characterization of *R. aegyptiacus* gut cells, we focused on the transcriptional evolution of enterocyte differentiation along the villus axis – a genetic program important for immune defense, interactions with microbes and nutrient absorption. It was previously shown that during their migration along the mouse gut villus, enterocytes express different gene sets that support region-specific functions^37^. By combining spatial and single-cell transcriptomics from *R. aegyptiacus* gut samples, and comparing gene expression between mouse, human and bat enterocytes, we found a significant correspondence between landmark gene expression along the villus, suggesting an overall conservation of this program. However, we also observed that top-villus enterocytes display the highest transcriptional divergence and that these enterocytes express genes that are evolutionarily younger in comparison with bottom-villus enterocytes. Our findings, of higher evolutionary divergence in later stages of enterocyte differentiation, are in line with comparative transcriptomics analyses of spermatogenesis^62^ and of embryonic development of mammalian tissues^63^. In these works, the authors showed gradual decrease in transcriptional similarity between species with developmental time, and enrichment of recently born genes in later developmental stages. Finally, in the top villus enterocytes of *R. aegyptiacus*, we found a significant depletion in expression of genes associated with absorption and transport of lipids and anions. These transcriptional changes likely stem from the large differences in diet and in gut structure and function between the frugivore *R. aegyptiacus*, and mouse and human, and demonstrate how these adaptations are reflected at the molecular level in bat cells.

We next searched for innate immune genes that differ in expression between *R. aegyptiacus*, human and mouse. Steady-state expression of immune genes in barrier tissues can have important consequences for species ability to resist invading pathogens, and is thought to be an important mechanism in the adaptation of several bat species to viral infection^64,65^. We found that in contrast to the to the majority of innate immune genes that do not significantly differ in steady-state expression between the species in either lung or gut cells, many complement system genes are uniquely expressed in bat epithelial cells from both tissues. Interestingly, recent evolutionary analyses comparing bat genes with orthologs in other mammals found evidence for lineage-specific adaptation in coding sequences of bat complement system genes^66–68^. In agreement with this, we found that a significant fraction of the complement system genes also show elevated evolutionary rates and signatures of positive selection in their coding sequences when compared across bat species. Taken together, our analyses points to concurrent evolutionary divergence in both sequence and expression of complement system genes in *R. aegyptiacus*.

Interestingly, previous studies of various bat species found evidence for evolutionary and functional divergence in other antiviral and antibacterial pathways that, like the complement pathway, involve extracellular secretion of proteins. These include defensins, an important class of secreted peptides that act as effectors against various pathogens or as signaling molecules that can modulate the immune system^69^. It was previously shown that many alpha– and beta-defensin-encoding genes were pseudogenized in bat genomes^27^. Furthermore, several inflammasome sensors are either lost^27,66,70^ or transcriptionally dampened^71^ in bats, leading to reduced secretion of inflammatory cytokines, including IL1B and IL18. Finally, a recent cross-bat evolutionary and functional analysis showed that ISG15, an antiviral factor that has extracellular pro-inflammatory activities in human, has undergone numerous changes in its coding sequence in bats and is not secreted into the extracellular space following expression in bat cells, unlike its human ortholog^68^. We hypothesize that these changes in bat secreted immune molecules and their upstream regulators may contribute to the observed tolerance of bats to various viral infection, for example by fine-tuning the inflammatory response and by reducing subsequent tissue damage. Unlike intracellular proteins that their evolution is constrained by interactions with numerous cellular proteins, secreted proteins may have greater capacity to evolve, as they usually interact with few proteins^46^, enabling rapid evolution in both sequence and expression. Following our analysis of steady-state expression, we studied the transcriptional evolution of induced expression of innate immune genes, by comparing the antiviral and the antibacterial responses in mouse and bat PBMCs. These programs are thought to evolve rapidly as part of the evolutionary arms race with pathogens, but are also under regulatory constraints to avoid an excessive immune reaction that can damage host tissues^46,72^. We observed that the primary stages of the transcriptional responses are more conserved between the species than the secondary stage of response, and that genes more highly upregulated in one of the two mammals belong to specific modules that regulate the response: Genes involved in early viral detection are more highly upregulated in *R. aegyptiacus*, while genes belonging to the STAT signaling pathway are more strongly regulated in mouse cells.

Focusing on the primary antiviral response in monocytes, the most responsive cells to dsRNA, we found that in both mouse and *R. aegyptiacus* a distinct subset of dsRNA-stimulated monocytes expresses IFNs and other cytokines as well as stress-related genes. This is an agreement with previous studies on non-immune cells infected with viruses or stimulated with dsRNA that found a small subset of cytokine secreting cells, suggesting a conserved regulatory architecture of the innate immune response that likely limits the upregulation of cytokines to avoid an excessive immune reaction^46,73^. Interestingly, we find that rapid expansion of type-I IFN genes in both the rodent and the bat lineages, did not lead to transcriptional sub– or neo-functionalization of the lineage-specific IFN duplicates and that they are all co-expressed in the same monocyte subset. Thus, the remarkable gene expansion and contraction of IFNs in the course of mammalian evolution is likely associated with coding sequence divergence that contribute to functional differences in binding of these IFN paralogs and in their downstream signaling^74^.

In summary, our study reveals conserved and divergent modules in innate immune and in environmental interaction pathways, such as nutrient uptake, in *R. aegyptiacus* epithelial and immune cells. This gives insights into the molecular mechanisms of bat adaptation to viral infection and to a high-sugar diet. In addition to being a unique resource for studying bat biology and tissue evolution, these comparative data can also advance our understanding of pathological conditions in humans and provide a basis for novel therapeutics.

## Methods

### Ethical statements

Experimental protocols were approved by the Tel Aviv University Animal Care and Use Committee (04-22-027, 04-21-065 and 04-20-023) according to the Israel Welfare Law (1994) and the National Research Council (*Guide for the Care and Use of Laboratory Animals*, 2010), and by the Israeli National Park Authority (2021/42760). Healthy adult male Egyptian fruit bats (*R. aegyptiacus*) were housed in our facilities in the Zoological Garden at Tel Aviv University in a controlled environment and their health status was monitored by a veterinarian. Healthy C57BL/6J adult male mice were obtained from the Tel-Aviv University animal facility.

### Experimental procedures

Tissue dissociation was performed using protocols that were as similar as possible to protocols used in previous work on human tissues^30,75,76^. Cells mapped from these human tissue dissociations were used as comparisons to *R. aegyptiacus* and mouse cells in some of the analyses below.

### Isolation of gut cells for single-cell transcriptomics

Gut tissue sections, specifically the lower part of the upper third intestinal tract, from eight bats were collected following transcardial perfusion with cold D-PBS (02-023-1A, Biological Industries). Samples were cleaned and then sliced longitudinally, cleaned and again washed with D-PBS. Next, samples were minced and 0.15-0.2g of the minced tissue was transferred into the digestion mix (1.07 wU/ml Liberase DH (Roche, 5401054001), (70 U/ml) Hyaluronidase (Sigma-Aldrich 385931-25KU), (70 ug/ml) DNAse I (11284932001, Roche) in HBSS buffer (H6648, Sigma-Aldrich)). The samples were incubated for 25 minutes on a shaking platform at 150rpm, and gently mixed every 10 min. Following incubation, the cells were passed through a 40mM filter into a 50ml falcon and added with 12ml of Neutralization media (DMEM, Rhenium (41965039), 10% Heat inactivated FBS (Rhenium, 10270106)). The cells were pelleted using centrifugation (400 G, 4C for 5 minutes), resuspended in DMEM and counted. Depending on red blood cell (**RBC**) presence in the pellet, samples were then treated with RBC cell lysis (Sigma, R7757) according to the manufacturer’s protocol), quenched with 10% FBS in D-PBS,and again pelleted and resuspended. Dead-cell removal protocol (Biotec Miltenyi, 130-090-101) was applied according to the manufacturer’s protocol in cases when cell viability was lower than 75%. The flowthrough was then pelleted at 400G and resuspended in PBS with 0.05% BSA. Cells were counted and loaded on the 10X Genomics Chromium instrument for single-cell library preparation, aiming to capture 10,000 cells, using 3’ v2 chemistry according to the manufacturer’s protocol. Libraries were sequenced on an Illumina NovaSeq 6000 using the NovaSeq 6000 SP Reagent Kit v1.5 (100 cycles) to generate 84-bp paired end reads.

Mouse small intestine (**SI)** and **colon** samples (5 and 2, respectively) were processed in a similar manner. In the case of SI, we took samples from the jejuno-ileal region. In the case of colon, we took the whole colon. Incubation time with digestion mix was 25 minutes.

### Preparation and processing of gut sections for spatial transcriptomics

Gut sections were frozen in OCT according to 10X Genomics Visium protocol. Then 10x Genomics Visium protocol was applied on the (OCT)-embedded fresh frozen samples. All tissues were sectioned using a Leica CX3050S cryostat and were cut to obtain a 10 µm width. Samples were then tested for RNA integrity using a Tapestation 4200. Only blocks with RIN above 7 were used. Tissue optimization was performed, resulting in a permeabilization time of 23 minutes. The Visium spatial gene expression protocol from 10X Genomics was applied by loading a total of 11 gut sections from two individual bats onto the Library Preparation Slide (4 grid sections) and following the manufacturer’s protocol. All images for this process were scanned at 40× with an Aperio Versa 200 slide scanner. Libraries were sequenced on an Illumina NovaSeq 6000 using the NovaSeq 6000 SP Reagent Kit v1.5 (100 cycles) to generate 84-bp paired end reads.

### Isolation of lung cells for single-cell transcriptomics

Lung tissues of 8 bats were removed following transcardial perfusion and washed in cold D-PBS (Biological Industries, 02-023-1A). Trachea was removed and the tissue was immediately injected with an enzyme mix based on previous work^76^ (Dispase II 50 caseinolytic u/ml (Sigma-Aldrich, D4693), Elastase 4.3 u/ml (Worthington Biochem, Worthington, LS002292)), Dnase I 30 μg/ml (Roche), Collagenase A 2 mg/ml (Roche, 10103578001), CaCl2 5 Mm (Sigma)) followed by mincing and a brief incubation in a CO2 incubator at 37 °C. Samples were then transferred to a shaker-incubator for 20 min at 37 °C, 150 rpm, followed by an immediate addition of 15 ml of neutralization buffer (DMEM (Biological industries 01-052-1), 10% FBS (Rhenium, 10270106). Samples were then passed through a 40 um strainer twice and pelleted at 400 G, 5 min at 4 °C. Samples were then treated with RBC cell lysis (Sigma, R7757C, according to the manufacturer’s protocol) and quenched with (10% FBS, D-PBS). Cells were then pelleted at 400 G, 5 min at 4 °C, resuspended in DMEM and counted. Dead cell removal protocol (Miltney, 130-090-101) was applied according to manufacturer protocol in cases when cell viability was lower than 75%, and the flowthrough was pelleted at 400 G, 5 min at 4 °C and resuspended in PBS with 0.05% BSA. Cells were counted and loaded on the 10X Genomics Chromium instrument for single-cell sequencing, aiming to capture 10,000 cells, using 3’ V2 chemistry, according to the manufacturer’s protocol for CellPlex samples, or c_ChromiumNextGEMSingleCell 3’ v3.1_Rev D protocol for the standard protocol Libraries were sequenced on an Illumina NovaSeq 6000 using the NovaSeq 6000 SP Reagent Kit v1.5 (100 cycles) to generate 84-bp paired end reads.

In two cases, we tested an additional protocol: in one individual, we also tested the protocol without injection, and in another individual, we tested a cold dissociation protocol combining a similar digestion mix with an additional Bacillus Licheniformis protease – subtilisin A (Sigma, P5380). The procedure was performed as suggested in protocols.io (dx.doi.org/10.17504/protocols.io.ymgfu3w). For mouse lungs, we followed the same procedure we used for the majority of bat samples, with mouse lungs taken from 3 adult males individuals.

### Isolation of peripheral blood mononuclear cells and stimulation

*R. aegyptiacus* blood was collected from five individuals during transfusion and kept in heparinized tubes to prevent clot formation. peripheral blood mononuclear cells (**PBMCs**) were isolated using 63% Percoll (Sigma, Cytiva;17-0891-01) solution, according to standard density gradient centrifugation methods (30 min, 400G). Samples were then treated with RBC cell lysis (Sigma, R7757), quenched with 10% FBS in D-PBS, and again pelleted and resuspended.

In the case of mice, since the amount of blood from each individual is limited, we combined blood from several individuals (6 and 9 in two separate experiments) before continuing to PBMC isolation.

Cells were then counted and divided into 12-well plates with approximately 2×10^6^ cells in each well. Cells were incubated overnight at 37°C in medium (IMDM (Biological Industries), supplemented with 10% of Heat Inactivated FBS (Rhenium, 10270106), Penicillin-Streptomycin-Amphotericin-B (Biological Industries, 1 03-033-1C)).

In the following day, to profile the immediate innate immune response, cells were either stimulated with LPS 100 ng/ml (Invivogen, tlrl-3pelps), 1 μg/ml high-molecular mass poly(I:C) (Invivogen, tlrl-pic), transfected with 2 μg/ml Lipofectamine 2000 (ThermoFisher, 11668027), and either mock-transfected or kept untreated. Following 4 hours of incubation, the stimulation was terminated by collecting the floating cells and trypsinizing (Trypsin 0.05%, no EDTA, Sartorius, 03-046-1B) the attached cells on the bottom of the well. Cells were then washed in 0.05% PBS. Next, cells were counted and loaded on the 10X Genomics Chromium instrument for single cell sequencing, aiming to capture 10,000 cells. We used either 10x Genomics CG000390 Rev B protocol Chromium Single Cell 3′ gene expression V.3.1 (10x Genomics) for CellPlex samples, or CG000204_ChromiumNextGEMSingleCell 3’ v3.1_Rev D protocol for the standard protocol. Libraries were sequenced on an Illumina NovaSeq 6000 using the NovaSeq 6000 SP Reagent Kit v1.5 (100 cycles) to generate 84-bp paired end reads.

## Computational analysis

### Mapping of single-cell RNA-seq data

*R. aegyptiacus* and mouse single-cell libraries from raw sequencing data were mapped to existing genomes and converted into UMI count matrices using 10x Genomics Cell Ranger v6.0^77^. Human data was downloaded from previous works of the Human Cell^30,42,75^. For *R. aegyptiacus* libraries, we used the bat1K genome assembly^78^, deposited at the NCBI website as mRouAeg1.p (https://www.ncbi.nlm.nih.gov/assembly/GCF_014176215.1). We tested it against a different genome annotation^19^, and chose the bat1K assembly since it gave a higher number of cells and allowed better identification and separation of cell types. Mouse libraries were mapped to ENSEMBL v92 mouse transcriptome (GRCm38.p6), the same ENSEMBL version as the version used in the human mapping. Cell Ranger filtered count matrices were used for downstream analysis.

### Manual annotation of *R. aegyptiacus* genes

The current *R. aegyptiacus* genome assembly contains genes with missing annotations (appearing as ‘LOC’). To augment the current assembly, we used gene orthology relationship with human and mouse genes, as inferred by our orthology mapping using the EggNOG mapper and its vertebrate database (v2.1.2) (see orthology mapping procedure below). In addition, we used online annotations found in the online NCBI website: https://www.ncbi.nlm.nih.gov/genome/gdv/browser/genome/?id=GCF_014176215.1 The above mentioned genes were missing from the downloaded version. Manually annotated genes by us appear in the dataset as “gene_LOCi” where “LOCi” is the original identifier and “gene” is the added manual annotation. In cases where there are more than one *R. aegyptiacus* gene matching to an EggNOG entry (i.e., paralogs likely originating from a recent duplication), we added a suffix of “_P”.

### Comparative analysis of bat, mouse and human cells Orthology mapping across species

The basis for most comparative analyses in this work is a set of one-to-one orthologs between human, mouse and *R. aegyptiacus.* Otherwise, in cases where we study gene duplicates, we specifically mention it. We used orthology relationships for the relevant species by running EggNOG 5.0 mapping^31^ using the *R. aegyptiacus*, human and mouse CDS annotations, with the assemblies mentioned above. Genes were considered one-to-one orthologs across the three species if they matched the same EggNOG gene name.

### Integrated UMAPs across species

Cross-species UMAPs were based on (1) expression of one-to-one orthologs across the three species, as defined above, and (2) cells belonging to the same annotated cell type shared across at least two species (i.e., cell populations that were not found in two of the three species were excluded from these integrated maps). In each such shared cell type, we downsampled the cells to achieve an equal number of cells per cell type across the species. For integration between mouse and *R. aegyptiacus* we also removed cells with over 10% mitochondrial reads from both species. We integrated the data using Seurat v4.1.1 integration pipeline. Each dataset was normalized using LogNoramalize. FindVariableFeatures was used for finding highly variable genes (**HVGs**) for the following stages. CCA and MNN algorithms(Hao et al. 2021) were then used for batch correction before integrating the datasets with IntegrateData. The integrated data was then scaled and PCA dimensionality reduction was calculated based on HVGs (npcs = 30). RunUMAP(dim = 1:30), FindNeighbors (dim = 1:30) and FindClusters(resolution = 0.5) were then used for clustering and visualizing the integrated data. The integrated datasets were then visualized using UMAP (using R’s DimPlot).

### Correlation between cell clusters across species

Cross-species comparisons were performed between *R. aegyptiacus* cells and human and mouse cells in each of the three tissues we studied. The cell types were compared in a pair-wise manner (*R. aegyptiacus*-human and *R. aegyptiacus*-mouse). In each case, gene expression in each cell cluster was aggregated to represent a ‘pseudo-bulk’ gene expression of this cluster. Spearman’s rank correlation between the expression levels of orthologous genes between cell clusters in *R. aegyptiacus* and the other species were then computed using differentially expressed (DE) genes, representing the marker genes that define each cell cluster and computed using Seurat v4.1.1^79^ FindMarkers. We compared either genes that are found to be DE genes between clusters in both species (the intersection of markers across the species), or the genes that are found to be DE genes between clusters in at least one of the species (the union of markers across the species). In each case we only used genes that are one-to-one orthologs between the two species. In each tissue, we show the correlation values of both the intersection and union of marker genes between human – *R. aegyptiacus* and mouse – *R. aegyptiacus*.

## Gut analysis

### Single-cell RNA-seq quality control and processing

For QC and processing of the single-cell data, we used Pandas (v.1.2.3), NumPy (v.1.20.2), Annadata (v.0.7.8), ScanPy^80^ (1.8.1) and Python (v.3.9.2). Genes were filtered to include those expressed in more than 3 cells. Cells were filtered for more than 200 expressed genes and for less than 50% mitochondrial reads, to match with previous processing of human gut single-cell data^30^. For doublet removal, we first used Scrublet (v.0.2.3)^81^ with a cut-off of 0.25. For further doublet exclusion during the downstream processing, we searched for unexpected co-expression of canonical markers from different cell types. For example, cells co-expressing CD3E/CD3D (markers of T cells) and EpCAM or other epithelial markers, were excluded from further analysis. Gene expression for each cell was normalized (sc.pp.normalize_total) and log-transformed (sc.pp.log1p). Cell cycle score was calculated using the expression of 97 cell cycle genes [Dissecting the multicellular ecosystem of metastatic melanoma by single-cell RNA-seq^82^]. The percentage of mitochondrial reads and unique molecular identifiers (**UMIs**) were regressed (sc.pp.regress_out) before scaling the data. These procedures were performed for *R. aegyptiacus* gut and for mouse small and large intestine.

### Clustering and cell type identification

Integration and batch correction of datasets were performed using bbknn (v.1.5.1, neighbors=2, metric=‘euclidean’, n_pcs=30, batch_key=’Sample’’). Dimensionality reduction and Leiden clustering (resolution 0.4–2) were performed, followed by cell lineages annotations. We used computationally derived marker gene expression for each cluster – DE genes (using sc.tl.rank_genes_groups, method=‘wilcoxon’). In addition, we searched for known markers to further annotate our cell clusters, using a set of genes taken from previous studies^30,42,83^. To identify subsets and cell populations in greater resolution, we subclustered the data by lineages, and again performed batch correction and Leiden clustering in an iterative fashion.

### R. aegyptiacus cells

Epithelial cells were subdivided, based on markers (in brackets), into stem cells (LGR5, RGMB, SLC12A2), transit-amplifying (**TA**; MKI67,TOP2A, PCNA), enterocytes (APOA1, ALPI, ANPEP), goblet (MUC2, REG4), Paneth (DEFA5, TFF3, GUCA2A), tuft (AVIL, TRPM5, POU2F3) and enteroendocrine cells (**EEC**; CHGA, CHGB). We also identified two separate clusters of enterocytes and goblet cells that we term PLK2, based on the expression of this gene in these two populations.

Mesenchymal cells were subdivided into fibroblasts (DCN, LUM, PDGFRA), pericytes (NOTCH3, PDGFRB) and smooth muscle cells (**SMC**; DES, ACTA2, TAGLN), while myofibroblasts were identified by the expression of ACTA2, TAGLN and PDGFRA, yet lacked the SMC marker desmin (DES). Fibroblasts were further divided into two populations by the expression of either PI16 or ADAMDEC1. In addition, we identified glial cells (MPZ, CDH19, PLP1), and mesothelial cells on the basis of LRRN4 and PRG4.

Endothelial cells (PECAM1) were subdivided into vascular on the basis of VWF or lymphatic (**LEC**) on the basis of CCL21, PROX1 and LYVE1.

Myeloid cells were subdivided into monocytes (F13A1, IL1B), macrophages (C1QA, CD163) and dendritic cells. Dendritic cells (**DCs**) were defined as either cDC1 (CLEC9A, XCR1) or cDC2 (IL18, SIRPA), both expressing BATF3 and HLA-DRA, or as pDCs, on the basis of IRF8 and TCF4. Neutrophils (CSF3R, CCRL2, G0S2), Mast cells (GATA2, IL1RL1, MS4A2) and Eosinophils (PRG3, RFLNB, CD24) were identified as well, comprising the granulocytes group.

Lymphocyte cells were subdivided into T/NK cells and B cells. The T cell (CD3D) compartment is composed of CD4 T cells, expressing CD4 and not CD8A, and vice versa for CD8 T cells, as well as an additional T CD8 subset, expressing GZMK. Natural killer (**NK**) cells were identified based on the expression of NCR1 and KIR receptors (KLRB1 and KLRD1), yet lacking the expression of CD3D. Natural killer T (**NKT**) cells were identified by the expression of the KIR receptors together with CD3D and the lack of NCR1. Innate lymphoid cells (**ILCs**; TRDC) were defined as ILC3 (IL23R, KIT, RORC, FCER1G [An *in vitro* model of innate lymphoid cell function and differentiation]) or ILC2 based on the lack of ILC3 markers combined with a stronger expression of GATA3 and RORA. Cycling T cells were also identified, expressing CD3D together with STMN1. B cells were identified by the expression of CD79B together with MS4A1 while plasma B cells were expressing MZB1 as well. Cycling B and Plasma cells were identified by the additional expression of STMN1.

### Mouse cells

We used the same markers as above, with the following differences:

In SI: SCs – we also used OLFM4, Enterocytes – ALDOB, RBP2, goblet cells – TFF3, SPINK4, Paneth – DEFA24, LYZ1, Tuft – DCLK1, Mesenchymal – CD34, COL6A2. In the immune compartment, additional markers were used for macrophages – LYZ2, H2-EB1, cDC2 – CD209A, pDC – SIGLECH, BST2. In lymphocytes, CD28 was used for CD4 T cells and JCHAIN for B plasma cells.

In colon: In addition to the mouse SI markers and the *R. aegyptiacus* gut markers: Colonocytes were defined by the expression of CAR1, CAR4 and AQP8 and vascular endothelial cells by the additional marker SOX17. In the immune compartment, we also used the marker CCR2 for Neutrophils, for activated DC – CCR7 and FSCN1, and for CD4 T cells – LEF1.

### Spatial transcriptomics mapping, QC and spatial abundance of cell type analysis

BCL files were converted to fastq format and demultiplexed using 10x Genomics Space Ranger 2.0.0 (https://support.10xgenomics.com). The converted fastq files were then mapped to the reference *R. aegyptiacus* transcriptome (the same as with the single-cell RNA-seq data) using Space Ranger. Next, we spatially mapped gut cell types using our *R. aegyptiacus* spatial transcriptomics data, integrating scRNA-seq using Cell2location (v0.1)^32^. Only Visium spots aligned on the tissue were used for analysis, exclusion of spots was performed with Loupe Browser v6.3.0. We employed the Cell2location model to estimate the abundance of each cell population in each spot. This model leverages the transcriptional signatures of reference cell types to decompose the mRNA counts in the 10x Genomics Visium data. The gene expression matrix was filtered to remove lowly expressed genes using filter_genes with default values. Cell2location was employed to estimate reference gene expression signatures of cell types from using negative binomial regression. For the processing of the 10x Visium data, we adjusted the hyperparameters in Cell2location as follows: the model was trained for 30,000 iterations; The RNA detection sensitivity, detection alpha, was set to 20 and the number of cells per location to 18, based on comparison with histology images paired with 10x Genomics Visium. In this analysis we used libraries from Scanpy (v1.8.1)^80^, Anndata (v0.8.0) and Matplotlib (v3.6.2).

For mouse SI spatial transcriptomics data, we used data from a previous study^83^. We processed the mouse data as described above.

### Transcriptional analysis of small intestine versus colon analysis

To establish a set of SI enterocyte and colonocyte signature genes, a DE analysis was performed between the two cell types in human and, separately, on mouse, using 1:1 orthologs across human, mouse and bat. Enterocytes and colonocytes from mouse and human single-cell data (healthy adult samples) were down-sampled for equal cell numbers in each species (python random package with sample method, seed =1), 4,271 of each type in human and 4,456 in mouse. Human enterocytes of duodenal and rectal origins were excluded, as well as pediatric samples. sc.tl.rank_genes_groups (method=‘wilcoxon’) was then used for DE analysis between enterocytes and colonocytes in each species. Genes were filtered for 0.01 adjusted p-value; mitochondrial genes were excluded from the analysis. From the DE results, we obtained 100 signature genes that are highly expressed in enterocytes versus colonocytes and vice versa. This was done by taking the genes with the lowest “Max Q-value” (the higher adjusted p-value of a gene in the enterocytes-colonocytes DE analysis in human and mouse).

For heatmap representation of enterocyte and colonocyte genes in different gut sections and in different species, we downsampled the cells (human and mouse SI enterocytes and colonocytes and *R. aegyptiacus* enterocytes) to reach an equal number of 4,280 cells. Data was then normalized (sc.pp.normalize_total) and log-transformed (sc.pp.log1p) following regression of mitochondrial reads and unique molecular identifiers (UMIs) (sc.pp.regress_out) before scaling the data (sc.pp.scale). The data was plotted using sc.pl.heatmap. To test for relative gene expression of enterocyte and colonocyte signature genes in *R. aegyptiacus* enterocytes*, we used* a one-sided Mann-Whitney U test (scipy.stats.mannwhitneyu).

## Evolutionary analysis of enterocyte trans-differentiation along the villus

### Top and bottom villus scores

To identify the likely zonation of enterocytes in our single-cell and spatial transcriptomics data, we used 5 sets of landmark genes, highly expressed in specific regions along the villus, from a previous work that identified them in mouse villi using laser capture microdissection of five sequential regions along the villus and bulk-RNAseq^37^. We then used only those genes that were identified as one-to-one orthologs across human, mouse and bat. The scoring was done using sc.tl.score_genes() function with default parameters to calculate the average expression of selected genes subtracted with the average expression of a reference set of genes. The score was then standardized between 0 and 1. To visualize these scores in the spatial data, we used sc.pl.spatial.

### Inferences of enterocyte zonation along the villus

Following the scoring, enterocyte single-cell datasets from each of the three species were filtered for genes expressed in more than 3 enterocytes, and then normalized (sc.pp.normalize_total) and log-transformed (sc.pp.log1p), following by regressing out mitochondrial reads and unique molecular identifiers (UMIs) (sc.pp.regress_out). Log-normalized counts of highly variable genes (HVGs) were scaled (sc.pp.scale) before performing principal component analysis (PCA). We then plotted the 5 zonation scores in human,mouse and bat enterocytes. Since the scores largely followed PC1, we used PC1 coordinates to define enterocyte zonation, by dividing the cells into 5 sequential sections along PC1. For top versus bottom comparison, we further used 15% of the enterocytes found in the two extremes of the axis.

### scVelo trajectory analysis

The cell trajectory analysis for the bat and mouse enterocytes was carried out with scVelo 0.2.4^38^ package, as implemented in scanpy. Pre-processing of the enterocyte data was performed with the function scv.pp.filter_and_normalize(min_shared_counts=20), followed by scv.pp.moments function. Gene-specific velocities were determined using scv.tl.velocity(mode = ‘stochastic’) and scv.tl.velocity_graph(), by quantifying the steady-state equilibrium deviation from a ratio between unspliced to spliced mRNA levels. Gene-specific velocities were visualized with scv.pl.velocity_embedding_stream(basis=’pca’).

In order to plot heatmaps of expression along the villus, a dynamic model pipeline was used with the functions scv.tl.recover_dynamics() followed by scv.tl.velocity(mode=’dynamic57al’) and scv.tl.velocity_graph(). Heatmaps were plotted using scv.pl.heatmap for the 5 sets of gene signatures along the sections of the villus used for the enterocyte scores. Enterocytes were sorted by the PC1 coordinates used to identify the regions along the villus axis from which they likely originated.

### Differential expression analysis of top and bottom villi enterocytes

Top and bottom enterocytes were downsampled (python random package with sample method, seed =1) for equal cell numbers between regions and between species resulting in a total number of 627 enterocytes in each group. Downstream processing was performed in a similar manner as above (“Inferences of enterocyte zonation along the villus**”**), with the addition of a batch correction performed with bbknn (v.1.5.1, neighbors=2, metric=‘euclidean’, n_pcs=30, batch_key=’Sample’ or ‘sample name’), and a UMAP dimensional reduction. DE analysis was carried out using sc.tl.rank_genes_groups (method=‘wilcoxon’) for each species individually between the top and bottom regions. Genes were annotated as top (positive logFC) or bottom (negative logFC) in each species separately. In addition, genes were labeled based on expression levels as: not expressed, expressed in more or less than 5% of the population, or expressed in more than 20% of the population.

### Gene age

For the gene age analysis, we used genes expressed in more than 20% of the human enterocytes and based the analysis on the human DE analysis. We used human, as gene age estimations are available and best characterized for the human genome (although these one-to-one orthologous genes should not vary between species). For this, human enterocyte single-cell data was filtered for genes expressed in over 20% of the top or bottom enterocyte population by sc.pp.filter_genes(human, min_cells=int(0.2*human.shape[0])), then processed in a similar manner as for the DE analysis of the three species. 500 top (positive logFC) and 500 bottom (negative logFC) most significant genes were selected for the analysis by lowest adjusted p value. Gene age estimations according to three different methods and family reconstruction (Wagner/Ensembl, Wagner/OrthoMCL and Dollo/OrthoMCL) were obtained from ProteinHistorian^39^. For each gene, age was defined with respect to the species tree, where a gene’s age corresponds to the branch in which its family is estimated to have appeared (thus, larger numbers indicate evolutionarily older genes). Data was plotted using Seaborn package boxplot, for statistics we used a one-sided Mann-Whitney U test (scipy.stats.mannwhitneyu).

### Transcriptional conservation of top and bottom genes

To study the transcriptional conservation of the sets of top and bottom genes, we compared DE results between top and bottom genes in human data with those in mouse and *R. aegyptiacus*. Human DE results were selected as the reference for comparison since its DE genes were the most significant (top versus bottom genes) and since it was also used in the gene age estimates. For comparison across species, we used two unbiased sets of top and bottom genes, where we controlled for the adjusted p-values, such that both sets would have a similar distribution of these p-values. For this, we used a function we developed previously^84^: row_matchers.one_to_one_matches(col = –logQval_human, eps = 5). We used this script with initial sets of 250 most significant genes from each of the bottom and top gene sets. This method yielded 70 human-paired DE genes (70 top and 70 bottom genes). Top and bottom genes were then projected on the bat and mouse DE data, to test the distributions of adjusted p-values of these two gene sets in each of these species. Data was visualized and statistical significance was determined as described above (“gene age”) and the statistics as well, except for the Mann-Whitney U test performed for the human DE paired genes that was done as a two-sided test in order to check for biases in both directions. For volcano plot visualization, we used seaborn scatterplot with the DE analysis data.

### Species-specific top gene analysis

In this analysis we first searched for genes that are transcriptionally divergent such that they are either expressed as top villus genes only in the bat cells (in comparison with both human and mouse) or the opposite (i.e. – top genes in both human and mouse, but not in bat). We defined genes to be expressed as top in bat but not in human and mouse as follows: We first kept genes that are top in bat cells with an adjusted p-value<0.01 and that their logFC in the other two species is below 0.5. We also asked that the difference between the logFC values of bat versus either human or mouse will be positive. We then took the top 100 genes that are most significant in terms of having the highest differences of logFC in bats in comparison witheither species.

For genes that are top in human and mouse but not in bat, we performed a similar procedure, where we used genes that (1) had an adjusted p-value<0.01 in both human and mouse cells, (2) their logFC in bat cells was below 0.5 and (3) the differences between the logFC values of bat versus both human or mouse was negative. In each set, we searched for significantly enriched pathways and functions using g:Profiler^43^ with the top 100 genes.

### Expression analysis along the villus of specific sets of genes

In this analysis, we used one-to-one orthologous genes, found in gene sets with known functions. These sets were either the result of the previous analysis (see above – the two sets were those related to the terms: (1) Lipid transport GO:0006869, and (2) Anion binding GO:0043168) or involved in nutrient absorption and transport and taken from a previous study(doi.org/10.1084/jem.20191130). We only used genes expressed in each of the species in more than 10% of the enterocytes in at least one of the 5 regions along the villus. Expression was then visualized by sc.pl.matrixplot(standard_scale = ‘var’) across the villi axis with the groups V1-V5.

### Differential expression analysis of gut cells across species

DE analysis was performed between homologous cell types in *R. aegyptiacus gut* and human or mouse SI as integrated Seurat objects using FindMarkers() of Seurat v3 with default parameters (min.pct=10, logfc.threshold = 0.25, assay= “RNA”). This analysis included only those genes that were identified as one-to-one orthologs across human, mouse and *R. aegyptiacus* and equivalent cell types were down-sampled (python random package with sample method, seed =1) for equal cell numbers between the *R. aegyptiacus* and the other species. We later searched for significantly enriched pathways and functions using g:Profiler^43^ with the top 50 significant genes (adjusted p value< 0.01) with the highest logFC (see enriched pathways for R. *aegyptiacus* stem cells in **Table S2**).

The complement system gene set used in the analysis was taken from a previous work^44^ and is composed of pattern recognition, proteases, complement components, receptors and regulators from all three major pathways. In order to visualize the DE results regarding complement system proteases and components we used seaborn’s heatmap with –log(adjusted P value) values, and set a maximum threshold of 3. Only significant DE genes (default FindMarkers() parameters) in at least one cell type in either *R. aegyptiacus* –mouse DE or *R. aegyptiacus* –human DE analysis were included. For visualization of the expression of the complement system genes in all three species, we used Seurat’s (VlnPlot()) and dotplot(DotPlot()), with normalized log-transformed single cell data.

For volcano plot visualization, we used seaborn’s scatterplot with DE analysis data of all orthologs shared between the three species using Seurat v3’ FindMarkers with no threshold. The set of innate immune genes was obtained from InnateDB^41^.

### Spatial analysis of complement gene expression

Spatial complement gene expression analysis was carried out using ScanPy in the following manner: Spatial transcriptomics data was normalized, log-transformed and scaled, similarly to the single-cell data analysis. Next, dimension reduction was performed followed by Leiden clustering (resolution= 1.4). The clusters were visualized by UMAP and on the spatial grid. Selected clusters were then annotated for either the intestinal wall, the crypt or the villus, based on markers of the cells residing in these areas as follows: Wall: SMC – DES, Crypt: Paneth cells – DEFA5 and STMN1 (a marker expressed in cycling and differentiating cells), Villus: Enterocytes – APOA1. Following annotation, DE analyses were performed using rank_genes_groups (method= ‘wilcoxon’, group_by = ‘area’) and selected complement components and proteases gene expression were visualized on the spatial grid or using ScanPy’s dotplot.

In order to visualize the expression of the complement components and proteases along the villus, crypt cells (TA, Stem and Paneth cells) and enterocytes were plotted for expression using ScanPy’s matrix plot (sacpny.pl.matrixplot()) as was done in the top and bottom analysis section with single-cell data.

## Lung analysis

### Single-cell RNA-seq quality control, clustering and cell type identification

For QC and clustering of *R. aegyptiacus* and mouse lung single-cell data, we employed a similar approach to the approach described above for gut cells, with the exception of using a threshold of cells with fewer than 30% mitochondrial reads.

### Clustering and cell type identification

*R. aegyptiacus* data were integrated and Leiden-clustered as described in the gut section. Clusters and subclustres were identified using computationally derived marker genes between clusters. We also used known markers from previous studies, as follows:

### R. aegyptiacus cells

Epithelial cells were subdivided into Alveolar type I (**AT1**; AGER, CLIC5, RTKN2), Alveolar type I (**AT2**; SFTPC, SFTPB, MUC1), basal (KRT5, TP63, KRT17), ciliated (FOXJ1, DYNLRB2), club (SCGB3A2) and goblet cells (TFF3, MUC5AC, SPDEFF and FOXA3). Tuft cells, also known as brush cells, were identified on the basis of BIK, PLCG2, ALOX5AP and SOX9^85^.

Endothelial cells (PECAM1) were subdivided into vascular (VWF) and lymphatic (**LEC**; CCL21, MMRN1, PROX1) cells. The vascular endothelial cells were further divided into capillary (EDNRB, CA4, TBX2, PRX), venous (ACKR1, VWF) and arterial on the basis of EFNB2, SEMA3G and VWF^86^.

Mesenchymal cells were subdivided into fibroblasts (DCN, LUM, PDGFRA), pericytes (NOTCH3, PDGFRB) and smooth muscle cells (**SMCs**) based on the expression of ACTA2, MYH11 and DES. Fibroblasts were further divided into alveolar (COL13A1, NPNT), adventitial (PI16, COL1A1) and peribronchial (HHIP, FGF18, WIF1)^87^. Pericytes were divided into two populations based on the expression of ACTA2 in one of the subsets. In addition, we identified mesothelial cells on the basis of LRRN4, PRG4 and MSLN.

Myeloid cells were subdivided into CD14 monocytes (CD14, F13A1, VCAN), CD16 monocytes (FCGR3A, CX3CR1), macrophages (CD163) and dendritic cells (**DCs**). Macrophages were divided into alveolar macrophages, on the basis of MARCO, and to macrophages interstitial, based on a stronger expression of C1QA combined with a lack of MARCO expression. DCs were defined as either cDC1 (CLEC9A, XCR1), cDC2 (IL18), or activated DC (LAMP3, FSCN1, CCR7), all expressing HLA-DRA as well as BATF3, and pDCs (IRF8 and TCF4). Neutrophils (CSF3R, CCRL2, G0S2, S100A8), mast cells (GATA2, IL1RL1, MS4A2) and eosinophils (PRG3, RFLNB, CD24) were identified as well, comprising the granulocytes group. Neutrophils were further subdivided into subtypes based on the expression of OLFM4^88^.

Lymphocytes were subdivided into T/NK cells and B cells. The T/NK population was subdivided into T (CD3) CD4 or CD8 expressing cells, innate lymphoid cells (**ILCs**) and NK cells. The CD4-expressing T cells were further divided into naïve T cells on the basis of CCR7, TCF7 and LEF1, central memory CD4 (**CM**) T cells, a population with a weaker expression of these genes and effector memory (**EM**) CD4 T cells, on the basis of CXCR6 and CCL5. The CD8-expressing T cells were divided into CM and EM CD8 T on the basis of additional expression of GZMB and TBX21^89^, and NKT. NKT and NK cells were identified based on the expression of the KIR receptors KLRB1, KLRD1 and NCR1, and distinguished based on expression of CD3D and CD3G (NKT), or lack of such expression (NK cells). ILCs (TRDC) were defined as ILC2, based on GATA3, MAF and IL1RL1 expression, or as ILC3, based on RORC, IL23R, KIT and FCER1G expression^90^. Cycling T cells were identified by the expression of CD3D together with STMN1. B cells were identified by the expression of CD79B together with MS4A1, while B plasma cells were expressing MZB1 and JCHAIN. Cycling B and cycling B Plasma cells were defined based on the additional expression of STMN1.

Erythrocytes were also identified, using HBB and ALAS2 expression.

### Mouse cells

We used the same markers as above, with the following differences:

For goblet cells we used the marker MUC5B^91^. For arterial endothelial cells, we used SOX17^86^, and for the peribronchial fibroblasts we used ASPN as well^87^.

In the immune compartment we used additional markers as follows: Mast cells – CPA3, classical monocytes (LY6C^+^) – CCR2, non-classical monocytes (LY6C^-^) – ENO3^92^, alveolar macrophages – LPL, cDC2 – CD209A, activated DC – BIRC3, pDCs – BST2, SIGLECH, and regulatory T cells (**Treg**) – FOXP3 and CTLA4.

### Differential expression analysis of lung cells across species

The same procedures as in DE analysis of gut cells across species were performed for the lung-resident cells. See Go term analysis and enriched pathways for *R. aegyptiacus* AT2 cells in **Table S7**.

### Detection of Positive Selection in Complement System Genes in Bats

We employed a phylogenetic-based approach to analyze 14 genes involved in the classical and alternative pathways of the complement (C1S, C1R, C2, CFB, CFD, CFI, C3, C5, C6, C7, C9, C8A, C8B, and C8G). We used orthologous genes from up to 10 bat species, representing different branches across Chiroptera (*Rousettus aegyptiacus*, *Pteropus vampirus*, *Rhinolophus ferrumequinum*, *Molossus molossus*, *Phylostomus discolor, Artibeus jamaicensis*, *Pteropus_vampirus*, *Pipistrellus kuhlii*, *Myotis myotis*, *Myotis lucifugus*). Orthologous sequences were aligned using MAFFT^93^ with default parameters, and poorly aligned regions were removed (by columns) using GUIDANCE v2.0.2^94^ (default parameters). We employed PhyML v3.1^95^ for phylogenetic tree reconstruction for each of the MSAs. Finally, we used codeML from the PAML package v4.9h^45^ using a site model (NSsites = 8, model = 0). In order to evaluate the null hypothesis against the alternative model, we employed the likelihood-ratio test (**LRT**) and calculated the corresponding p-value using a chi-square distribution with 1 degree of freedom, followed by FDR correction. Positively selected genes were considered those with a corrected p-value<0.05 (8 out 14).To identify sites under positive selection, we utilized the Bayes Empirical Bayes (**BEB**) approach. Genes with residues having over 95% probability for positive selection (Pr > 0.95) were considered to exhibit strong evidence of positive selection.

## PBMC analysis

### Single-cell RNA-seq quality control, clustering and cell type identification

For QC and clustering of *R. aegyptiacus* PBMC single-cell data, we employed a similar approach to the one described above for gut cells, with the exception of using a threshold of cells with less than 20% mitochondrial reads. *R. aegyptiacus* data were integrated and Leiden-clustered as described in the gut section. Clusters and subclustres were identified using computationally derived marker genes between clusters as well as using known markers from previous studies^30,42,89,96–98^.

### R. aegyptiacus cells

Myeloid cells were subdivided into monocytes (MAFB), cDCs (IL18,CD1A)^99^, activated DC (LAMP3, FSCN1, BATF3, CCR7) and pDCs on the basis of IRF8 and TCF4. Monocytes were further divided into CD14 monocytes (CD14, F13A1, VCAN)^92^, CD16 monocytes (FCGR3A, CX3CR1) and a population defined by the expression of IFNB1. Neutrophils were identified as well, based on CSF3R, CCRL2 and G0S2.

The lymphocytes were subdivided into T/NK cells and B cells. The T/NK population was subdivided into T (CD3) CD4 or CD8 expressing cells and NK cells. The CD4 expressing T cells were further divided into naïve T cells on the basis of CCR7, TCF7 and LEF1, central memory CD4 (CM) T cells, a population with a weaker expression of these genes and effector memory (EM) CD4 T cells, on the basis of CXCR6. Another CD4-expressing population is the regulatory T cells (Tregs), identified by the markers FOXP3 and CTLA4. The CD8-expressing T cells were divided into central memory and effector memory CD8 T cells on the basis of additional expression of GZMB and a lack of the differentiation marker CD7^98^. NKT and NK cells were identified based on the expression of the KIR receptor KLRD1 together with NCR1, while NK cells lacked the expression of CD3D and CD3G. In a similar manner to CD8 T cells, NKT and NK cells were identified as activated when lacking the expression of CD7 while expressing GZMB.

B cells were identified by the expression of CD79B together with MS4A1 while plasma cells were expressing MZB1 and JCHAIN as well. Another B cell population was annotated based on the additional expression of the activation marker CD83.

### Mouse cells

Mouse datasets were integrated with Seurat CCA and annotations were done as described for *R. aegyptiacus* with the following differences:

For the Neutrophils, we used the S100A8 marker as well, for the LY6C2+ monocytes we used the canonical marker LY6C2, for the LY6C2-monocytes – APOE and FCGT4^100^,for the pDCs – BST2 and for the NKT cells we used NKG7 as well.

### Comparative analysis of expression of specific sets of immune genes across cell types

To compare gene expression of specific gene sets of interest, we created scores using Scanpy’s tl.score_genes() with default parameters. For each cell, a score is the average expression of selected genes subtracted from the average expression of all genes. The score is then standardized to be between 0 and 1. We used three scores representing different stages and pathways known to be upregulated during the innate immune response against pathogens: (1) Inflammatory score: A set of inflammatory response-related and inflammatory cytokine and chemokine genes^47^; (2) Primary antiviral response score: Primary antiviral response genes were taken from a set of genes upregulated following 4h of dsRNA-stimulation in dermal fibroblasts taken from previous studies^20,46^. The set we used here includes 175 genes that were upregulated in all studied species – *R. aegyptiacus*, mouse and human. (3) Interferon response score: 79 Interferon-stimulated genes (ISGs), representing the second wave of response, were taken from a list of ISGs known to be conserved across mammals^48^.

### Differential expression analysis of LPS and dsRNA stimulation in monocytes

DE analysis of monocyte stimulation was performed using FindMarkers() of Seurat v3 with default parameters (min.pct=10, logfc.threshold = 0.25, assay= “RNA”). We compared LPS-stimulated cells to unstimulated cells, and dsRNA-transfected cells to unstimulated cells.

For gene enrichment analysis we used DE genes of stimulated (LPS and dsRNA) and unstimulated bat monocytes. The top 100 genes with the lowest adjusted P-values for each condition were tested for functional enrichment using g:Profiler^43^. We then filtered for terms unique to each condition and plotted these using Python’s seaborn barplot.

For producing the numbers of DE genes between different cell types in different conditions, as appears in Table S11, we performed downsampling of the data prior to DE calculations. All cell types in an Unstimulated-Stimulated DE calculation were downsampled to having the same number of cells as the cell type with the smallest number of cells. Similar procedures of downsasmpling were performed in all DE analysis between species, and/or across cell types and/or conditions (e.g. Table S14).

## Analysis of dsRNA-response in in *R. aegyptiacus* and *mouse* monocytes

### Characterization of 3 states in monocytes during dsRNA stimulation

We divided CD16 monocytes cells into 3 groups as follows: monocytes were found in two distinct clusters in the UMAP: a cluster with almost exclusively dsRNA-stimulated cells, enriched with IFNB1 expressing cells (termed ‘**IFN monocytes**’), and a separate cluster composed of both unstimulated, mock-stimulated and dsRNA-stimulated cells. In the latter cluster, we divided the cells based on those that are dsRNA-stimulated and all others, resulting in a total of three cell sets. In this analysis, we excluded cells from one particular individual that were outliers in the UMAP. We then used DE analysis (using the same functions and parameters as above) to obtain the top upregulated genes in each cell group, by comparing each group of cells to the other two and taking the 100 most significantly upregulated (logFC > 0 and Q-value < 0.01).The expression of the expression of the resulting 300 genes across all CD16 monocyte cells were then clustered and plotted using Python’s seaborn.clustermap(). Both genes and cells were clustered using the “ward” clustering method. We then performed GO term analysis on each of the gene groups using g:Profiler^43^.

### Conservation of the IFN monocytes DE gene between bat and mouse

We next defined the groups of *R. aegyptiacus* genes that are uniquely up– and down-regulated in IFN monocytes in comparison with the other two groups (dsRNA monocytes and unstimulated monocytes). This was based on the DE analysis between the three groups described in the previous section. We define 59 genes to be upregulated only in IFN monocytes as those (1) having logFC > 0 and Q-value < 0.05 in the IFN monocytes in comparison with the two other groups as well as (2) not upregulated in other dsRNA-stimulated monocytes. Similarly, we define 725 genes to be uniquely downregulated in IFN monocytes with respect to the other groups of monocytes.

### Analysis of recent IFN gene duplicates

Type-I IFN genes in *R. aegyptiacus* genome were detected through their orthology assignments in EggNOG. Log-normalized expression levels of type-I IFN genes that were expressed in *R. aegyptiacus* and mouse cells (IFNB1 and various IFNA and IFNW genes) were plotted in different cell groups in mouse and *R. aegyptiacus* cells using Scanpy’s dotplot(). As the expression was almost entirely in the group of IFN monocytes, violin plots of Log-normalized expression of IFN genes were generated using Python’s seaborn.violinplot() function using data from this group of cells.

## Data availability

Raw sequencing data and processed count matrices with metadata will become available at ArrayExpress (https://www.ebi.ac.uk/biostudies/arrayexpress/studies) upon publication. Previously published human data is detailed in the relevant sections and can be downloaded from ArrayExpress with the accession numbers E-MTAB-9543, E-MTAB-9536, E-MTAB-9532, E-MTAB-9533 and E-MTAB-10386.

## Code availability

Code generated during this study is available at: https://github.com/HagaiLab/bat_immunty_evolution.

## Acknowledgements

We would like to thank Amit Zeisel, Osnat Hadad Ophir, Muhammad Tibi, Xi Chen, Matthias Friedrich, Roser Vento-Tormo, Luz Garcia-Alonso and Naama Peshes-Yaloz for helpful discussions during the project and on the manuscript. We would like to thank Stefan Kaltenbach and Rami Khosravi for technical assistance with NGS library preparations. This research was supported by the Israel Science Foundation (ISF, grant No. 435/20), the Chan Zuckerberg Initiative (Single-Cell Analysis of Inflammation, grant No. DAF2020-217532), by a joint QBI/UCSF-TAU research grant in computational biology and drug discovery. GD was supported by the Chan Zuckerberg Initiative (Single-Cell Analysis of Inflammation, Id. DAF2020-217532) and by AIRC, Associazione Italiana per la Ricerca sul Cancro (MFAG 2018 – Id. 21640).

