## Supplemental_Figures for "Spatial and single-cell transcriptomics illuminate bat immunity and barrier tissue evolution"

### Supplemental Data

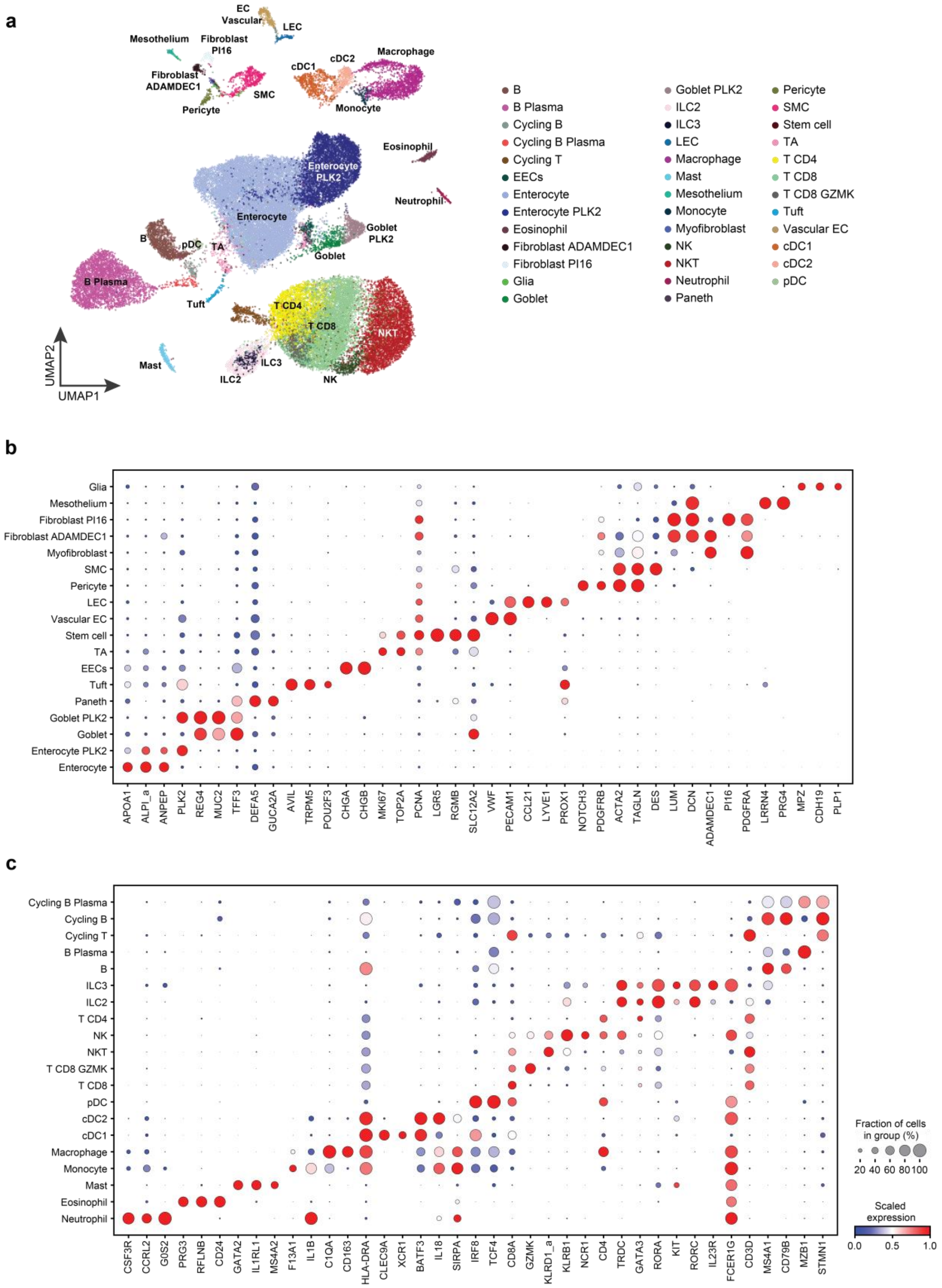


**Supp Data Fig. 1: *R. aegyptiacus* gut cell subsets defined the study. (a)** Uniform manifold approximation and projection (UMAP) of *R. aegyptiacus* gut cell clusters. **(b-c)** Dot plots for expression of marker genes of **(b)** non- immune and **(c)** immune cell types and states in the scRNA-seq dataset.

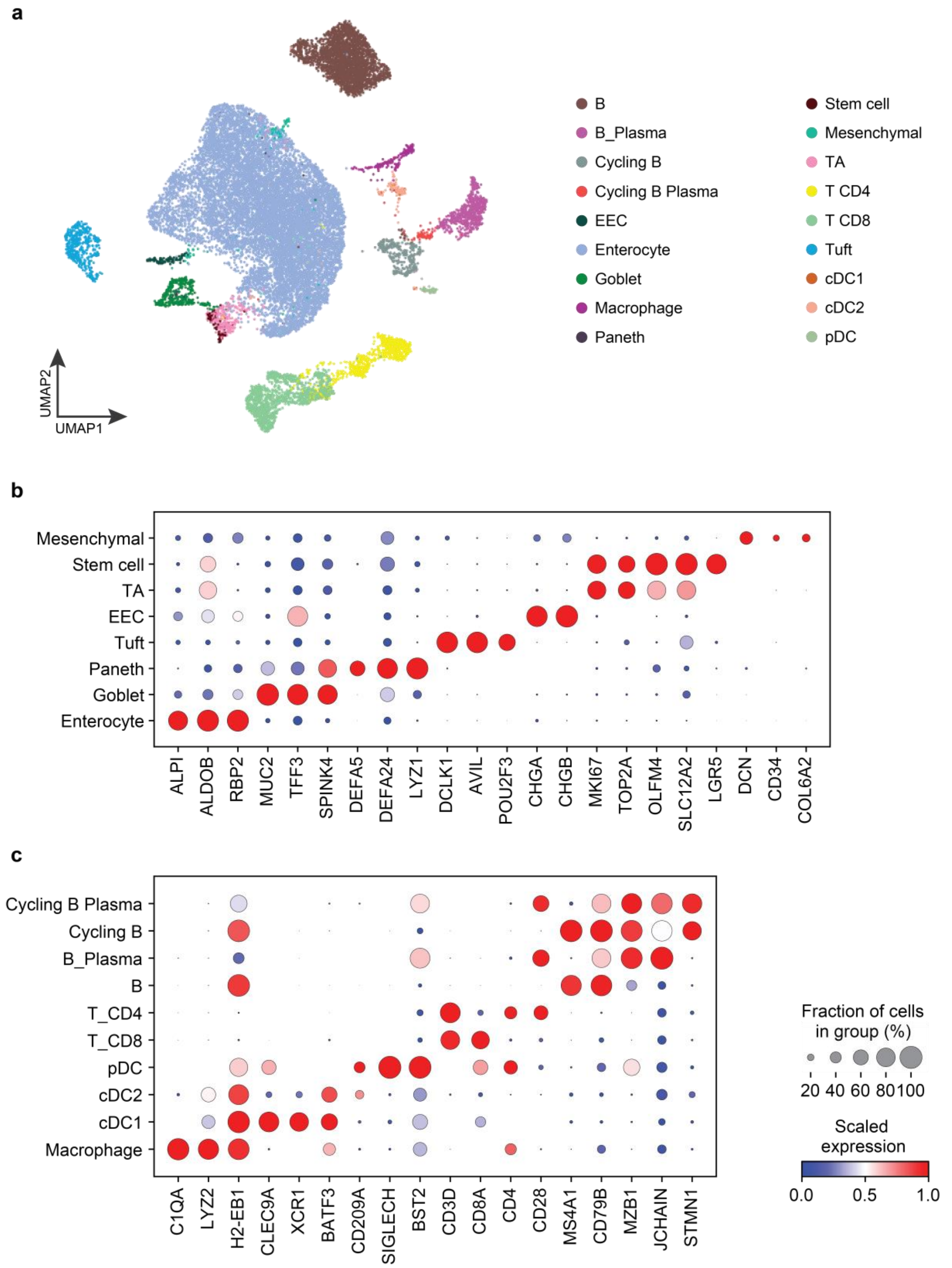


**Supp Data Fig. 2: Mouse small intestine characterization and cell subsets defined the study. (a)** Uniform manifold approximation and projection (UMAP) of mouse small intestinal cell clusters (5 samples, n= 20346 cells). **(b-c)** Dot plots for expression of marker genes of **(b)** non- immune and **(c)** immune cell types and states in the scRNA-seq dataset.

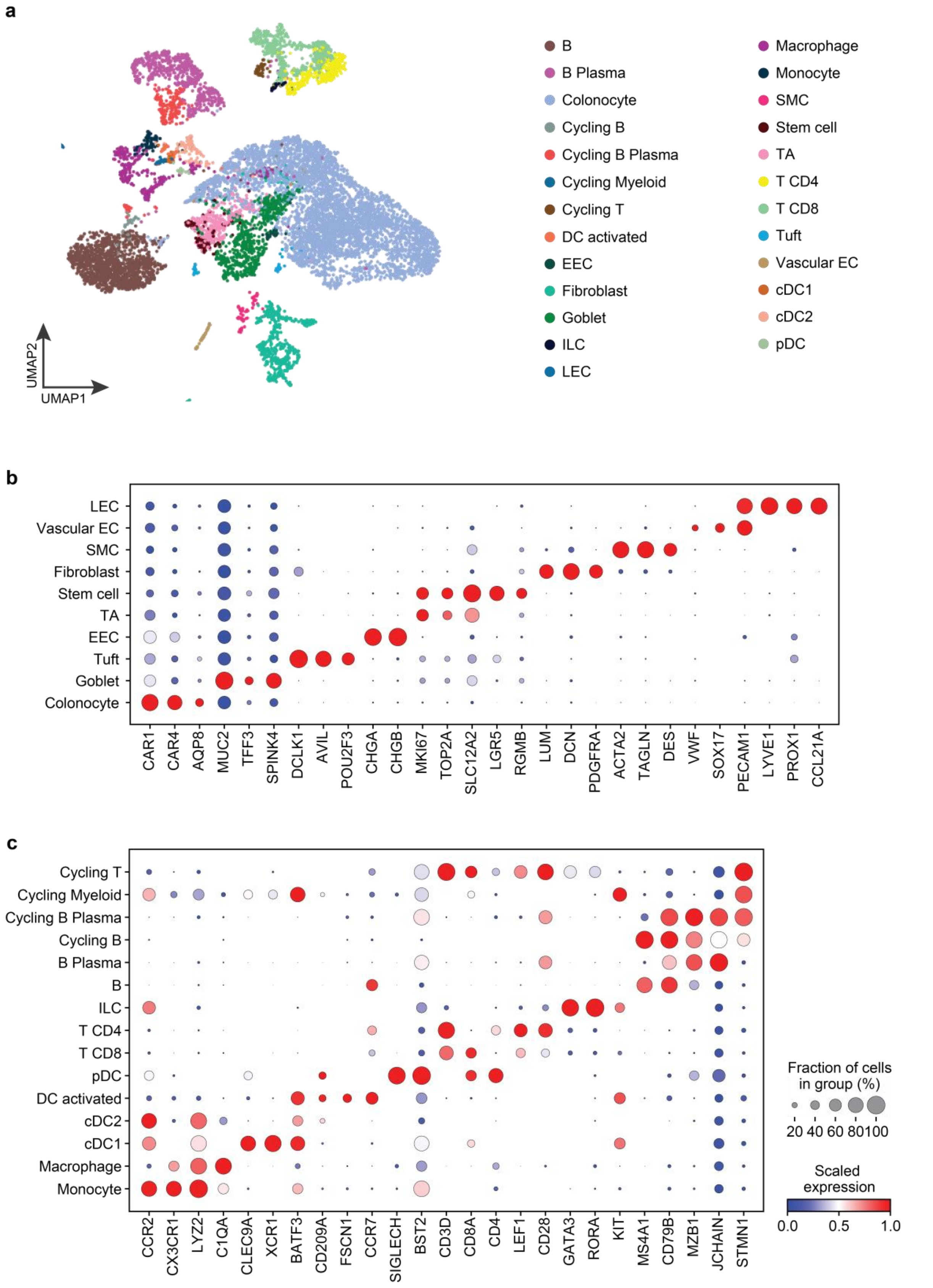


**Supp Data Fig. 3: Mouse large intestine characterization and cell subsets defined the study. (a)** Uniform manifold approximation and projection (UMAP) of mouse small intestinal cell clusters (2 samples, n= 10070 cells). **(b-c)** Dot plots for expression of marker genes of **(b)** non- immune and **(c)** immune cell types and states in the scRNA-seq dataset.

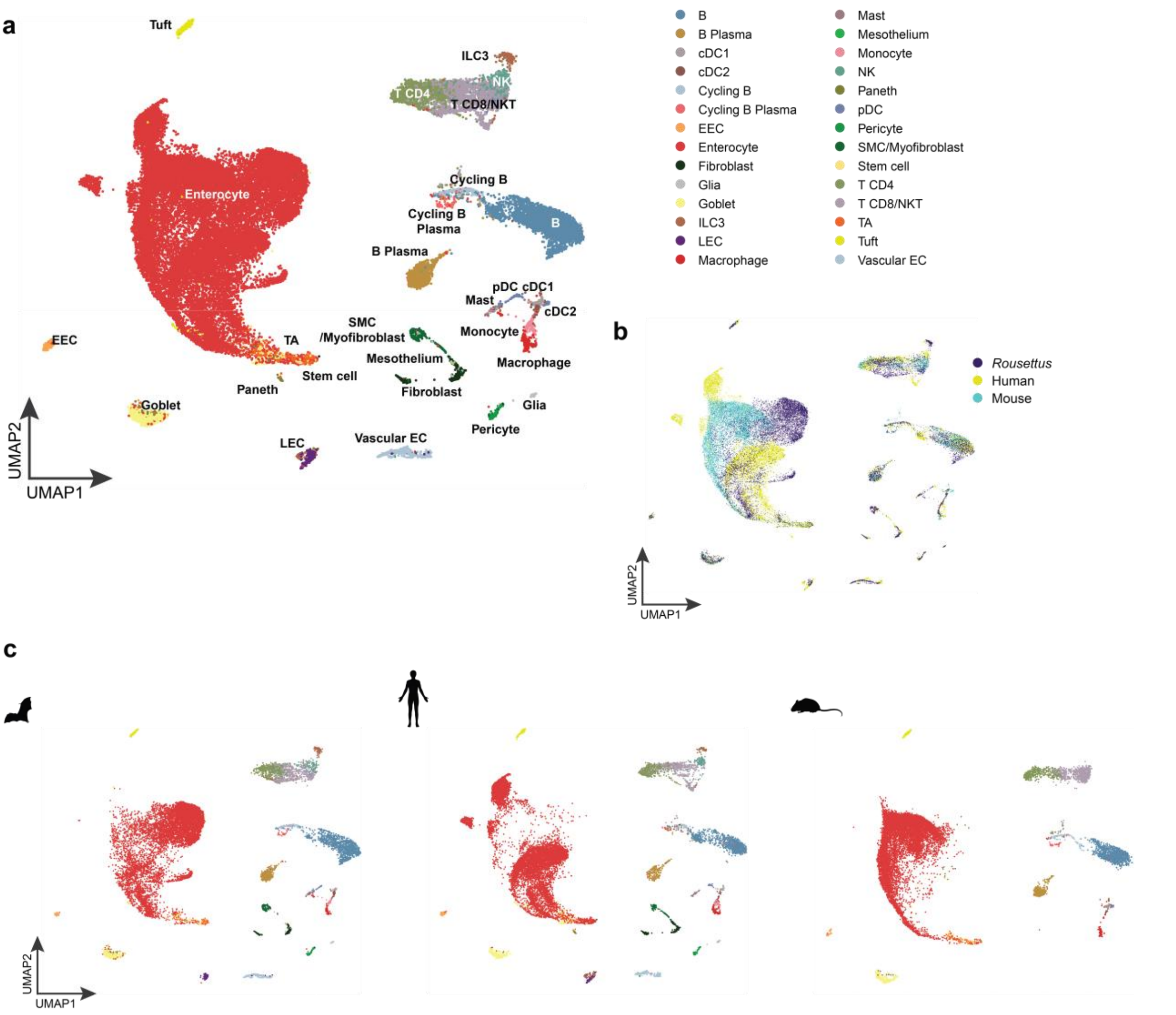


**Supp Data Fig. 4: Cross-species gut UMAP. (a)** Uniform manifold approximation and projection (UMAP) of *R. aegyptiacus,* mouse and human integrated gut scRNA-seq data colored by mutual cell subsets **(a),** and by species **(b). (c)** As in (a), but cells are separated into three panels by species),

a

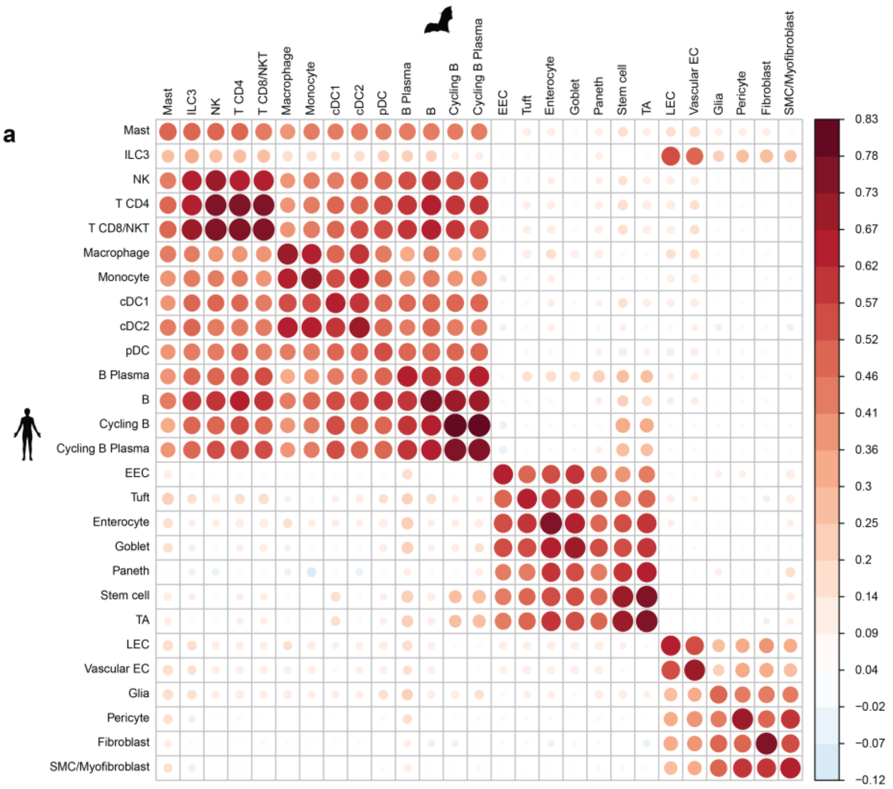
Mast
ILC3
NK
T CD4
T CD8/NKT
Macrophage
Monocyte
cDC1
cDC2
pDC
B Plasma
B
Cycling B
Cycling B Plasma
EEC
Tuft
Enterocyte
Goblet
Paneth
Stem cell
TA
LEC
Vascular EC
Glia
Pericyte
Fibroblast
SMC/Myofibroblast
0.83
0.78
0.73
0.67
0.62
0.57
0.52
0.46
0.41
0.36
0.3
0.25
0.2
0.14
0.09
0.04
−0.02
−0.07
−0.12

b

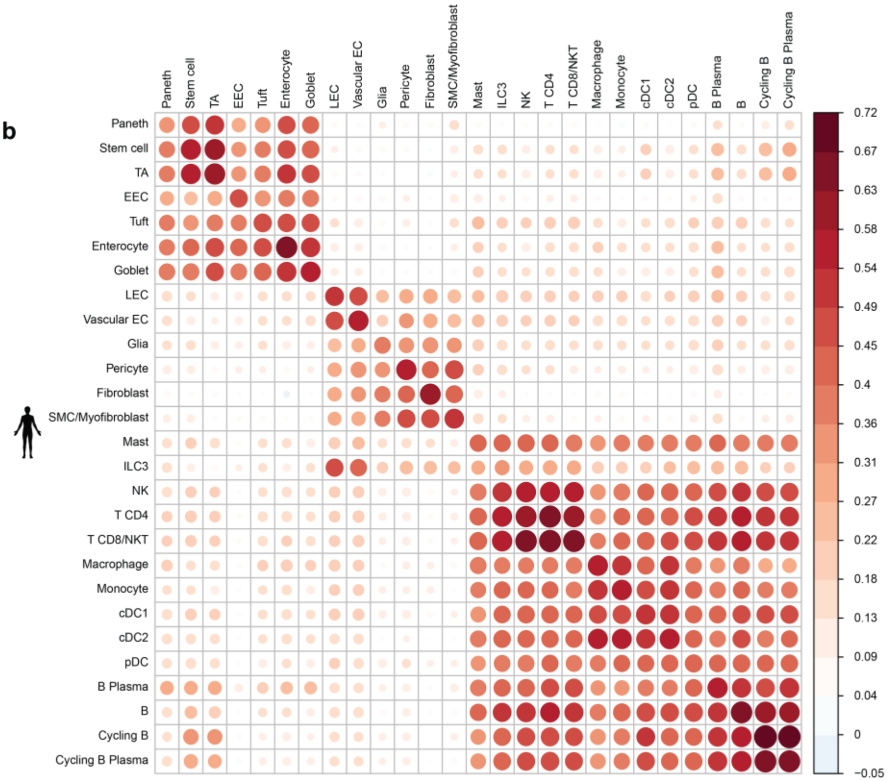
Paneth
Stem cell
TA
EEC
Tuft
Enterocyte
Goblet
LEC
Vascular EC
Glia
Pericyte
Fibroblast
SMC/Myofibroblast
Mast
ILC3
NK
T CD4
T CD8/NKT
Macrophage
Monocyte
cDC1
cDC2
pDC
B Plasma
B
Cycling B
Cycling B Plasma
0.72
0.67
0.63
0.58
0.54
0.49
0.45
0.4
0.36
0.31
0.27
0.22
0.18
0.13
0.09
0.04
0
−0.05

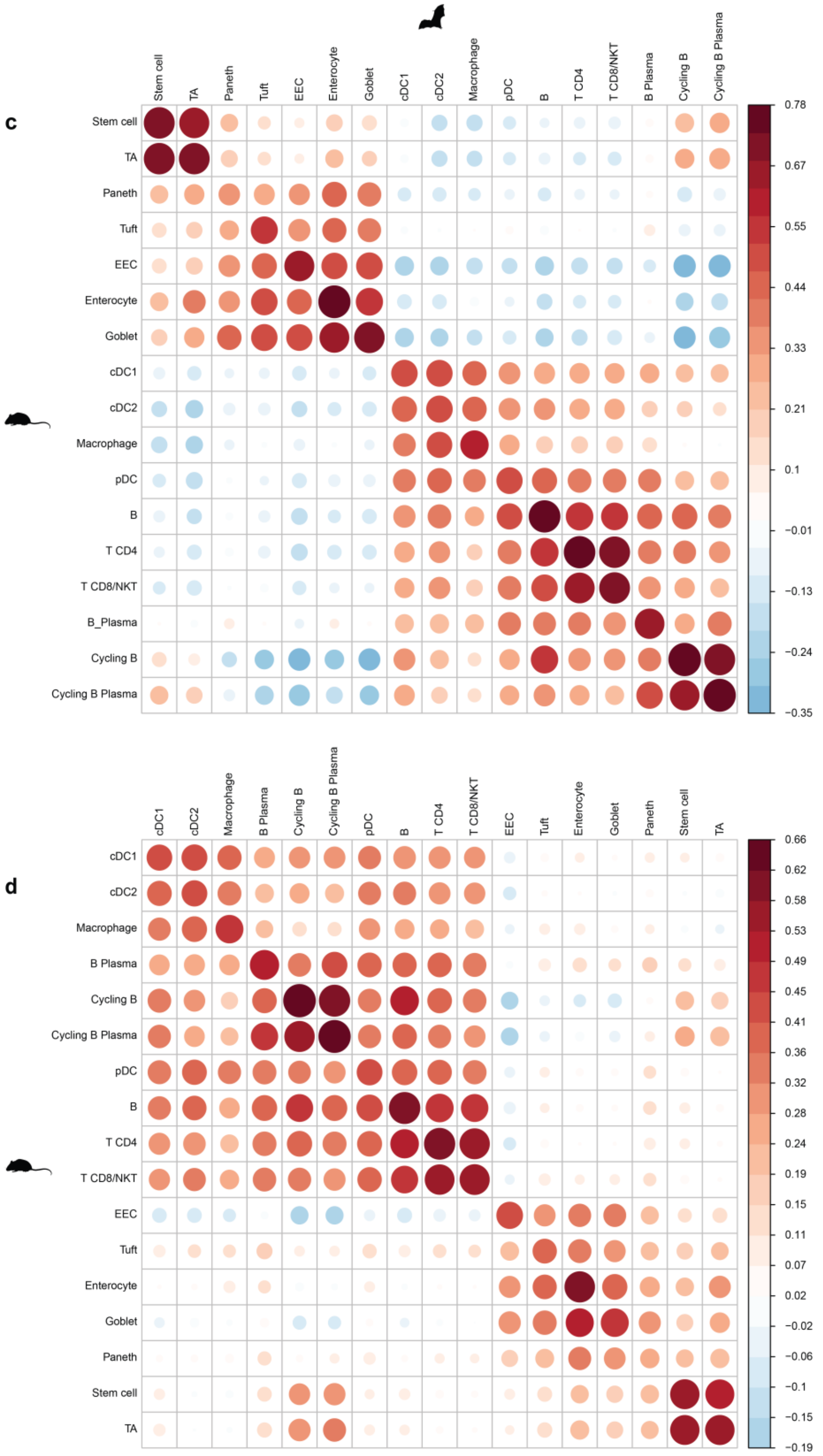


**Supp Data Fig. 5: Spearman's correlation - Lungs.** Intersection **(a, c)** and union **(b, d)** of marker genes between human-*R. aegyptiacus* **(a-b)** and mouse-*R. aegyptiacus* **(c-d)**.

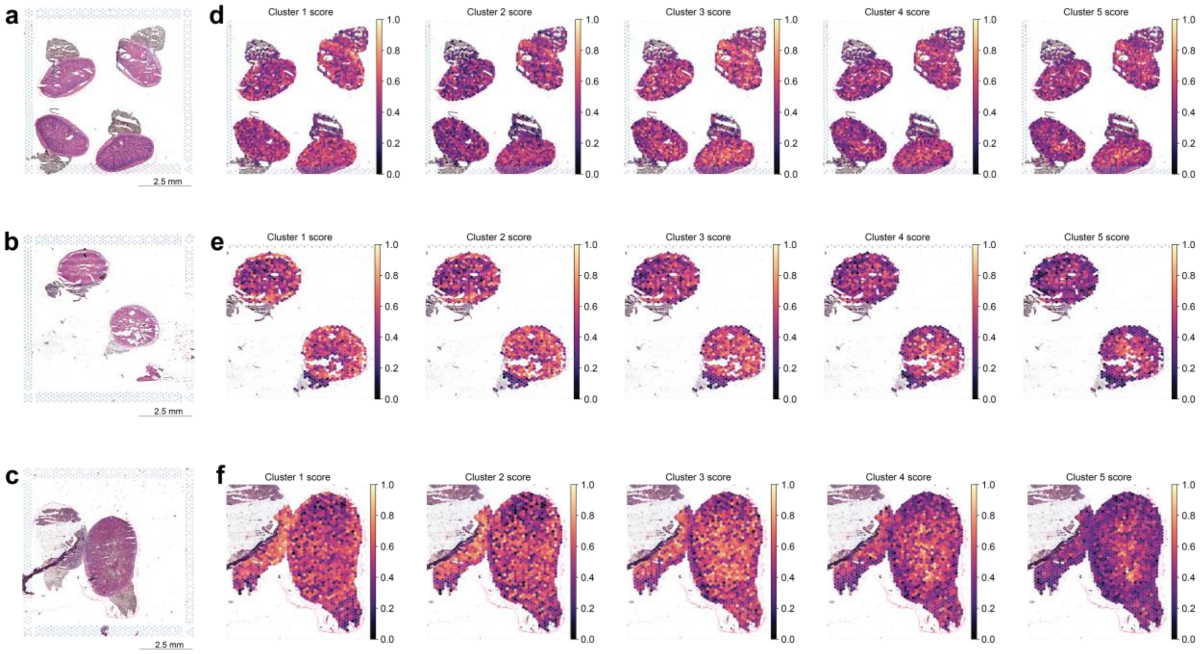


**Supp Data Fig. 6: Hematoxylin and eosin (H&E) staining of transverse cross-section of *R. aegyptiacus* gut and scores of landmark gene expression. (a)** Same individual as in Figures 1-2, **(b-c) –** other individual. **(d-f)** Scores of landmark gene expression (using mouse-to-*R. aegyptiacus* orthologs) from each of the bottom- top regions (v1-v5) overlayed on a 10x Genomics Visium data of a *R. aegyptiacus* gut cross- section.

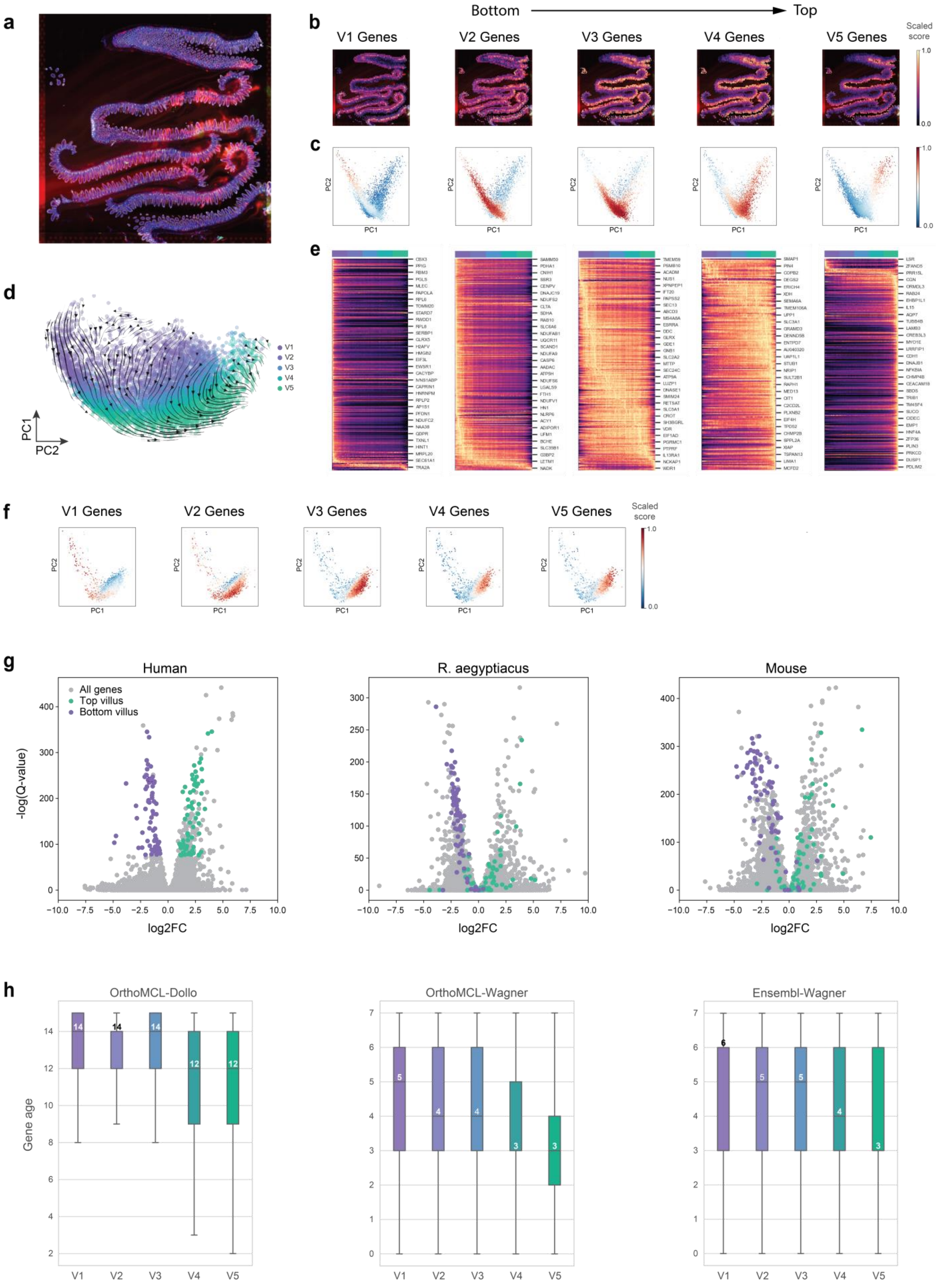

a
b
Bottom
Top
V1 Genes
V2 Genes
V3 Genes
V4 Genes
V5 Genes
Scaled score
1.0
0.0
c
PC2
PC1
d
V1
V2
V3
V4
V5
PC1
PC2
e
f
V1 Genes
V2 Genes
V3 Genes
V4 Genes
V5 Genes
Scaled score
g
Human
R. aegyptiacus
Mouse
All genes
Top villus
Bottom villus
-log(Q-value)
log2FC
h
OrthoMCL-Dollo
OrthoMCL-Wagner
Ensembl-Wagner
Gene age

**Supp Data Fig. 7: Zonation marker genes analogous analysis on mouse and human and functional absorption and transport along the villus. (a)** Hematoxylin and eosin (H&E) staining of longitudinal cross-section of mouse small intestines**. (b)** Scores of landmark gene expression (using mouse-to-*R. aegyptiacus* orthologs) from each of the bottom-top regions (v1-v5) overlayed on a 10x Genomics Visium data of a mouse small intestines cross-section. **(c, f)** Principal component analysis (PCAs) of mouse **(c)** and human **(f)** enterocytes (based on single-cell data) overlayed with scores of bottom-to-top regions (v1-v5) (red and blue mark high and low expression, respectively). **(d)** PCA of mouse enterocytes colored by their inferred spatial localization along the tissues (v1-v5) with dashed lines and arrows depicting inferred transcriptional trajectories as determined by scVelo. **(e)** Heatmaps showing landmark gene expression in mouse enterocytes, each panel including a different set of regional genes (v1-v5 regions, genes are based on mouse data[57]). Cells are ordered based on PC1 axis from 2c that denotes bottom-to-top villus localization. **(g)** Volcano plots showing DE analysis between top and bottom enterocytes in human, *R. aegyptiacus* and mouse. Human top genes are in green and human bottom genes are in purple, in the other two species we show their ortholog gene DE values, all other genes in grey. Sets of top and bottom genes are matched by q-value in human. Outliers genes that have high fold change values but not significant Q-values were omitted for visualization (these are not included in the top and bottom genes as defined above). **(h)** Gene evolutionary age analysis of landmarks genes of 5 regions (based on human genes and their age estimation using three different approaches, and taken from proteinHistorian[58]). Numbers indicate the median values for clarity.

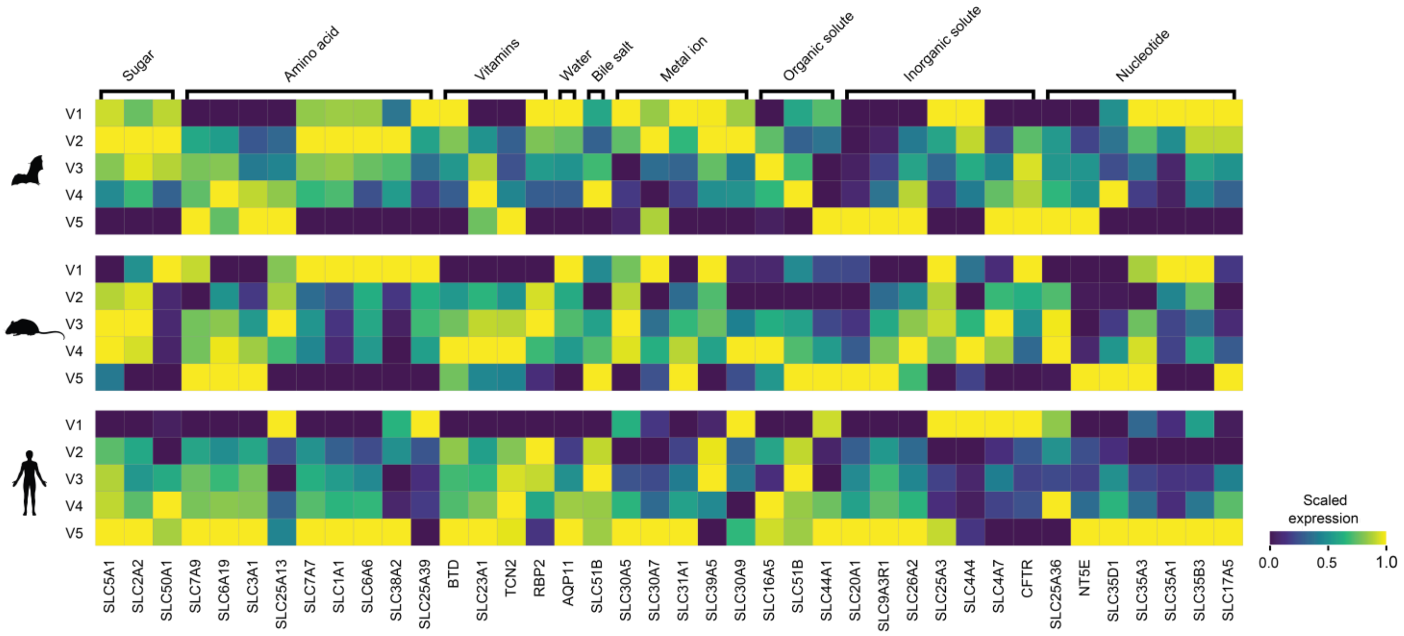


**Supp Data Fig. 8.** Heatmaps of gene expression across the 5 bottom-to-top villus regions (v1-v5), of genes involved in nutrient absorption and transport.

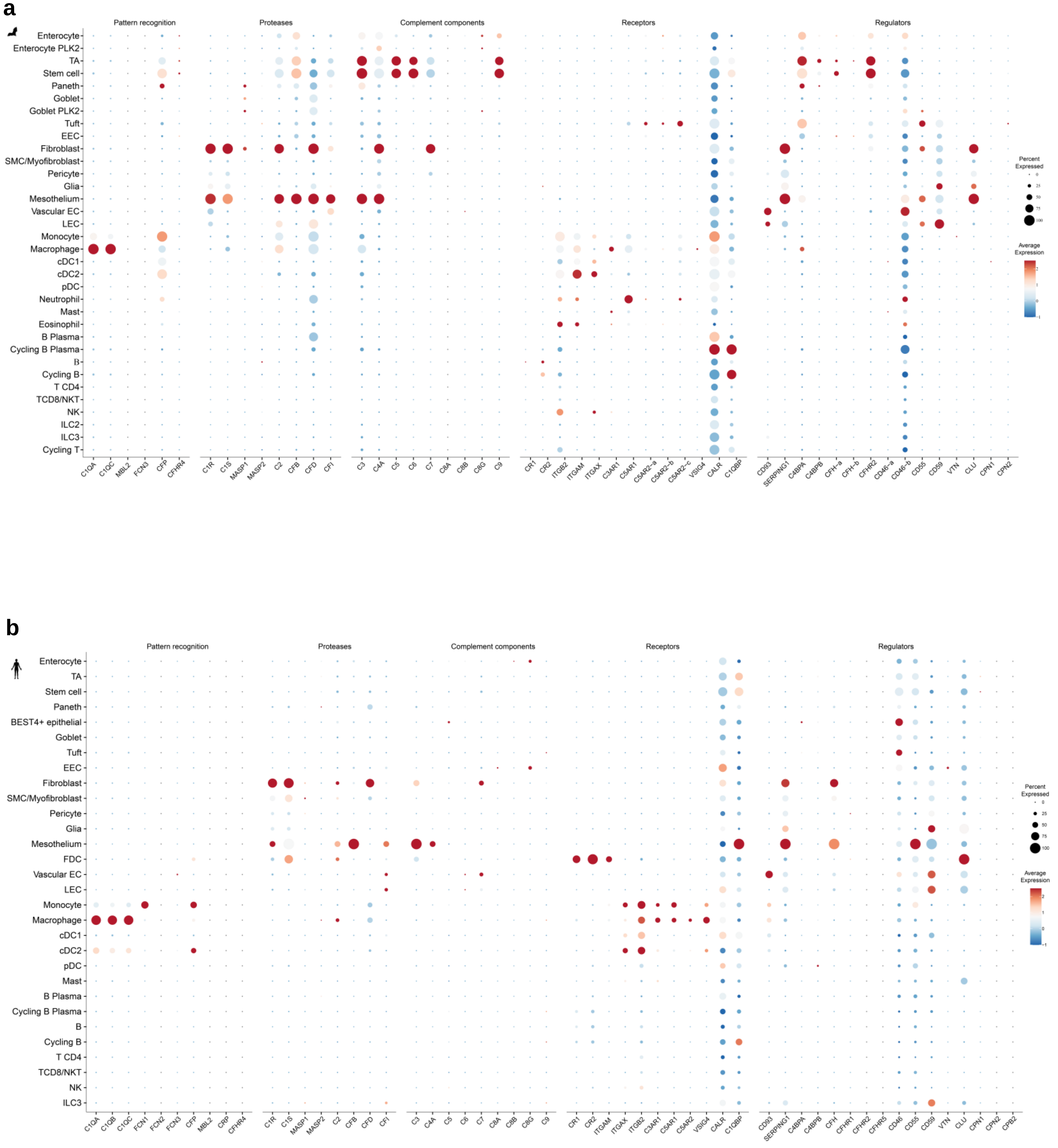
a
Pattern recognition
Proteases
Complement components
Receptors
Regulators
Enterocyte
Enterocyte PLK2
TA
Stem cell
Paneth
Goblet
Goblet PLK2
Tuft
EEC
Fibroblast
SMC/Myofibroblast
Pericyte
Glia
Mesothelium
Vascular EC
LEC
Monocyte
Macrophage
cDC1
cDC2
pDC
Neutrophil
Mast
Eosinophil
B Plasma
Cycling B Plasma
B
Cycling B
T CD4
TCD8/NKT
NK
ILC2
ILC3
Cycling T
C1QA C1QC MBL2 FCN3 CFP CFHR4
C1R C1S MASP1 MASP2 C2 CFB CFD CFI
C3 C4A C5 C6 C7 C8A C8B C8G C9
CR1 CR2 ITGB2 ITGAM ITGAX C3AR1 C5AR1 C5AR2-a C5AR2-b C5AR2-c VSIG4 CALR C1QBP
CD93 SERPING1 C4BPA C4BPB CFH-a CFH-b CFHR2 CD46-a CD46-b CD55 CD59 VTN CLU CPN1 CPN2
Percent Expressed
Average Expression
b
Pattern recognition
Proteases
Complement components
Receptors
Regulators
Enterocyte
TA
Stem cell
Paneth
BEST4+ epithelial
Goblet
Tuft
EEC
Fibroblast
SMC/Myofibroblast
Pericyte
Glia
Mesothelium
FDC
Vascular EC
LEC
Monocyte
Macrophage
cDC1
cDC2
pDC
Mast
B Plasma
Cycling B Plasma
B
Cycling B
T CD4
TCD8/NKT
NK
ILC3
C1QA C1QB C1QC FCN1 FCN2 FCN3 CFP MBL2 CRP CFHR4
C1R C1S MASP1 MASP2 C2 CFB CFD CFI
C3 C4A C5 C6 C7 C8A C8B C8G C9
CR1 CR2 ITGAM ITGAX ITGB2 C3AR1 C5AR1 C5AR2 VSIG4 CALR C1QBP
CD93 SERPING1 C4BPA C4BPB CFH CFHR1 CFHR2 CFHR5 CD46 CD55 CD59 VTN CLU CPN1 CPN2 CPB2
Percent Expressed
25
50
75
100
Average Expression

**c**

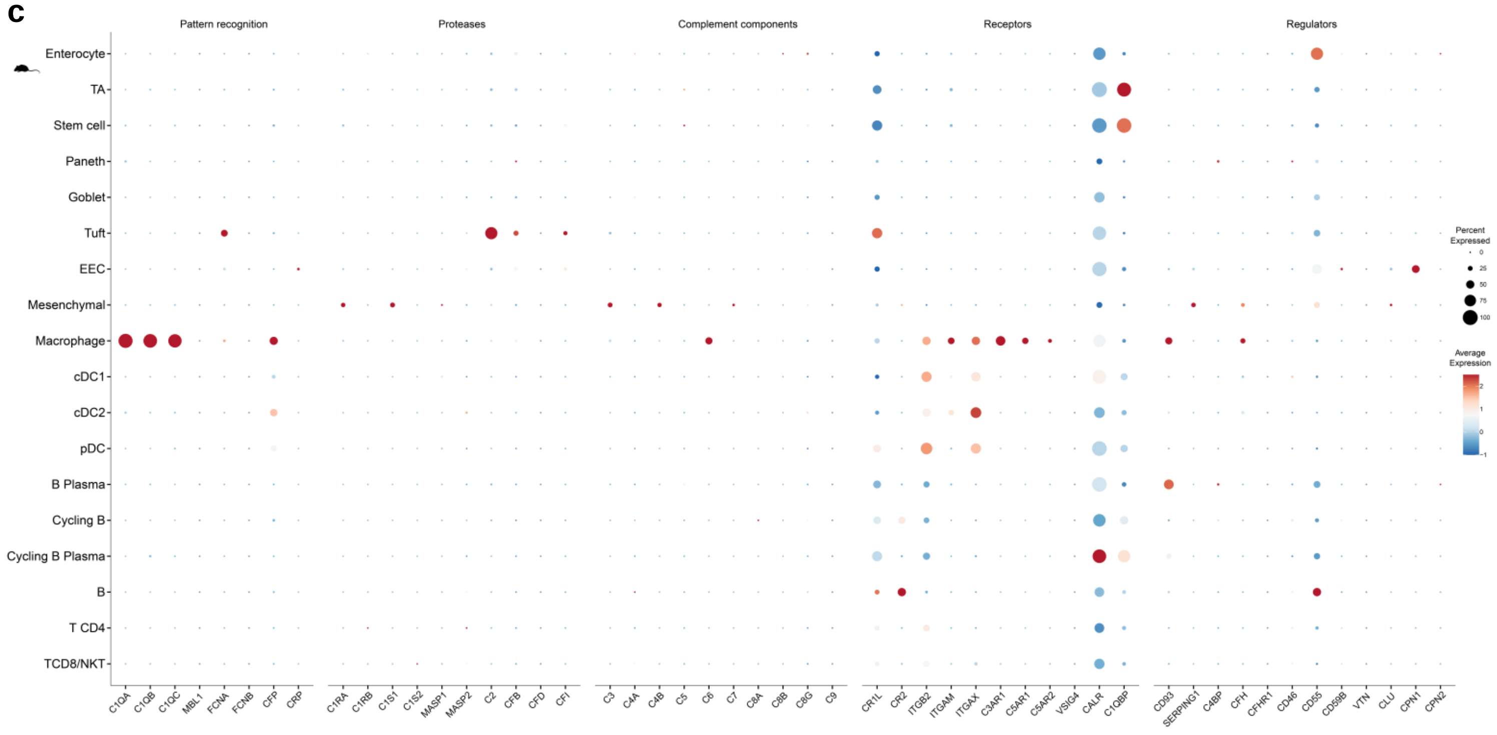


**Supp Data Fig. 9.** Dot plot illustrating the expression of all complement genes in all cell subsets identified in *R. aegyptiacus* **(a)**, human **(b)** and mouse **(c)** gut.

**a**

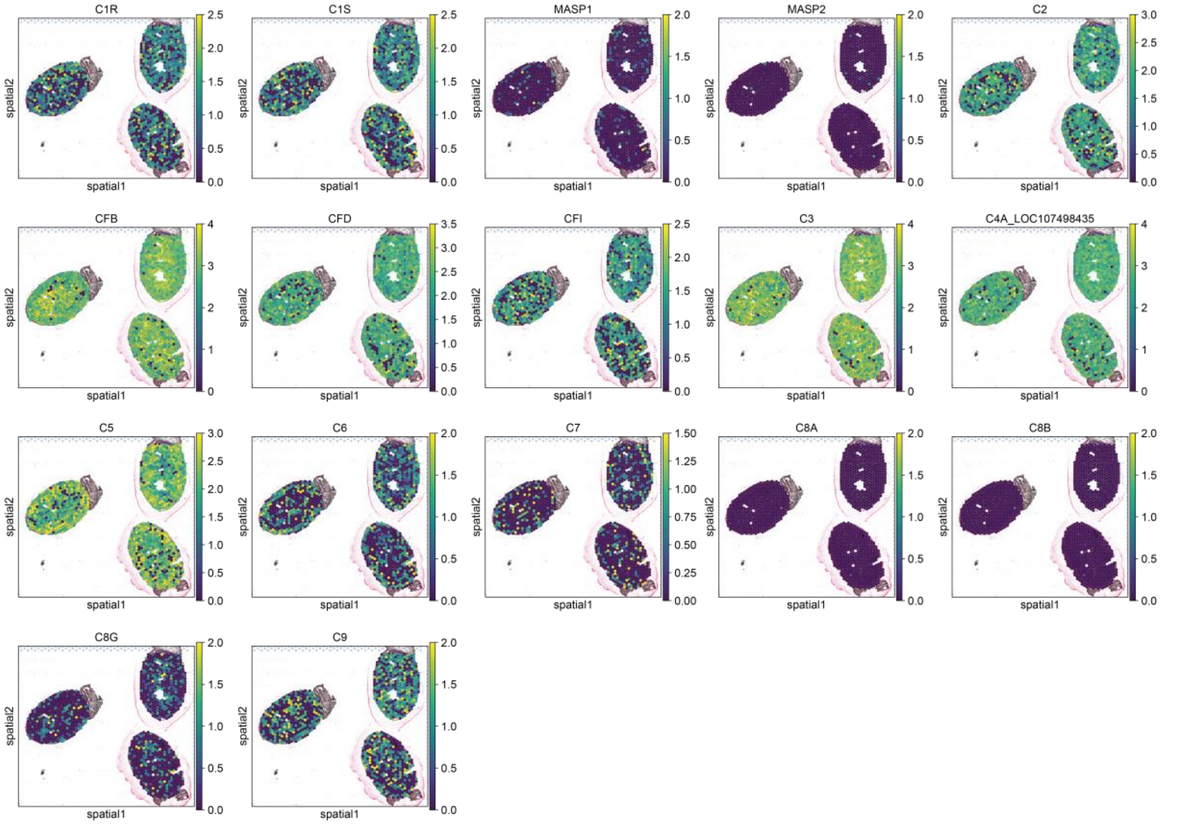


**b**

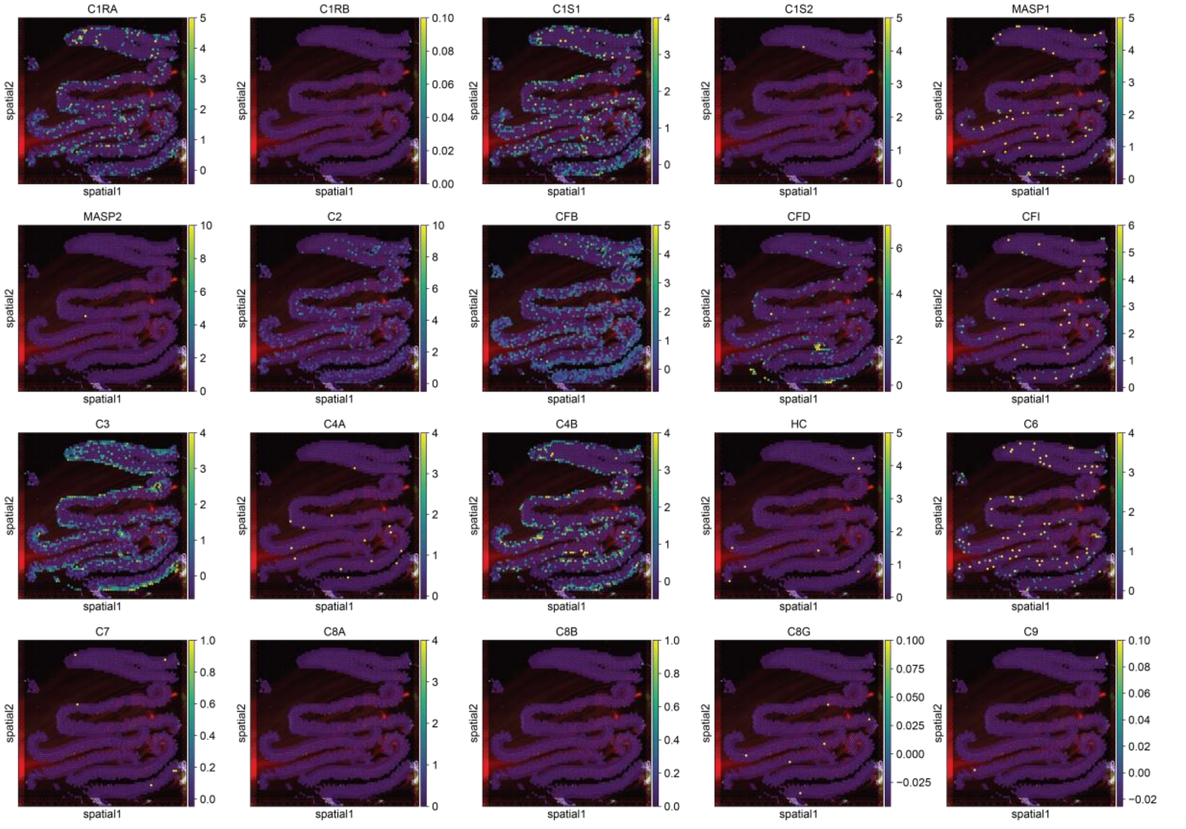


**Supp Data Fig. 10.** Expression levels of complement components and proteases genes across sections of *R. aegyptiacus* gut **(a)** and mouse small intestines **(b)**

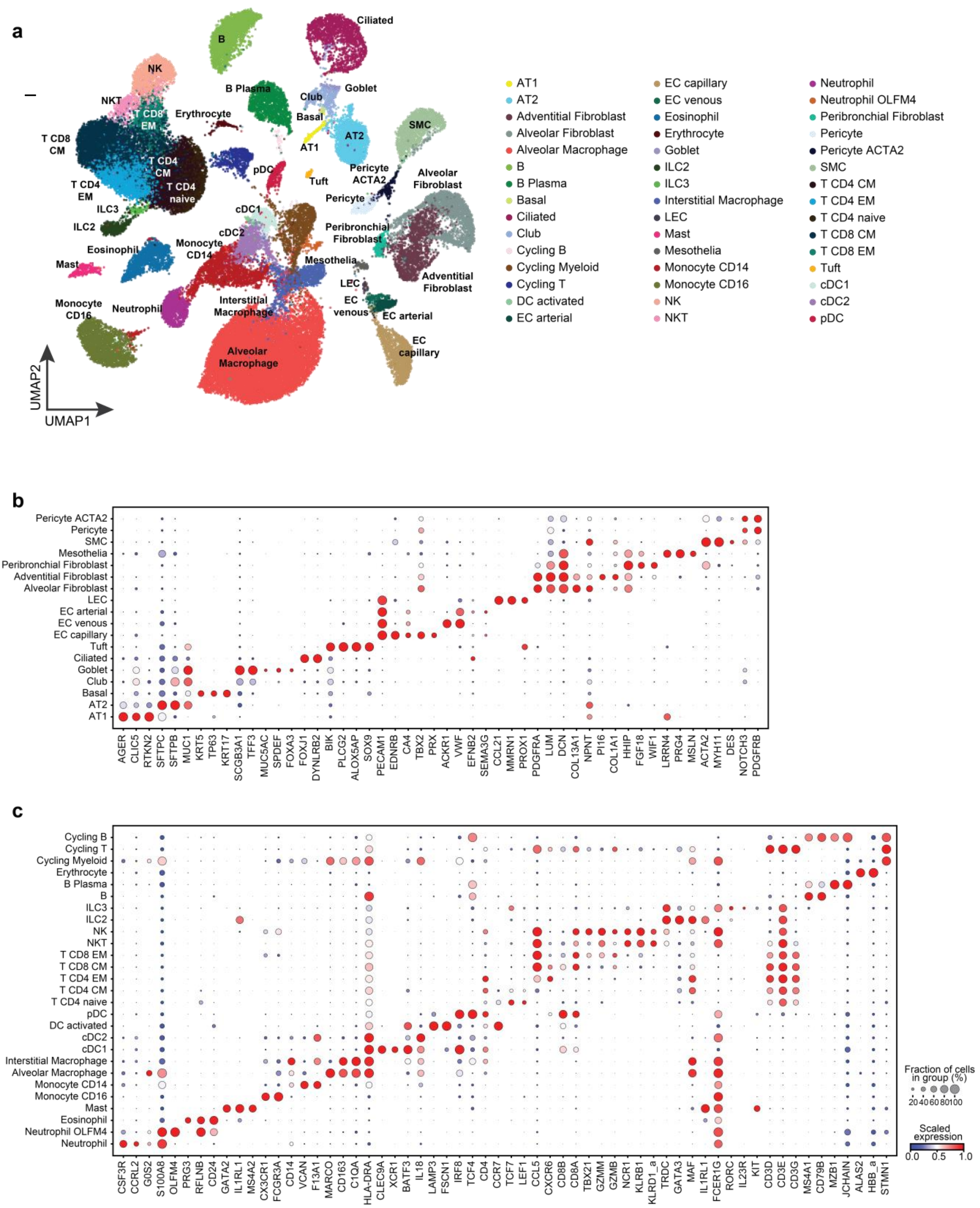


**Supp Data Fig. 11: *R. aegyptiacus* lung cell subsets defined the study. (a)** Uniform manifold approximation and projection (UMAP) of *R. aegyptiacus* lung cells clusters. Dot plots for expression of marker genes of **(b)** non- immune and **(c)** immune cell types and states in the scRNA-seq dataset.

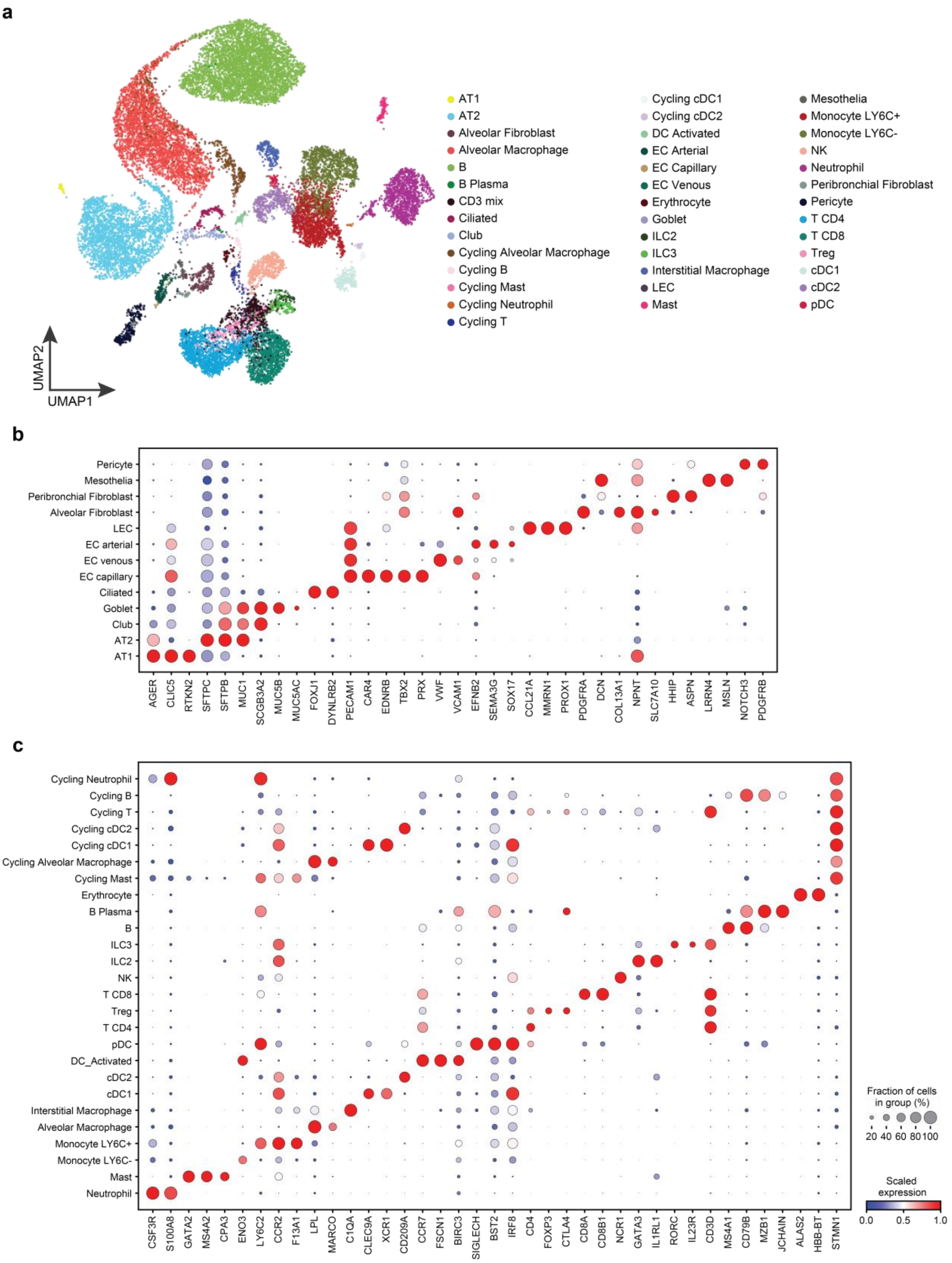


**Supp Data Fig. 12: Mouse lung characterization and cell subsets defined the study. (a)** Uniform manifold approximation and projection (UMAP) of mouse small intestinal cell clusters (3 samples, n= 21,232 cells). **(b-c)** Dot plots for expression of marker genes of **(b)** non- immune and **(c)** immune cell types and states in the scRNA-seq dataset.

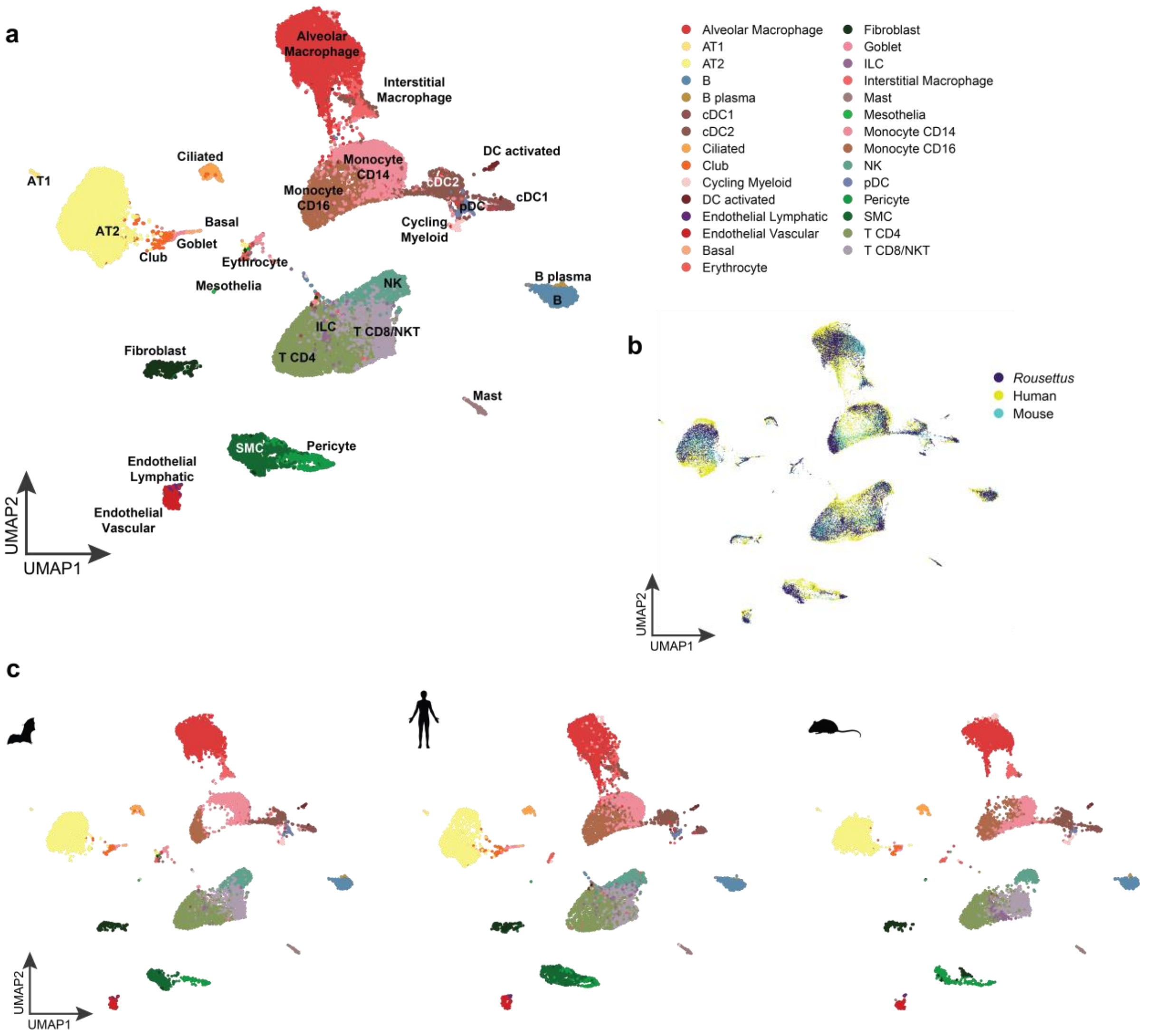


**Supp Data Fig. 13: Cross-species lung UMAP. (a)** Uniform manifold approximation and projection (UMAP) of *R. aegyptiacus,* mouse and human integrated lung scRNA-seq data colored by mutual cell subsets **(a, c**- Separate individuals), and by species **(b). (c)** As in (a) but separate panels for each species.

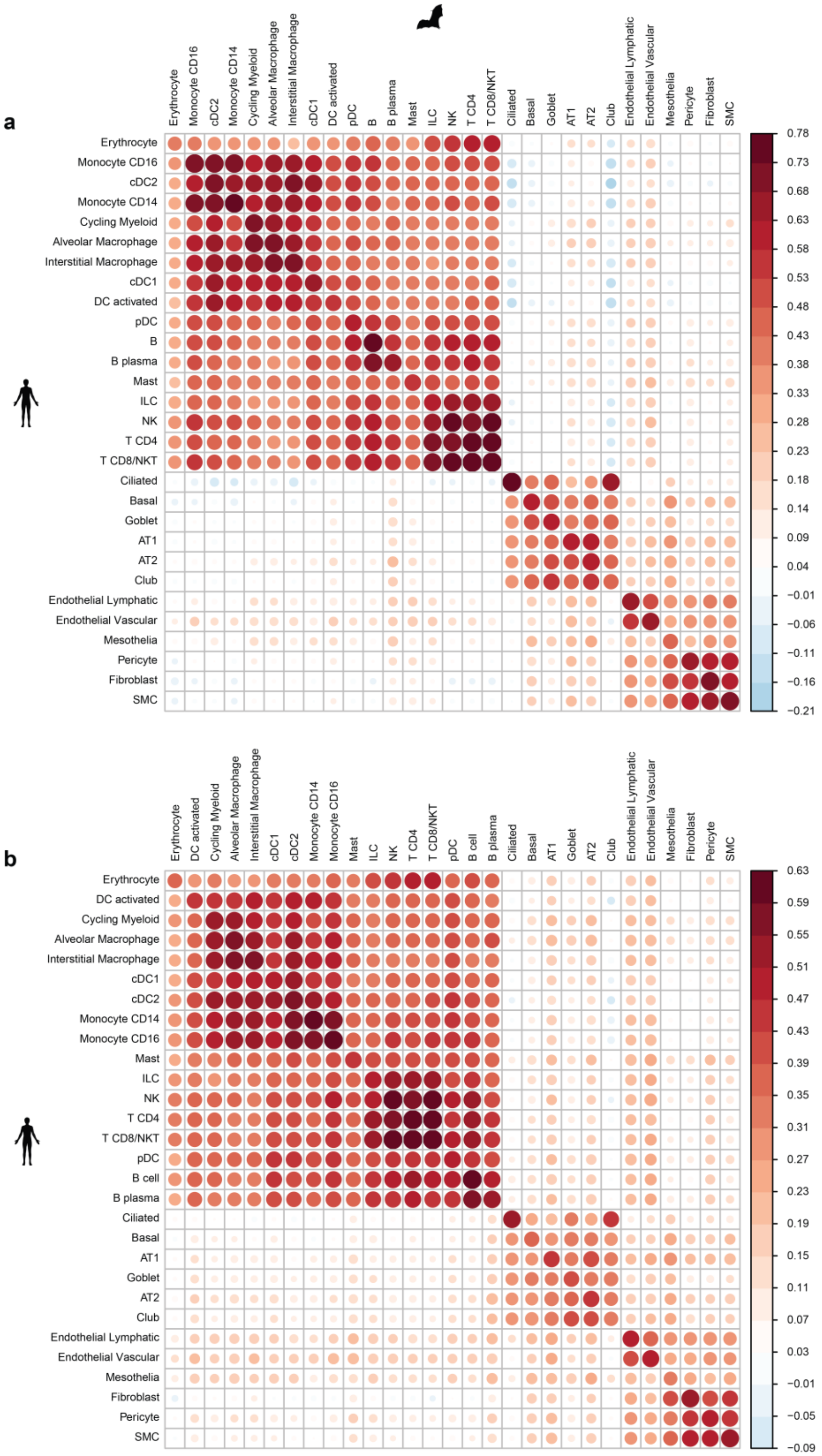
a
Erythrocyte
Monocyte CD16
cDC2
Monocyte CD14
Cycling Myeloid
Alveolar Macrophage
Interstitial Macrophage
cDC1
DC activated
pDC
B
B plasma
Mast
ILC
NK
T CD4
T CD8/NKT
Ciliated
Basal
Goblet
AT1
AT2
Club
Endothelial Lymphatic
Endothelial Vascular
Mesothelia
Pericyte
Fibroblast
SMC
0.78
0.73
0.68
0.63
0.58
0.53
0.48
0.43
0.38
0.33
0.28
0.23
0.18
0.14
0.09
0.04
−0.01
−0.06
−0.11
−0.16
−0.21
b
Erythrocyte
DC activated
Cycling Myeloid
Alveolar Macrophage
Interstitial Macrophage
cDC1
cDC2
Monocyte CD14
Monocyte CD16
Mast
ILC
NK
T CD4
T CD8/NKT
pDC
B cell
B plasma
Ciliated
Basal
AT1
Goblet
AT2
Club
Endothelial Lymphatic
Endothelial Vascular
Mesothelia
Fibroblast
Pericyte
SMC
0.63
0.59
0.55
0.51
0.47
0.43
0.39
0.35
0.31
0.27
0.23
0.19
0.15
0.11
0.07
0.03
−0.01
−0.05
−0.09

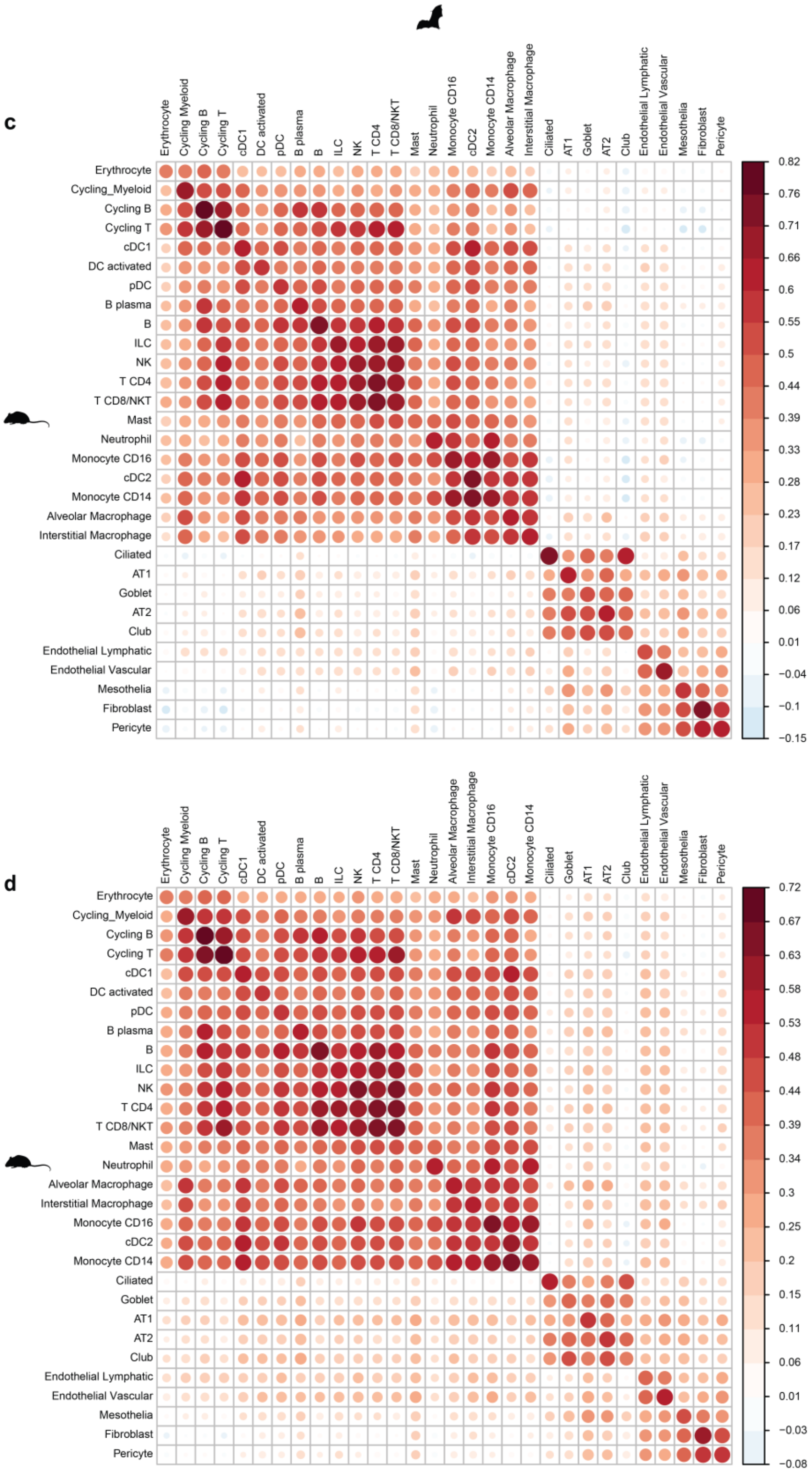


**Supp Data Fig. 14: Spearman's correlation - Gut.** Intersection **(a, c)** and union **(b, d)** of marker genes between human-*R. aegyptiacus* **(a-b)** and mouse-*R. aegyptiacus* **(c-d)**.

a

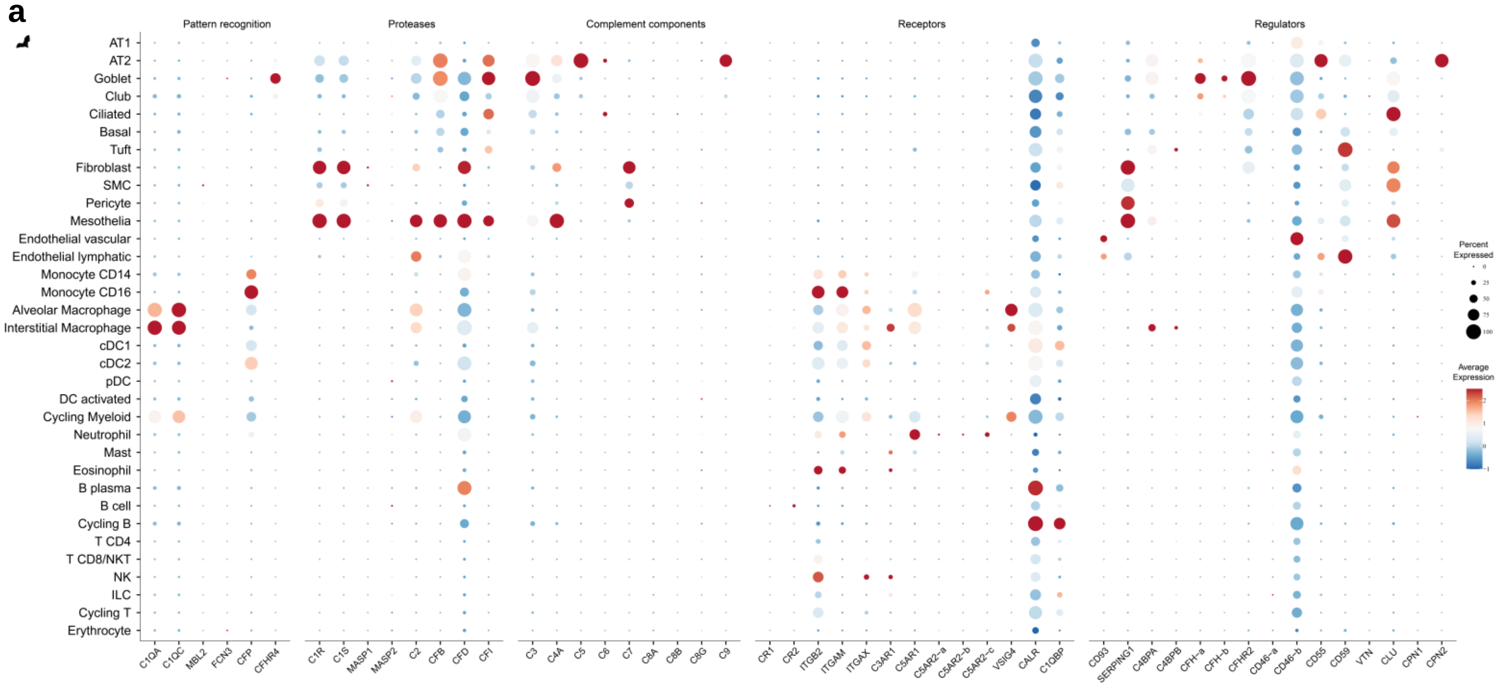
Pattern recognition
Proteases
Complement components
Receptors
Regulators
AT1
AT2
Goblet
Club
Ciliated
Basal
Tuft
Fibroblast
SMC
Pericyte
Mesothelia
Endothelial vascular
Endothelial lymphatic
Monocyte CD14
Monocyte CD16
Alveolar Macrophage
Interstitial Macrophage
cDC1
cDC2
pDC
DC activated
Cycling Myeloid
Neutrophil
Mast
Eosinophil
B plasma
B cell
Cycling B
T CD4
T CD8/NKT
NK
ILC
Cycling T
Erythrocyte
C1QA
C1QC
MBL2
FCN3
CFP
CFHR4
C1R
C1S
MASP1
MASP2
C2
CFB
CFD
CFI
C3
C4A
C5
C6
C7
C8A
C8B
C8G
C9
CR1
CR2
ITGB2
ITGAM
ITGAX
C3AR1
C5AR1
C5AR2-a
C5AR2-b
C5AR2-c
VSIG4
CALR
C1QBP
CD93
SERPING1
C4BPA
C4BPB
CFH-a
CFH-b
CFHR2
CD46-a
CD46-b
CD55
CD59
VTN
CLU
CPN1
CPN2
Percent Expressed
Average Expression

b

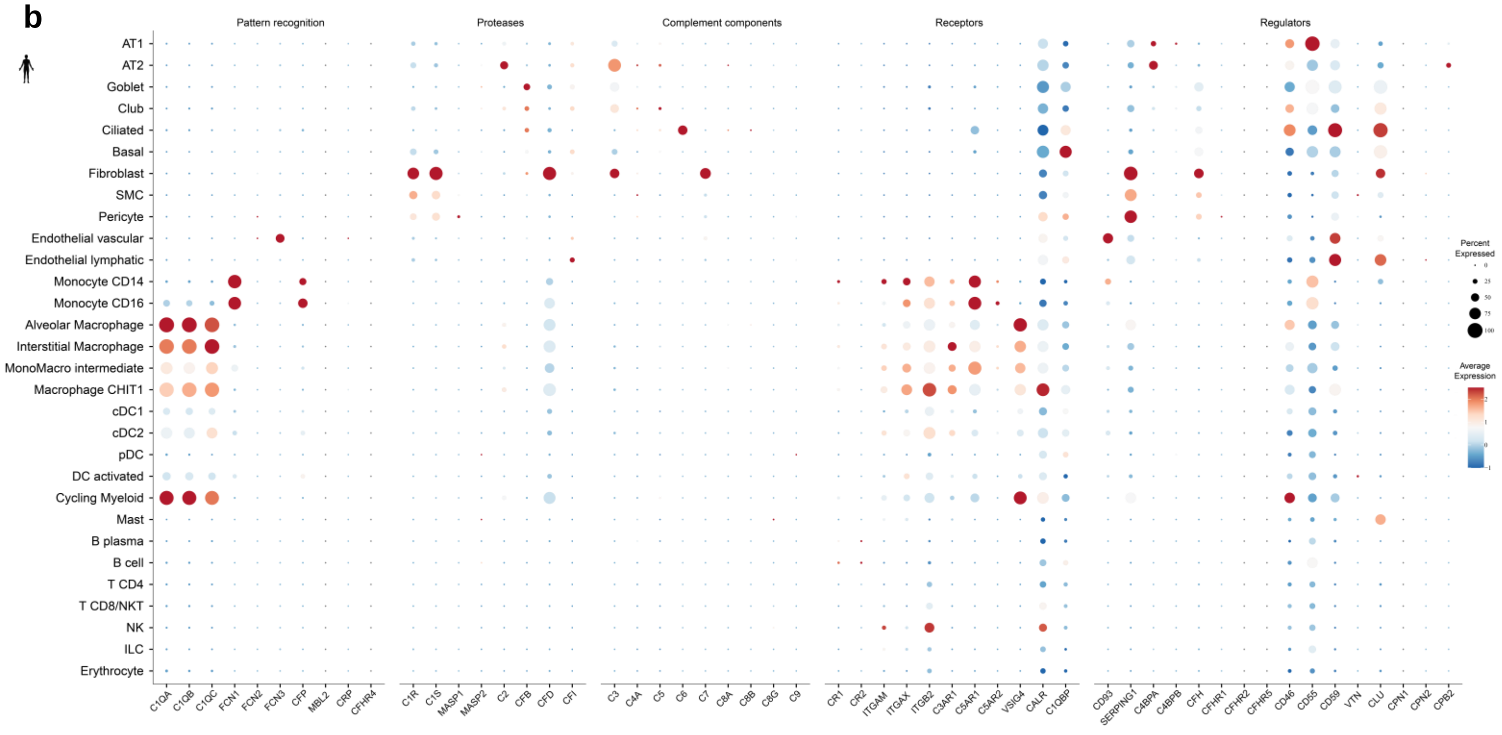
Pattern recognition
Proteases
Complement components
Receptors
Regulators
AT1
AT2
Goblet
Club
Ciliated
Basal
Fibroblast
SMC
Pericyte
Endothelial vascular
Endothelial lymphatic
Monocyte CD14
Monocyte CD16
Alveolar Macrophage
Interstitial Macrophage
MonoMacro intermediate
Macrophage CHIT1
cDC1
cDC2
pDC
DC activated
Cycling Myeloid
Mast
B plasma
B cell
T CD4
T CD8/NKT
NK
ILC
Erythrocyte
C1QA
C1QB
C1QC
FCN1
FCN2
FCN3
CFP
MBL2
CRP
CFHR4
C1R
C1S
MASP1
MASP2
C2
CFB
CFD
CFI
C3
C4A
C5
C6
C7
C8A
C8B
C8G
C9
CR1
CR2
ITGAM
ITGAX
ITGB2
C3AR1
C5AR1
C5AR2
VSIG4
CALR
C1QBP
CD93
SERPING1
C4BPA
C4BPB
CFH
CFHR1
CFHR2
CFHR5
CD46
CD55
CD59
VTN
CLU
CPN1
CPN2
CPB2
Percent Expressed
Average Expression

c

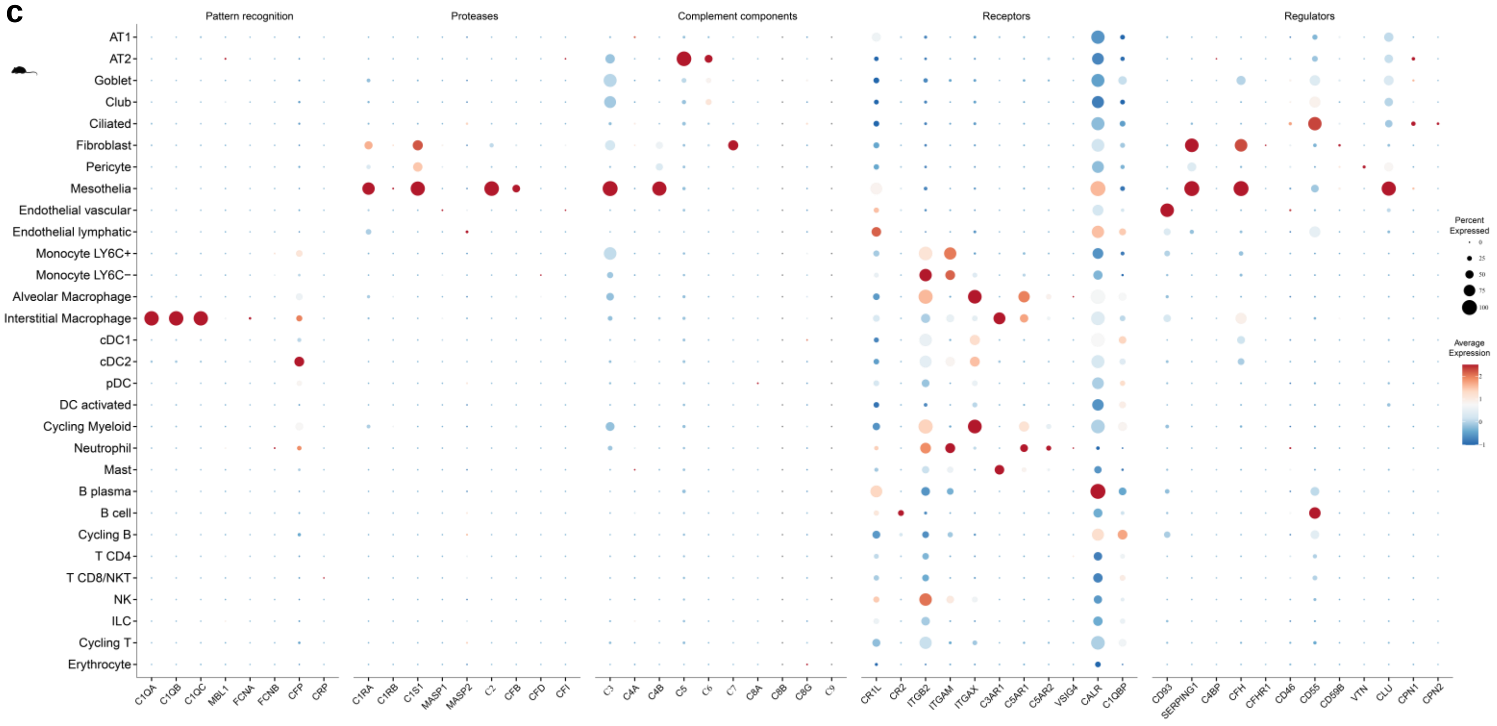


**Supp Data Fig. 15.** Dot plot illustrating the expression of all complement genes in all cell subsets identified in *R. aegyptiacus* **(a)**, human **(b)** and mouse **(c)** lungs.

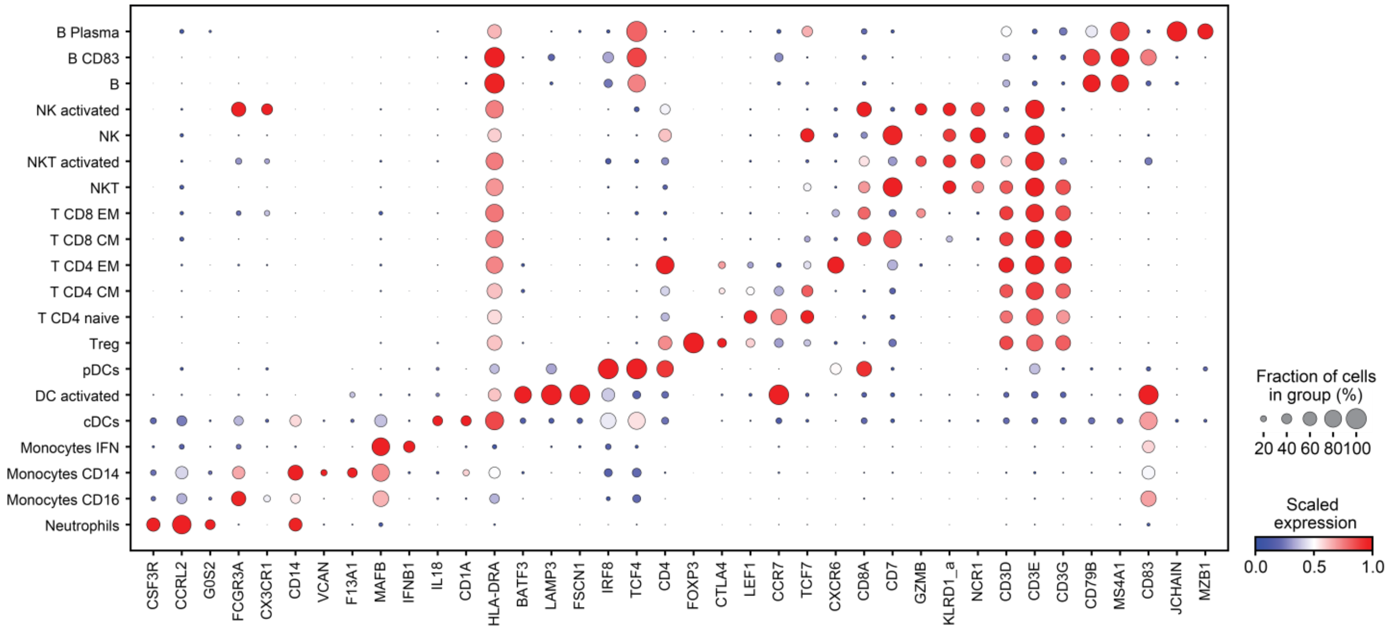


**Supp Data Fig. 16: *R. aegyptiacus* PBMCs defined the study.** Dot plot for expression of marker genes of cell types and states in the scRNA-seq dataset.

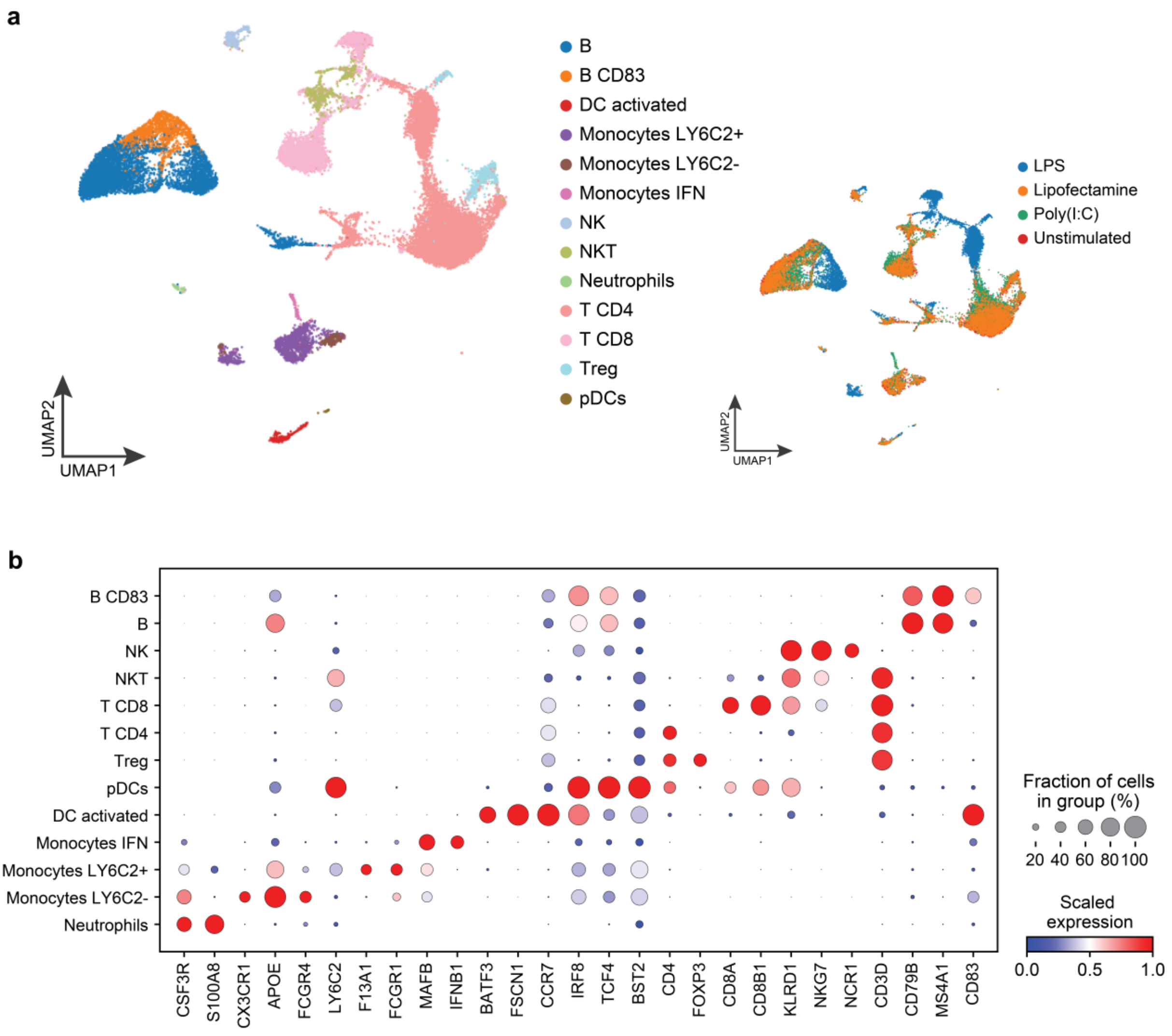


**Supp Data Fig. 17: Mouse PBMCs characterization and subsets defined the study. (a)** Uniform manifold approximation and projection (UMAP) of mouse PBMC colored by cell clusters (left) and by conditions (right) (15 samples, n= 24,240 cells). **(b)** Dot plot for expression of marker genes of cell types and states in the scRNA-seq dataset.

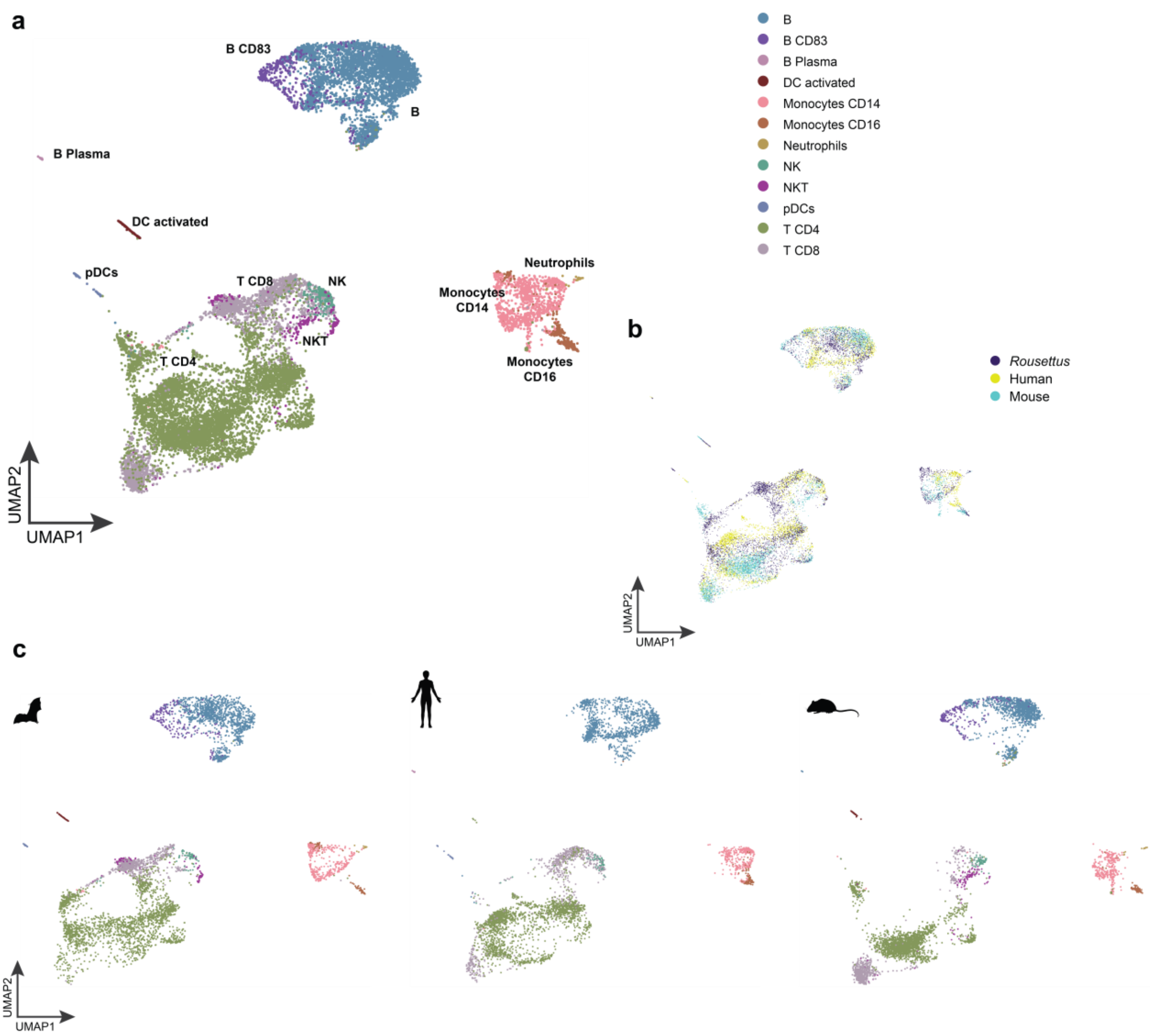


**Supp Data Fig. 18: Cross-species PBMCs UMAP. (a)** Uniform manifold approximation and projection (UMAP) of *R. aegyptiacus,* mouse and human integrated PBMCs scRNA-seq data colored by mutual cell subsets (**a**), and by species **(b). (c)** As in (a) but separate panels for each species.

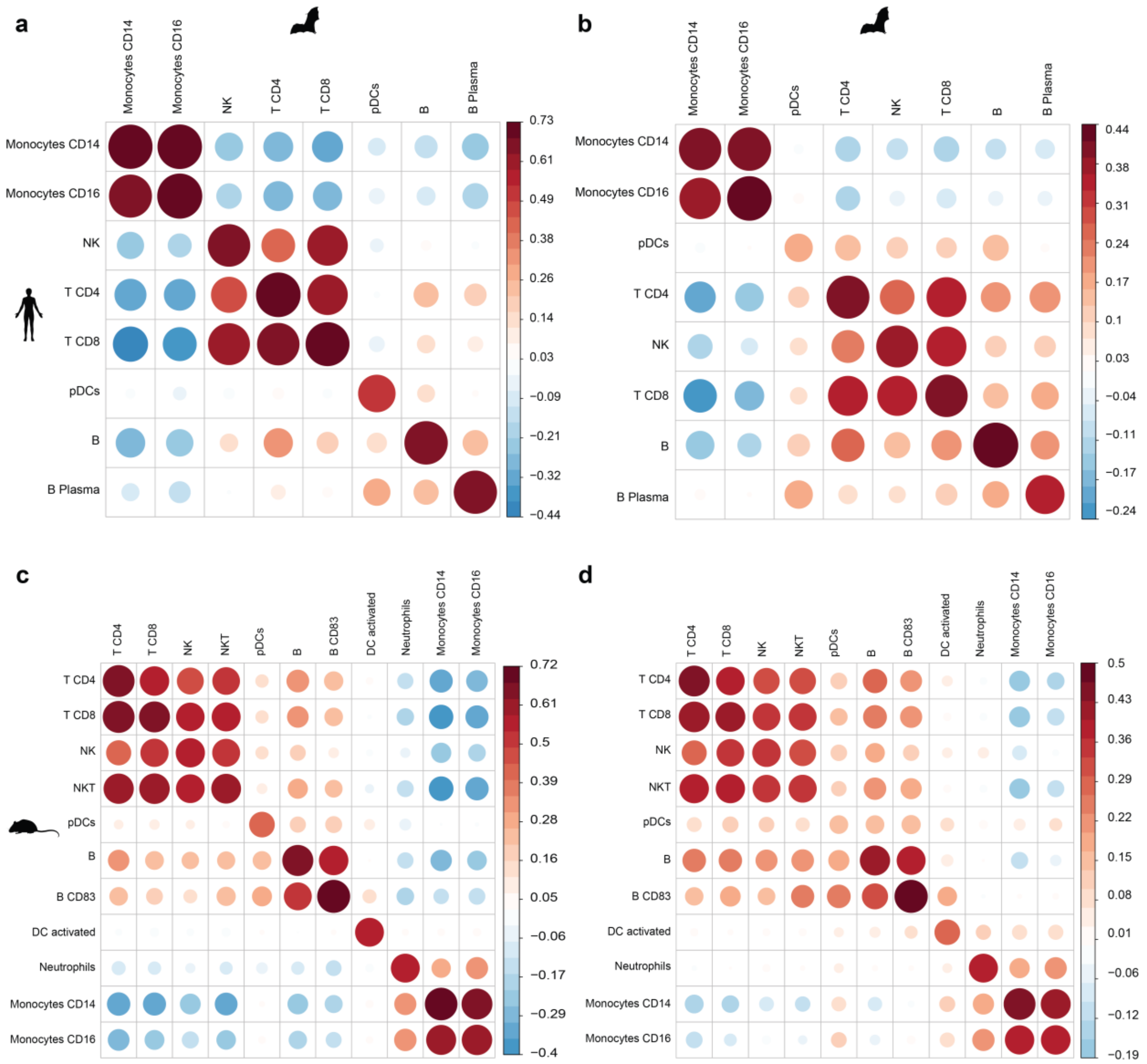


**Supp Data Fig. 19: Spearman's correlation - PBMCs.** Intersection **(a, c)** and union **(b, d)** of marker genes between human-*R. aegyptiacus* **(a-b)** and mouse-*R. aegyptiacus* **(c-d)**.

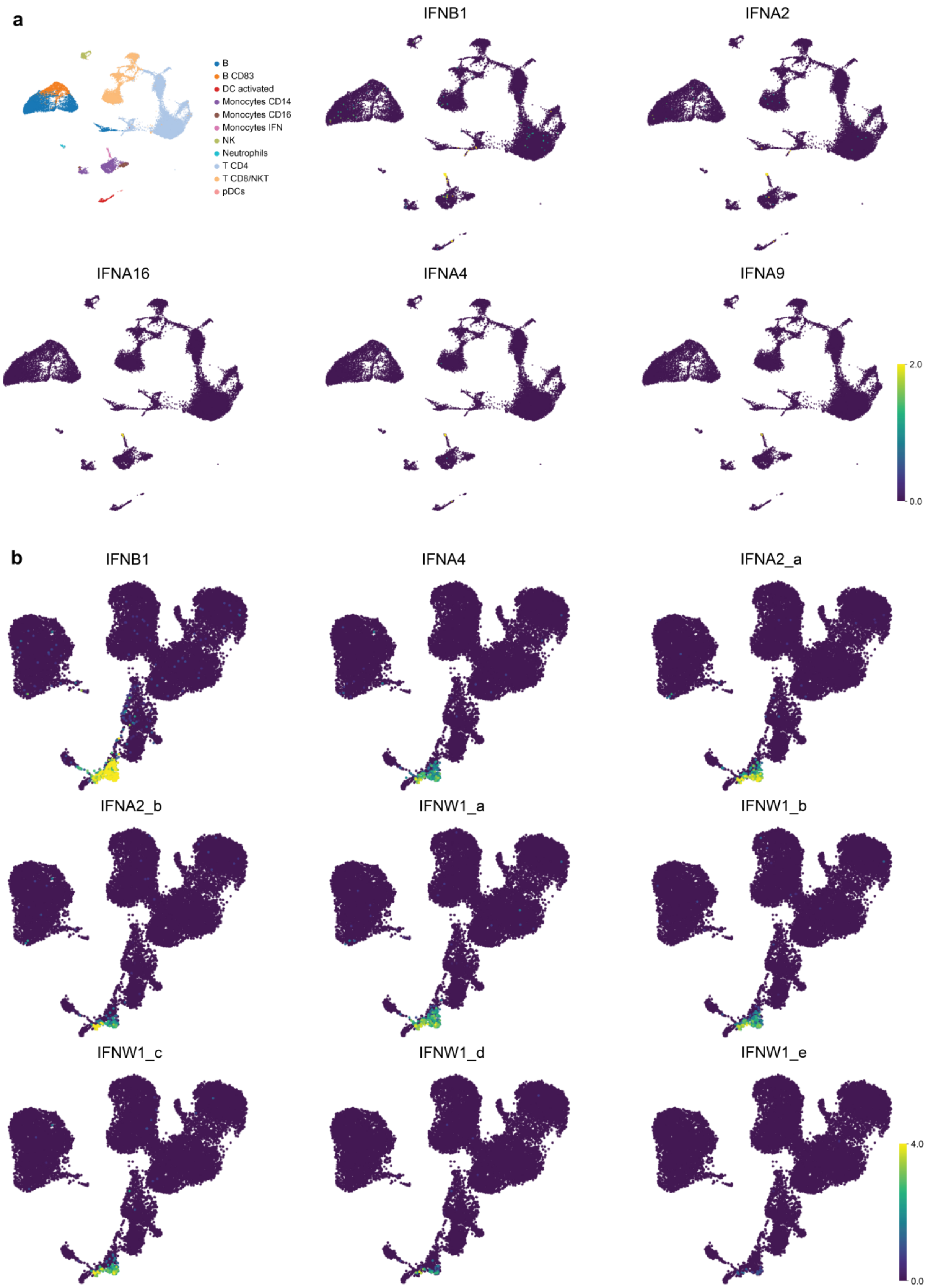


**Supp Data Fig. 20. (a)** UMAP of dsRNA-stimulated mouse PBMCs, colored by cell type (upper- left), and showing expression levels of IFNB1 and other type-I IFNs. **(b)** Additional type-I IFNs expression in *R. aegyptiacus* dsRNA-stimulated PBMCs.
